# Cryptic Patterns of Speciation in Cryptic Primates: Microendemic Mouse Lemurs and the Multispecies Coalescent

**DOI:** 10.1101/742361

**Authors:** Jelmer Poelstra, Jordi Salmona, George P. Tiley, Dominik Schüßler, Marina B. Blanco, Jean B. Andriambeloson, Sophie Manzi, C. Ryan Campbell, Olivier Bouchez, Paul D. Etter, Amaia Iribar, Paul A. Hohenlohe, Kelsie E. Hunnicutt, Eric A. Johnson, Peter M. Kappeler, Peter A. Larsen, José M. Ralison, Blanchard Randrianambinina, Rodin M. Rasoloarison, David W. Rasolofoson, Amanda R. Stahlke, David Weisrock, Rachel C. Williams, Lounès Chikhi, Edward E Louis, Ute Radespiel, Anne D. Yoder

## Abstract

Mouse lemurs (*Microcebus*) are a radiation of morphologically cryptic primates distributed throughout Madagascar for which the number of recognized species has exploded in the past two decades. This taxonomic explosion has prompted understandable concern that there has been substantial oversplitting in the mouse lemur clade. Here, we take an integrative approach to investigate species diversity in two pairs of sister lineages that occur in a region in northeastern Madagascar with high levels of microendemism and predicted habitat loss. We analyzed RADseq data with multispecies coalescent (MSC) species delimitation methods for three named species and an undescribed lineage previously identified to have divergent mtDNA. Marked differences in effective population sizes, levels of gene flow, patterns of isolation-by-distance, and species delimitation results were found among them. Whereas all tests support the recognition of the presently undescribed lineage as a separate species, the species-level distinction of two previously described species, *M. mittermeieri* and *M. lehilahytsara* is not supported – a result that is particularly striking when using the genealogical discordance index (*gdi*). Non-sister lineages occur sympatrically in two of the localities sampled for this study, despite an estimated divergence time of less than 1 Ma. This suggests rapid evolution of reproductive isolation in the focal lineages, and in the mouse lemur clade generally. The divergence time estimates reported here are based on the MSC and calibrated with pedigree-based mutation rates and are considerably more recent than previously published fossil-calibrated concatenated likelihood estimates, however. We discuss the possible explanations for this discrepancy, noting that there are theoretical justifications for preferring the MSC estimates in this case.

## Introduction

Mouse lemurs (*Microcebus* spp.) are small, nocturnal primates that are widespread in the forests of Madagascar (Mittermeier et al. 2010), one of the world’s most biodiverse environments (Myers et al. 2000; Goodman and Benstead 2005; Estrada et al. 2017). Mouse lemur diversity was long overlooked (Zimmermann and Radespiel 2014) until the introduction of genetic analyses made it feasible to identify diverging lineages despite similar phenotypes and ecological niches. This genetic perspective has led to the description of many new species, with 24 species recognized at present. In one such study, Radespiel et al. (2008) surveyed the forests of the Makira region (Fig. 1) in northeastern Madagascar and found evidence for three divergent mitochondrial lineages occurring in sympatry. One of these was identified as *M*. *mittermeieri* (Louis et al. 2006), while the second was newly described as *M*. *macarthurii*. A third lineage, provisionally called M. sp. #3, was hypothesized to represent a new species closely related to *M*. *macarthurii* but was not formally named because the data was limited to mtDNA sequence data from one individual. Furthermore, two other species occur in the region, *M*. *lehilahytsara* (Roos and Kappeler in Kappeler et al. 2005) at higher elevations, and *M*. *simmonsi* (Louis et al. 2006) in lowland forests in the south (Fig. 1).

**Figure 1:**
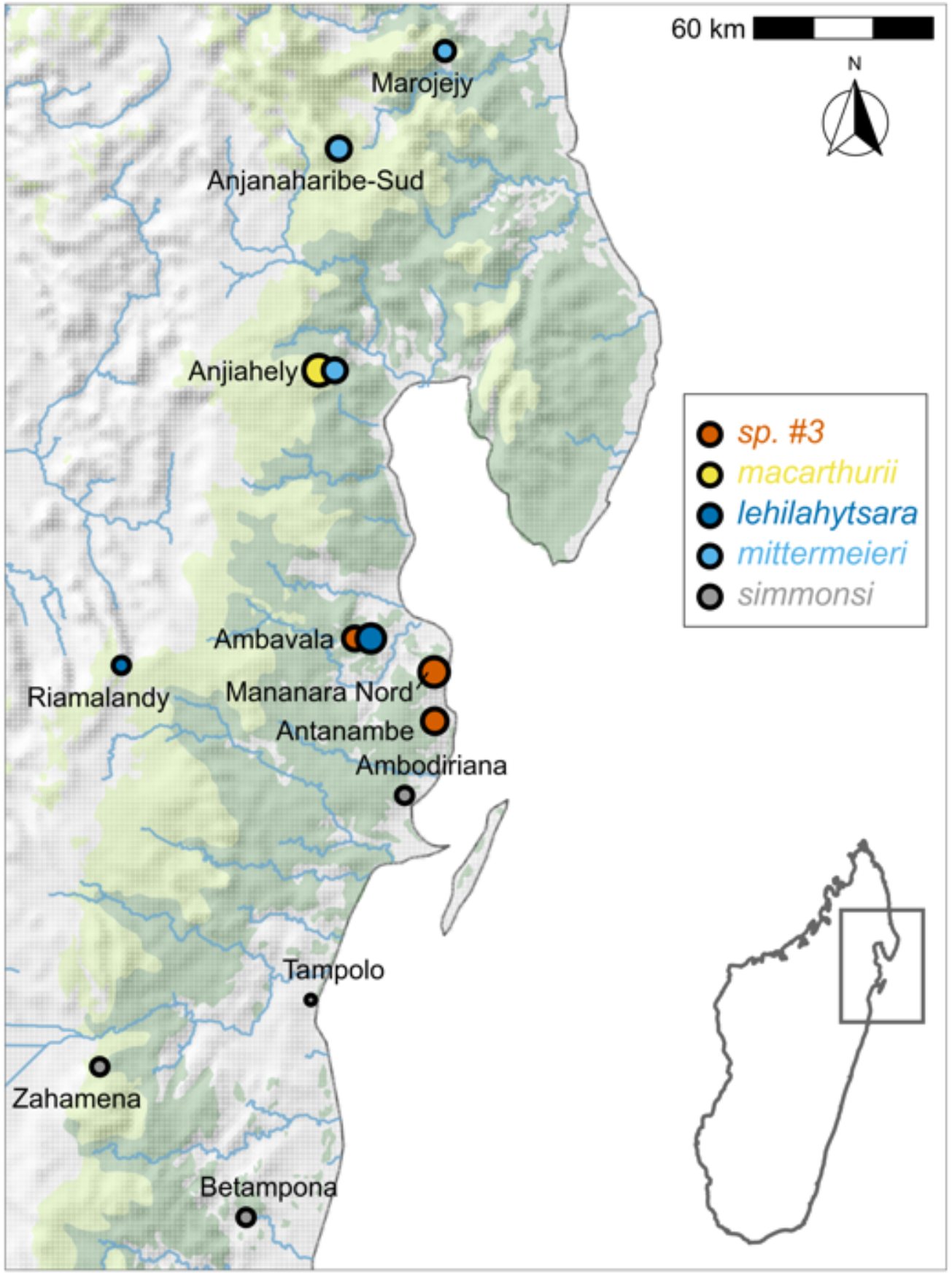
Sampling sites in northeastern Madagascar. Size of the circles scales with the number of individuals sequenced for a given site. Green background indicates forest cover as per Du Put & Moat (1998), with darker green indicating “low altitude” and paler green indicating “mid altitude” evergreen humid forest. At Anjiahely and Ambavala, two species were detected; in both cases, the leftmost site marker was slightly

Given that many previous taxonomic descriptions of mouse lemurs have relied strongly, if not entirely, on mtDNA sequence divergence, there has been criticism that mouse lemurs (and lemurs more generally) may have been oversplit (Tattersall 2007; Markolf et al. 2011). Species delimitation using only mtDNA is now widely regarded as problematic, given that the mitochondrial genome represents a single non-recombining locus whose gene tree may not represent the underlying species tree (e.g., Pamilo and Nei 1988; Maddison 1997). Mitochondria are also maternally inherited and therefore susceptible to effects of male-biased dispersal (e.g., Dávalos and Russell 2014), which is prevalent in mouse lemurs (reviewed in Radespiel 2016). Moreover, previous attempts to resolve mouse lemur relationships using nuclear sequences have been complicated by high gene tree discordance, consistent with strong incomplete lineage sorting (e.g., Heckman et al. 2007; Weisrock et al. 2010). These issues can be overcome with genomic approaches, which provide power for simultaneously resolving phylogenetic relationships and estimating demographic parameters such as divergence times, effective population sizes, and rates of gene flow — even among closely related species (e.g., Palkopoulou et al. 2018; Pedersen et al. 2018).

Given that cryptic species are by definition difficult to identify based on phenotypic characters (Bickford et al. 2007), recently developed methods for genomic species delimitation have advanced our ability to recognize and quantify their species diversity. In the past decade, both theory and methods for species delimitation have seen substantial progress, especially those which leverage the multispecies coalescent (MSC) model (Pamilo and Nei 1988; Rannala and Yang 2003). MSC-based species delimitation methods have been increasingly applied to genomic data (e.g. Carstens and Dewey 2010; Yang and Rannala 2010; Grummer et al. 2014; Dincă et al. 2019; Hundsdoerfer et al. 2019), though they have also been considered controversial (Edwards and Knowles 2014; Sukumaran and Knowles 2017; Barley et al. 2018). The controversy largely relates to the idea that strong population structure can be mistaken for species boundaries, which may lead to oversplitting (Jackson et al. 2017; Sukumaran and Knowles 2017; Luo et al. 2018; Leaché et al. 2019; Chambers and Hillis 2020). To overcome this potential weakness, Jackson et al. (2017) proposed a heuristic criterion, the genealogical divergence index (*gdi*), with Leaché et al. (2019) further suggesting that *gdi* helps to differentiate between population structure and species-level divergence. In parallel, sophisticated statistical approaches have been developed that can detect the presence and magnitude of gene flow during or after speciation (Gronau et al. 2011; Payseur and Rieseberg 2016; Dalquen et al. 2017; Wen et al. 2018). Taken together, these analytical developments are crucial to our ability to recognize the patterns that characterize the speciation process, despite the challenge of identifying species without universally agreed upon criteria (de Queiroz 2007).

In this study, we use a structured framework starting with phylogenetic placement of lineages and culminating with the MSC to delimit species, estimate divergence times, identify post-divergence gene flow, and to estimate both current and ancestral effective population sizes (`Fig. S1`). We take advantage of increased geographic, population-level, and genomic sampling to comparatively examine speciation dynamics for two pairs of closely related lineages in the region (described below as Clades I and II) and perform MSC species delimitation methods with Restriction-site Associated DNA sequencing (RADseq) data to infer divergence times, effective population sizes, and rates of gene flow between these lineages. We also provide a novel wholegenome assembly for the previously undescribed lineage and compare inferences of effective population size (*N_e_*) through time from whole-genome versus RADseq data. We find notably different species delimitation results for the lineages in the two mouse lemur clades and believe that the integrative analytic framework here used can be applied more generally to allow investigators to test hypotheses of population-versus species-level differentiation.

## Materials and Methods

### Summary of Analyses

We generated RADseq data for 63 individuals from 6 lineages, of which 48 were from the two focal clades and passed quality control. First, we used maximum likelihood approaches to infer relationships among lineages and to provide a framework for subsequent species delimitation analyses (Fig. 2A and C). To delimit species, we performed clustering (Fig. 2B) and PCA analyses (Fig. 3A-C), as well as formal MSC species delimitation analyses using SNAPP and BPP. We also used the recently developed genealogical divergence index *gdi* based on BPP parameter estimates (Fig. 3D) and performed an isolation-by-distance analysis (Fig. 4). To determine to what extent ongoing and ancestral gene flow may have contributed to current patterns of divergence, we used G-PhoCS and D-statistics (Fig. 5). Finally, we generated wholegenome sequencing data for a single individual designated as *M.* sp. #3 comparing it to one for *M*. *mittermeieri* from a previous study (Hunnicutt et al., 2020). The genomes were used to infer *N_e_* though time with Multiple Sequentially Markovian Coalescent (MSMC) analysis and compared those estimated from G-PhoCS (Fig. 6). Below, we describe the methods in some detail, while further details can be found in the Supplementary Material.

**Figure 2:**
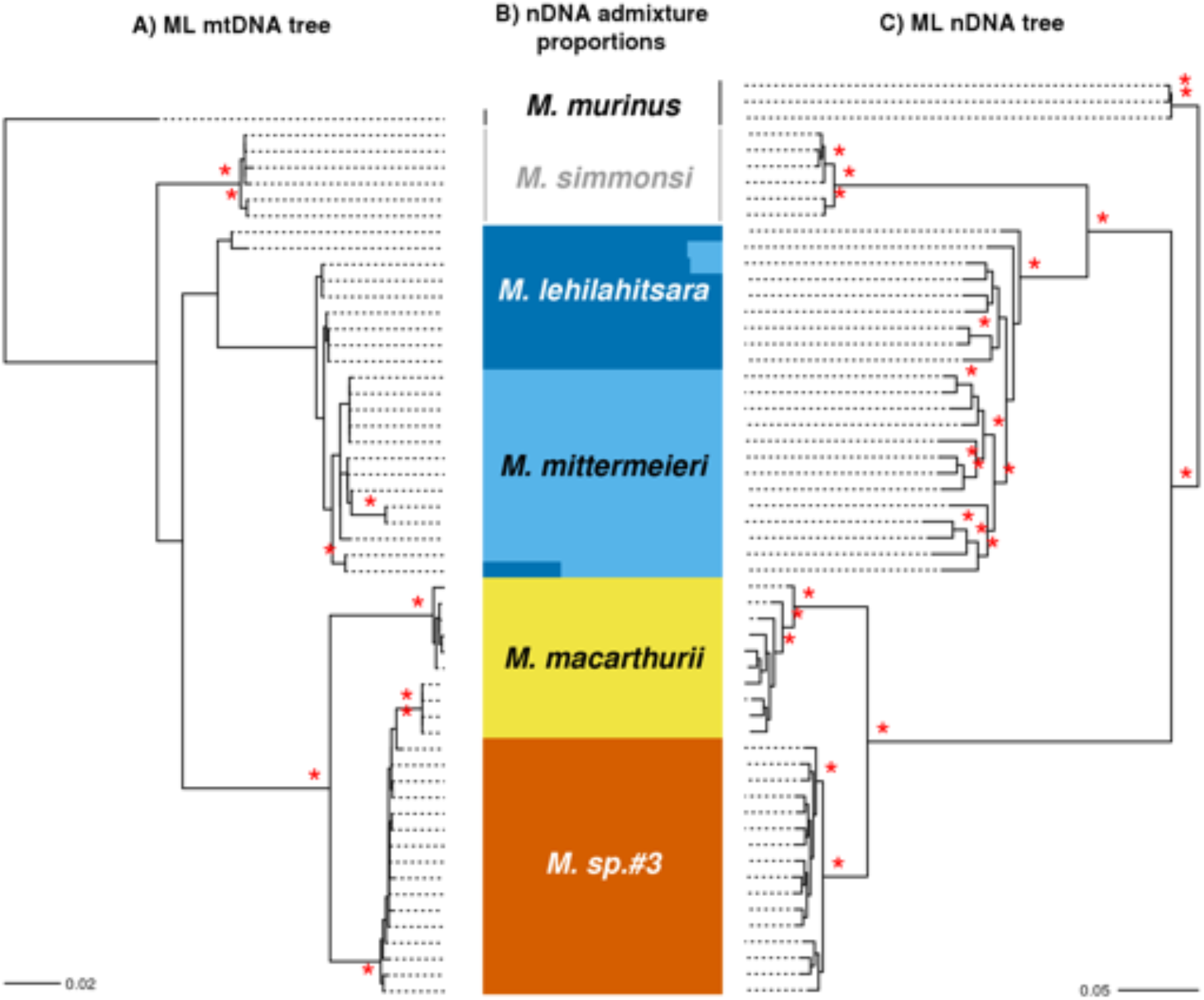
Phylogenetic relationships and ancestry proportions. **a)** Maximum-likelihood RAxML tree of 57 samples represented by 4,060 bp of mtDNA recovered from RADseq and Sanger sequencing (Table S1). The gray shaded box highlights individuals of *M*. *macarthurii* with *M*. sp. #3 mtDNA haplotypes. **b)** Clustering results for all species except the outgroup *M. murinus*, using NgsAdmix at *K* = 5. **c)** Maximum-likelihood RAXML tree obtained using RADseq nuclear data (nDNA). For all trees, *M. murinus* is used as the outgroup. In a) and c), bootstrap support values >90% are indicated above each node as a red asterisk. Clades

### Study Sites and Sampling

*Microcebus* samples were obtained by taking ~2 mm2 ear biopsies of captured (and thereafter released) individuals between 2008 and 2017 at seven humid evergreen forest sites (50-979 m a.s.l.) in the Analanjirofo and Sava regions of northeastern Madagascar (Fig. 1; Table S1). Additional samples were used from Riamalandy, Zahamena National Park (NP), Betampona Strict Nature Reserve (SNR) and Tampolo (Louis et al. 2006; Weisrock et al. 2010; Louis and Lei 2016) (Fig. 1). With this sampling strategy, we expected to include all mouse lemur species thought to occur in the region (from north to south): *M. mittermeieri*, *M. macarthurii*, *M*. sp. #3, *M. lehilahytsara*, and *M. simmonsi* (Fig. 1). *Microcebus murinus*, which occurs in western and southeastern Madagascar, was used as an outgroup.

### Sequencing Data, Genotyping and Genome Assembly

We generated RADseq libraries using the *Sbf*I restriction enzyme, following three protocols (Supplementary Methods, `Table S1`). Sequences were aligned to the *M.* sp. #3 nuclear genome generated by this study, and to the published *M. murinus* mitochondrial genome (LeCompte et al. 2016). We used two genotyping approaches to ensure robustness of our results. First, we estimated genotype likelihoods (GL) with ANGSD v0.92 (Nielsen et al. 2012; Korneliussen et al. 2014), which retains information about uncertainty in base calls, thereby alleviating some issues commonly associated with RADseq data such as unevenness in sequencing depth and allele dropout (Lozier 2014; Pedersen et al. 2018; Warmuth and Ellegren 2019). Second, we called genotypes with GATK v4.0.7.0 (DePristo et al. 2011), and filtered GATK genotypes following the “FS6” filter of O’Leary et al. (2018; their Table 2). We furthermore used three mtDNA fragments [Cytochrome Oxidase II (COII), Cytochrome B (cytB), and d-loop] that were amplified and Sanger sequenced for additional phylogenetic analyses.

The genome of the *M*. sp. #3 individual sampled in Mananara-Nord NP (`Table S3`) was sequenced with a single 500bp insert library on a single lane of an Illumina HiSeq 3000 with paired-end 150bp reads. We used MaSuRCA v3.2.2 (Zimin et al. 2013) for contig assembly and SSPACE (Boetzer et al. 2011) for scaffolding. Scaffolds potentially containing mitochondrial or X-chromosome sequence data were removed for downstream analyses.

### Phylogenetic Analyses

We used three phylogenetic approaches to infer relationships among lineages: (1) maximum likelihood using RAxML v8.2.11 (Stamatakis 2014), (2) SVDquartets, an MSC method that uses phylogenetic invariants, implemented in PAUP v4a163 (Chifman and Kubatko 2014), and (3) SNAPP, a full-likelihood MSC method for biallelic data that does not require joint gene tree estimation (v1.3.0; Bryant et al. 2012). Analyses with RAxML and SVDquartets used all available individuals, whereas SNAPP analyses were performed with subsets of 12 and 22 individuals for computational feasibility (see Supplementary Methods).

### Species Delimitation

#### Clustering approaches and summary statistics

Clustering analyses were performed using corresponding methods based on ANGSD genotype likelihoods [clustering in NgsAdmix v32 (Skotte et al. 2013) and PCA in ngsTools va4d338d (Fumagalli et al. 2014)] and on GATK-called genotypes [clustering in ADMIXTURE v1.3.0 (Alexander et al. 2009) and PCA using the glPca() function in adegenet v2.1.1 (Jombart and Ahmed 2011)]. These analyses were run for Clade I and II together and separately.

#### MSC-based approaches

We used SNAPP to test if the two lineages each in Clade I and II could be delimited using Bayes factors (Leaché et al. 2014), interpreting 2ln Bayes factors greater than six as strong evidence for a given model (Kass and Raftery 1995). We also applied guided species delimitation analyses with BPP (Yang and Rannala 2010; Rannala and Yang 2013) using full-length fasta files for a subset of individuals based on the species tree estimated by SVDquartets and SNAPP.

#### g*di*

Coalescent node heights (τ) and ancestral effective population sizes (θ) estimated by BPP were used to compute the genealogical divergence index (*gdi*; Jackson et al. 2017; Leaché et al. 2019) for the lineages in Clade I and II. We calculated *gdi* as in Leaché et al. (2019), using their equation 7 (*gdi* = 1-e^-2τ/θ^), where 2τ/θ represents the population divergence time between two taxa in coalescent units. θ is taken from one of the two taxa and therefore *gdi* was calculated twice for each species pair, alternating the focal taxon. We computed *gdi* using τ and θ parameter estimates for each posterior BPP sample to incorporate uncertainty in the estimates. Jackson et al. (2017) suggested the following interpretation of *gdi* values: the taxon pair (a) is unambiguously a single species for *gdi* < 0.2, (b) is unambiguously two separate species for *gdi* > 0.7, and (c) falls in an ambiguous zone for 0.7 > *gdi* < 0.2.

#### Isolation-by-distance

We tested for isolation-by-distance using the VCF file produced by GATK with the gl.ibd() function in the R package dartR 1.1.11 (Gruber et al. 2018).

### Inference of gene flow and divergence times

G-PhoCS v1.3 (Gronau et al. 2011), a Bayesian MSC approach that allows for the estimation of periods of gene flow (*i.e.* “migration bands”), was used to jointly infer divergence times, population sizes, and rates of gene flow between specific lineages. Based on the results of exploratory models each containing a single “migration band” between two lineages, we ran a final model with a migration band allowing gene flow from *mittermeieri* to *lehilahytsara*, and a migration band allowing gene flow from *macarthurii* to *M.* sp. #3. Given the observed mitonuclear discordance between *M.* sp. #3 and *M. macarthurii* (see Results), we investigated gene flow between them in more detail by running G-PhoCS using a dataset with only *M.* sp. #3, *M. macarthurii*, and *M. lehilahytsara* individuals, wherein *M.* sp. #3 was divided into two populations detected using clustering approaches.

The D-statistic and related formal statistics for admixture use phylogenetic invariants to infer post-divergence gene flow between non-sister populations or taxa. We used the qpDstat tool of admixtools v4.1 (Patterson et al. 2012) to compute four-taxon D-statistics for all possible configurations in which gene flow between non-sister lineages among the five ingroup lineages could be tested. We additionally tested for gene flow between *M. macarthurii* and *M.* sp. #3 by separately treating (1) the two distinct *M.* sp. #3 populations detected by clustering approaches, and (2) *M. macarthurii* individuals with and without “*M.* sp. #3-type” mtDNA (see Results). In all tests, *M. murinus* was used as P4 (outgroup).

### Effective Population Size Through Time

Studies have shown that population structure can generate spurious signals of population size change (Beaumont 2004; Chikhi et al. 2010; Heller et al. 2013). For example, sequentially Markovian coalescent approaches such as MSMC (Schiffels and Durbin 2014) actually estimate the inverse instantaneous coalescence rate, which is only equivalent to an effective size in panmictic models (Mazet et al. 2016; Rodríguez et al. 2018). We therefore inferred and compared population size histories using two methods. We estimated *N_e_* over time with MSMC for two species, using the whole-genome data of *M.* sp. #3 and *M*. *mittermeieri* (Hunnicutt et al. 2020) mapped to the chromosome-level genome assembly of *M*. *murinus* (Larsen et al. 2017). These estimates were compared to inferred changes in *N_e_* over time based on θ estimates from G-PhoCS for each predefined extant or ancestral population. Although the MSC was not expressly developed to estimate change in *N_e_* over time, this allowed us to examine broad demographic trends for relatively small population-level sampling with RADseq data, and to explicitly incorporate divergence events.

### Mutation Rate and Generation Time

We used empirical estimates of mutation rate and generation time to convert coalescent units from BPP, G-PhoCS and MSMC analyses into absolute times and population sizes. We incorporated uncertainty by drawing from mutation rate and generation time distributions for each sampled generation of the MCMC chains in BPP and G-PhoCS (MSMC parameter estimates were converted using the point estimates). For the mutation rate, we used a gamma distribution based on the mean (1.236 x 10^-8^) and variance (0.107 x 10^-8^) of seven pedigree-based mutation rate estimates for primates (see Campbell et al. 2019, `Table S1`). For the generation time, we used a lognormal distribution with a mean of ln(3.5) and standard deviation of ln(1.16) based on estimates of 4.5 years calculated from survival data (Zohdy et al. 2014; Yoder et al. 2016) from *M. rufus*, and 2.5 years from average parent age based on capture-mark-recapture and parentage data in the wild (Radespiel et al. 2019) for *M. murinus*.

## Results

### RADseq Data and Whole-Genome Assembly

We used three library generation protocols, two sequencing lengths, and a combination of single and paired-end sequencing, yielding data for all 63 individuals in the study and demonstrating the utility of cross-laboratory RAD sequencing, as previously shown in other taxa (e.g., Gonen et al. 2015). From more than 447 million raw reads (`Table S1`), over 394 million passed quality filters, with approximately 182 million successfully aligning to the *M.* sp. #3 reference genome. We obtained an average of 120,000 loci per individual with coverage ranging from ~1 to ~22x (*Table S1*).

We assembled approximately 2.5 Gb of nuclear genome sequence data for *M*. sp. #3 with a contig N50 around 36 Kb (`Table S3`). While the final assembly was fragmented, as expected for a single Illumina library genome, only 6.4% of mammalian BUSCOs were found to be missing. The genome sequence and associated gene annotations can be accessed through NCBI (Bioproject PRJNA512515).

### Phylogenetic Relationships

RAxML and SVDquartets recovered well-supported nDNA clades for *M. simmonsi*, *M. macarthurii*, and *M.* sp. #3, the latter two as sister taxa with 100% bootstrap support (Fig. 2; Fig. S2). SNAPP also supported *M.* sp. #3 as sister taxon to *M. macarthurii* (referred to as Clade I) and placed *M. lehilahytsara* as sister taxon to *M. mittermeieri* (referred to as Clade II) (`Fig. S2`). However, *M. lehilahytsara* was not monophyletic in RAxML analyses of nDNA (Fig. 2C) or mtDNA (Fig. 2A), and a SVDquartets analysis of nDNA placed one *M. lehilahytsara* individual from Ambavala as sister to all other *M. lehilahytsara* and *M. mittermeieri*, and only weakly supported a monophyletic *M. mittermeieri* (`Fig. S2A`).

Although mtDNA analyses placed several individuals from Anjiahely in a well-supported clade with *M.* sp. #3, individuals from Ambavala (see Fig. 1), Mananara-Nord NP, and Antanambe (Fig. 2A; see lower gray box), nuclear RADseq data placed them unambiguously within the *M. macarthurii* clade (Fig. 2B,C). This suggests that individuals from Anjiahely are in fact *M. macarthurii*, but carry two divergent mtDNA lineages, and that true *M.* sp. #3 are only found between Ambavala and Antanambe (Fig. 1). The cause of this mitonuclear discordance for *macarthurii* in Anjiahely was investigated further (see the section “Interspecific Gene Flow”).

### Species Delimitation

#### Genetic structure

A PCA with both pairs of sister lineages (Clade I: *M*. *macarthurii* and *M.* sp. #3; Clade II: *M*. *mittermeieri* and *M*. *lehilahytsara*) distinguished the two clades along PC1, and distinguished *M*. *macarthurii* and *M.* sp. #3 along PC2 (Fig. 3B). When restricting clustering analyses to Clade I, K = 2 was the best-supported number of clusters with both approaches, distinguishing *M*. *macarthurii* and *M.* sp. #3 (`Fig. S5; Fig. S7B`). At K = 3, *M.* sp. #3 was divided into two clusters with individuals from Mananara-Nord NP and Antanambe separated from Ambavala individuals (`Fig. S7B, Fig. S10`). A separate PCA analysis for Clade I also distinguished these two groups along PC2 (Fig. 3C), which we hereafter refer to as “southern *M*. sp. #3” (Mananara-Nord NP and Antanambe are south of the Mananara river) and “northern *M*. sp. #3” (Ambavala is north of the river, and 24.0 km from Mananara-Nord NP and 35.2 km from Antanambe; Fig. 1). When restricting clustering analyses to Clade II, ADMIXTURE and ngsAdmix suggested optimal values of 1 and 2, respectively; at K=2, *M*. *mittermeieri* and *M. lehilahytsara* were largely but not entirely separated by both approaches (`Fig. S5, Fig. S7C, Fig. S11`). A PCA distinguished *M*. *mittermeieri* and *M*. *lehilahytsara* along PC1 but with little separation (Fig. 3D).

**Figure 3:**
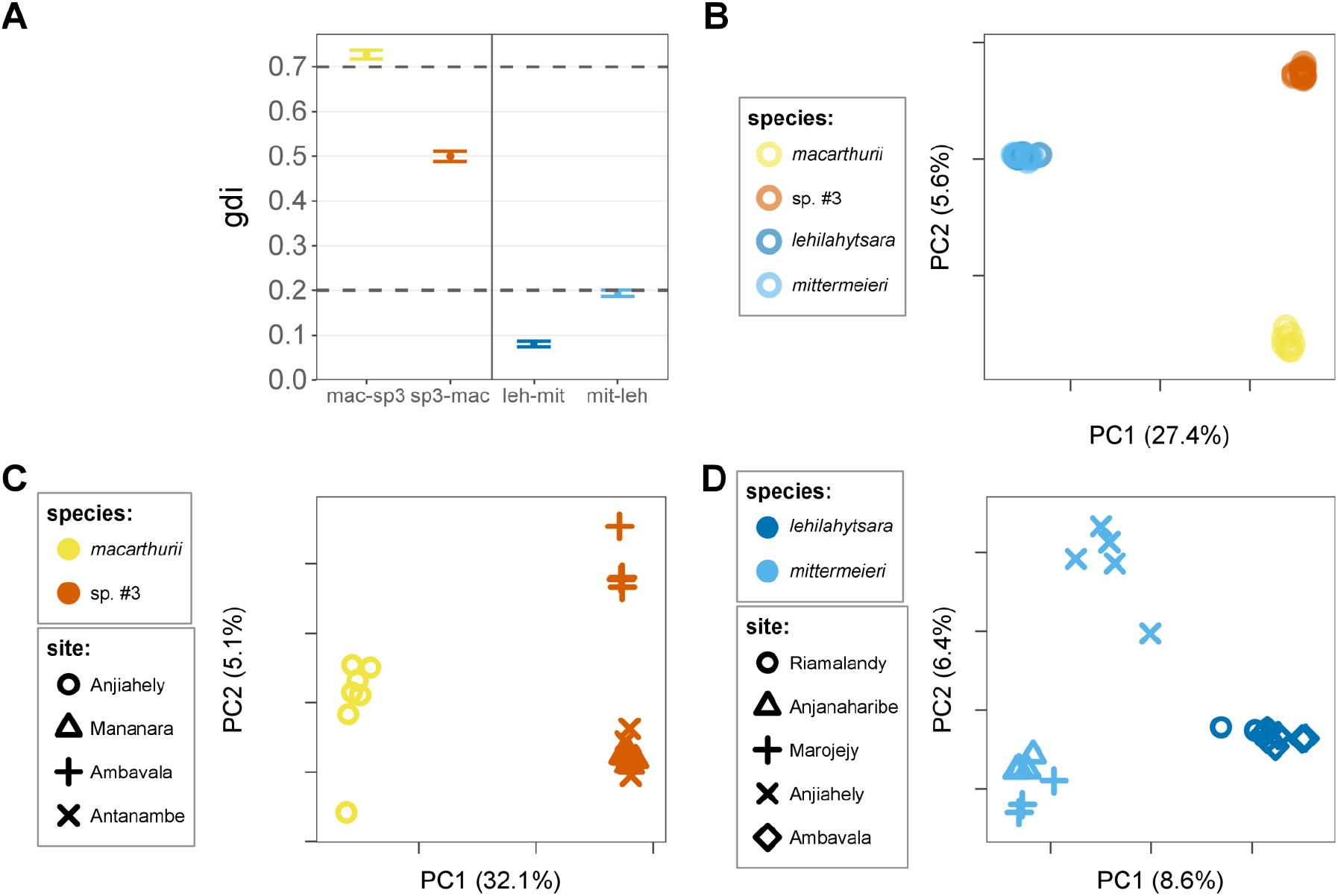
Population genetic structure and the gdi. **a)** Genealogical divergence index (gdi) for *M. macarthurii* – *M.* sp. #3 and *M. lehilahytsara* – *M. mittermeieri*. *gdi* values > 0.7 suggest separate species, *gdi* values < 0.2 are below the lower threshold for species delimitation, and 0.2 < *gdi* < 0.7 is an “ambiguous” range (Jackson et al. 2017). **b-d)** PCA analyses for b) all four species in Clades I and II, c) Clade I only: *M*. sp. #3 and *M. macarthurii*, with the former showing a split into two population groups: “northern” (Ambavala) and “southern” *M*. sp. #3 (Antanambe and Mananara-Nord NP), d) Clade II only: *M. lehilahytsara* and *M. mittermeieri.*

#### SNAPP and BPP

SNAPP Bayes factors strongly favored splitting Clade I into two species (2lnBF = 17,304 and 34,326 for two different datasets, `Table S6`), as well as splitting Clade II, although with a smaller difference in marginal likelihood scores (2lnBF = 1,828 and 993). All putative species assignments were recovered by the guided delimitation analysis with BPP (`Fig. S12`).

#### Genealogical divergence index (gdi)

For the Clade I sister pair, *gdi* was 0.727 (95% HPD: 0.718-0.737) from the perspective of *M*. *macarthurii* (i.e. above the upper threshold for species delimitation), and 0.500 (0.488-0.511) from the perspective of *M*. sp. #3 (i.e. in the upper ambiguous zone for species delimitation; Fig. 3A). In contrast, *gdi* values for the Clade II putative species pair were much lower and even below the lower threshold for species delimitation: 0.080 (0.074-0.086) from the perspective of *M*. *lehilahytsara*, and 0.193 (0.187-0.201) from the perspective of *M*. *mittermeieri* (Fig. 3A).

#### Isolation-by-distance (IBD)

While comparisons within and between lineages appeared to follow a single isolation-by-distance pattern for *M*. *mittermeieri* and *M*. *lehilahytsara* (Clade II, r=0.693, p=0.002, Fig. 4B), comparisons within versus between lineages differed strongly for *M*. *macarthurii* and *M.* sp. #3 (Clade I, Fig. 4A). Specifically, genetic distances between *M*. *macarthurii* and *M.* sp. #3 were much larger than within lineages and were also much larger than between *M*. *mittermeieri* and *M*. *lehilahytsara*, despite similar geographic distances.

**Figure 4:**
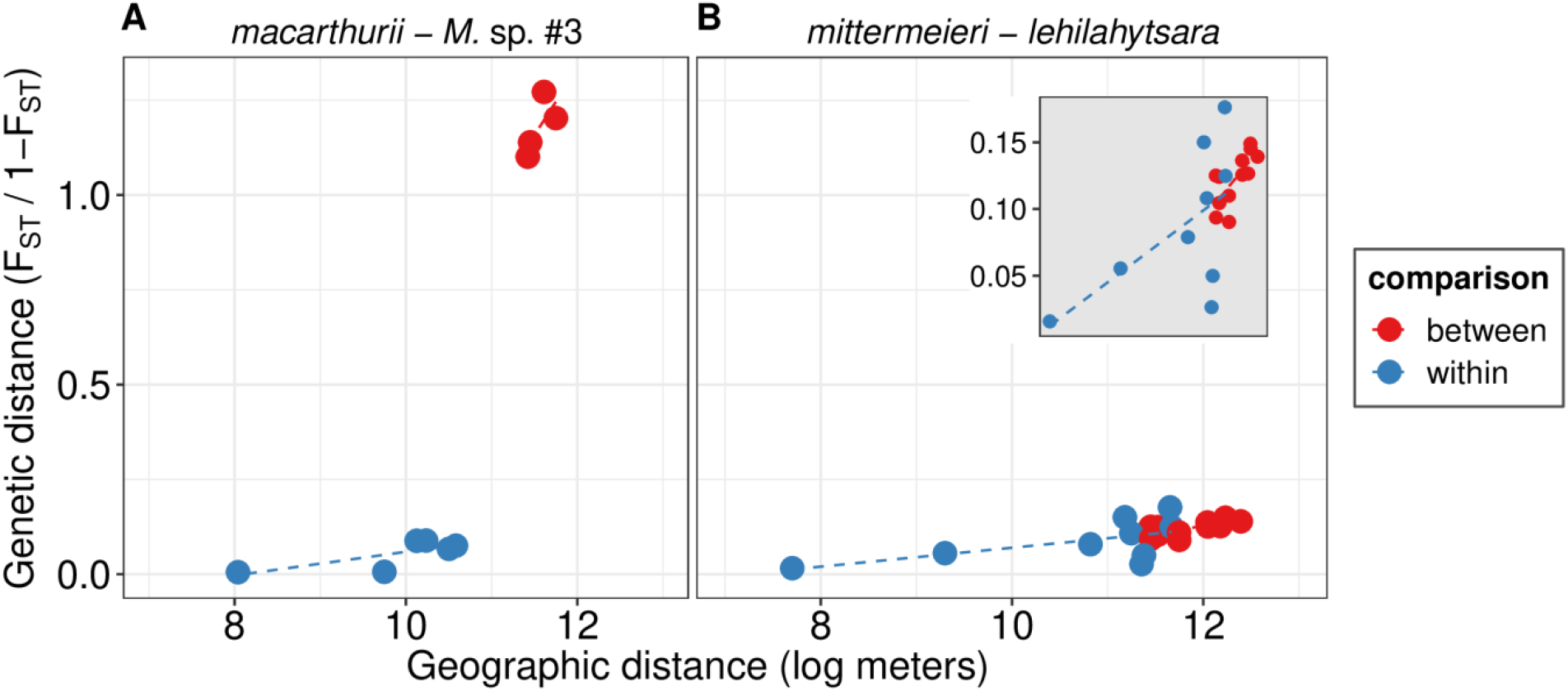
Patterns of isolation-by-distance in the two clades. **a)** Clade I (*M. macarthurii* and *M. sp. #3*). **b)** Clade II (*M. mittermeieri* and *M. lehilahytsara*). Population comparisons within lineages are shown as blue points, and comparisons between lineages are shown as red points. Both panels have the same y-axis scale, while the inset in B has a lower limit on the y-axis to better show the spread of points, given the smaller genetic distances between *M. mittermeieri* and *M. lehilahytsara*.

#### Interspecific gene flow

G-PhoCS inferred high levels of gene flow in Clade II, from *M*. *mittermeieri* to *M*. *lehilahytsara* [population migration rate (2Nm) = 1.59 (95% HPD: 1.50-1.68), migrants per generation: 0.18% (95% HPD: 0.09-0.27%)], and much lower levels of gene flow in Clade I, from *M.* sp. #3 to *macarthurii* [2Nm = 0.08 (95% HPD: 0.07-0.09), migrants per generation: 0.10% (0.05-0.15%)] (Fig. 5C). G-PhoCS also inferred low levels of gene flow between the two clades, most likely between ancestral populations, but the timing and direction of gene flow could not be determined (Supplementary Results; `Fig. S13`), and D-statistics testing for gene flow between the clades were not significant (`Fig. S14`).

**Figure 5:**
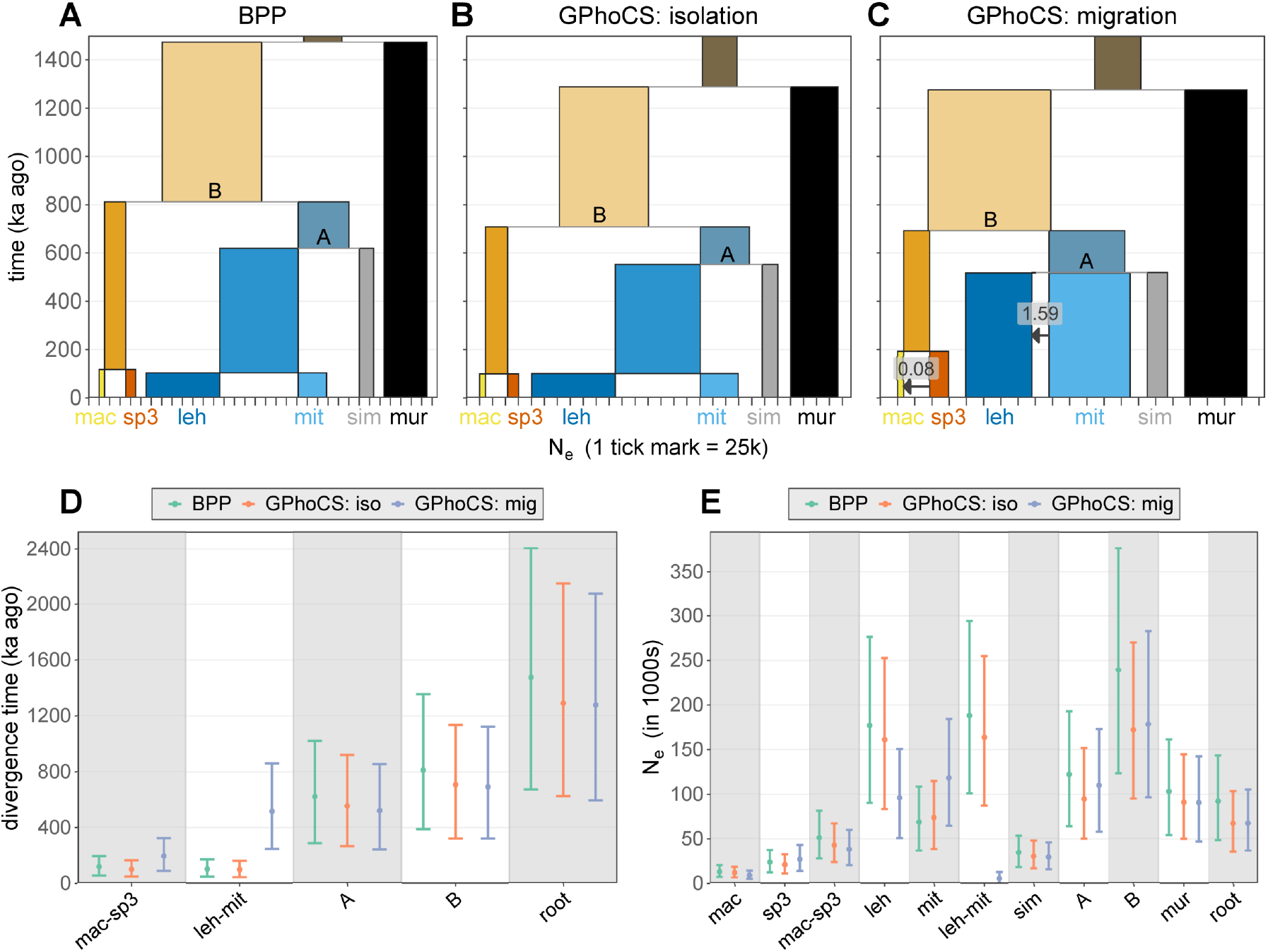
Demographic histories inferred by G-PhoCS and BPP. **a-c)** Divergence times (y-axis) and effective population sizes (x-axis) inferred with and without migration. Migration bands representing the estimated magnitude of gene flow are illustrated in **(c)**. **d-e)** Comparison of divergence times and effective population sizes for each node and lineage, respectively. The symbol “A” represents the lineage ancestral to *M. simmonsi*, *M. mittermeieri* and *M. lehilahytsara*, “B” represents the lineage ancestral to *M*. sp. #3, *M. macarthurii*, *M*. *simmonsi*, *M*. *mittermeieri* and *M*. *lehilahytsara*, and “root” represents the lineage ancestral to all six species included.

We further investigated gene flow between *M.* sp. #3 and *M*. *macarthurii* by taking the strong population structure within *M.* sp. #3 into account. D-statistics suggested that northern *M.* sp. #3 and *M*. *macarthurii* with “*M.* sp. #3-type” mtDNA share a slight excess of derived alleles in relation to southern *M*. sp. #3, significantly deviating from 0, which indicates gene flow (`Fig. S15A`). Using a G-PhoCS model with separate northern and southern groups for *M*. sp. #3, we found that (1) gene flow with *M. macarthurii* took place before and after the onset of divergence between northern and southern *M.* sp. #3, (2) gene flow between extant lineages occurred or occurs only between northern (and not southern) *M.* sp. #3 and *M*. *macarthurii*, and (3) gene flow is asymmetric, predominantly into *M*. *macarthurii* (`Fig. S15B`).

### Divergence Times

We estimated divergence times under the MSC model using BPP and G-PhoCS both with and without interspecific gene flow (Fig. 5; Fig. S17). Results were similar across these approaches, with the exception of divergence times between sister lineages in G-PhoCS models with versus without gene flow (Fig. 5). Specifically, the divergence time between *M.* sp. #3 and *M. macarthurii* (Clade I) without gene flow was estimated at 115 ka ago (95% HPD range: 52-190 ka across G-PhoCS and BPP models) (Fig. 5; Fig. S17), but at 193 ka ago (95% HPD: 89-318 ka) when incorporating gene flow (Fig. 5C-D). In Clade II, this difference in estimated divergence times was considerably larger: under an isolation model it was estimated to be 103 ka ago (95% HPD: 49-171 ka; Fig. 5) and as much as 520 ka ago (95% HPD: 249-871 ka) when modeled with gene flow (Fig. 5). Deeper nodes were not as strongly affected: divergence time between Clades I and II was estimated at 687 ka ago (95% HPD: 337-1126 ka) across G-PhoCS and BPP isolation models, and at 796 ka ago (95% HPD: 360-1311 ka) in a G-PhoCS model with gene flow (Fig. 5D).

### Effective Population Sizes

We found large differences in *N*e among lineages, with considerably larger *N*e for the lineages in Clade II, *M*. *lehilahytsara* (point estimate and 95% HPD range across the BPP and G-PhoCS models with and without interspecific gene flow: 159 k; 58-265 k) and *M*. *mittermeieri* (78 k; 36-140 k), than the lineages in Clade I, *M.* sp. #3 (24 k; 12-38 k) and *M*. *macarthurii* (12 k; 5-19 k) (Fig. 5A-C). Wide HPD intervals for *M*. *mittermeieri* and *lehilahytsara* are due to differences between models with and without gene flow. Using the G-PhoCS model focused on Clade I, fairly similar effective population sizes were estimated separately for northern (47 k; 17-78 k), southern (23 k; 12-37 k), and ancestral (33 k; 17-53 k) *M.* sp. #3 lineages (`Fig. S13`).

Using the whole-genome data for one individual of *M*. sp. #3 (from the southern group) and for *M*. *mittermeieri*, a comparison of MSMC analyses and G-PhoCS models with and without gene flow (Fig. 6) showed highly similar and markedly declining estimates of population sizes towards the present for *M*. sp. #3 (Fig. 6A). Estimates for *M*. *mittermeieri* were more variable across analyses and did not show a consistent decline towards the present (Fig. 6B).

**Figure 6:**
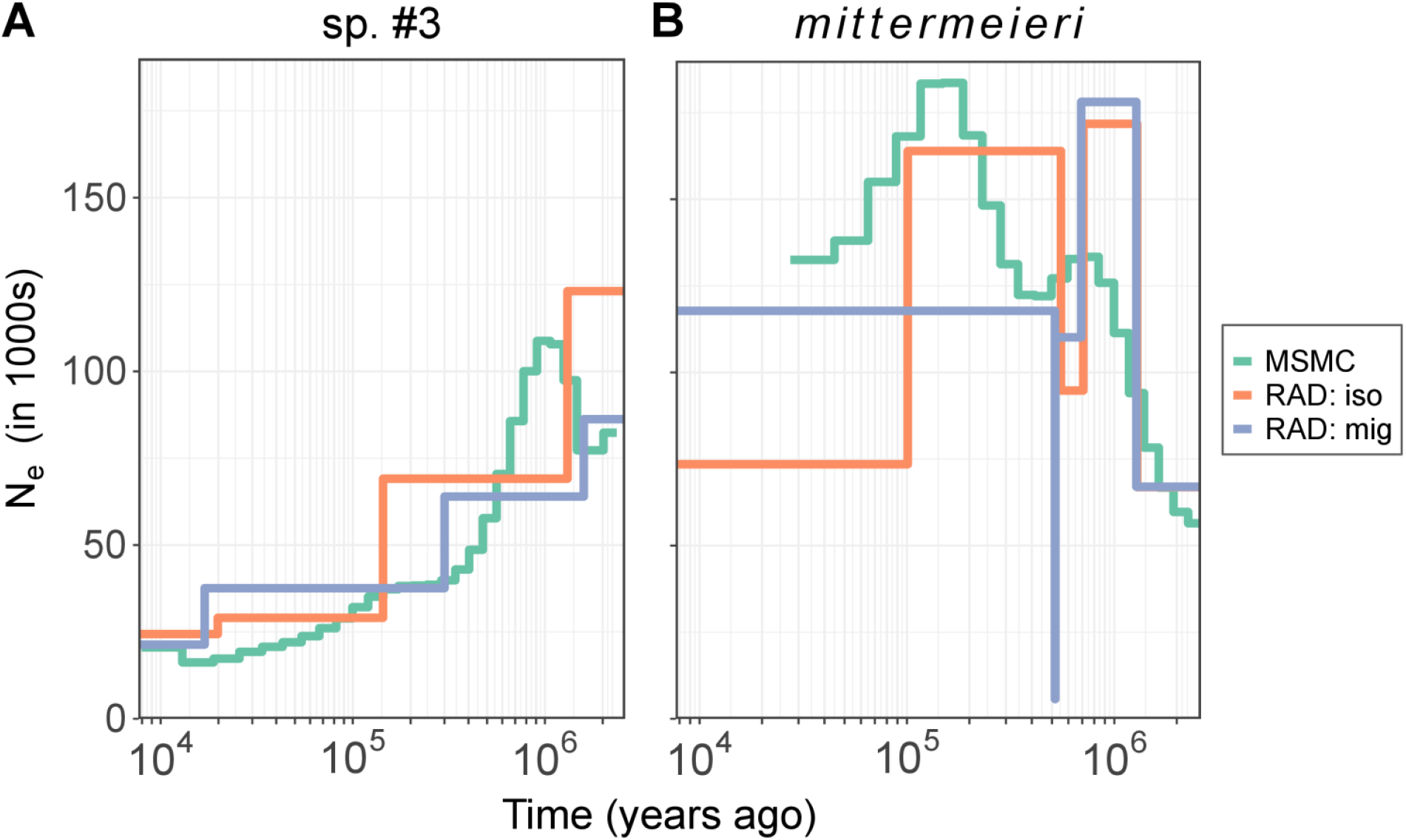
Estimates of effective population size through time for two species. Effective population sizes through time as inferred by MSMC for whole-genome data from a single individual (green lines), and by G-PhoCS for RADseq data without (“RAD: iso”, orange lines) and with (“RAD: mig”, blue lines) gene flow. **(A)** *M*. sp. #3. G-PhoCS analyses are shown for the southern *M*. sp. #3 population group and its ancestral lineages in the 3-species model given that the whole-genome individual was sampled from that population. **(B)** *M*. *mittermeieri*. G-PhoCS analyses are shown for *M*. *mittermeieri* and its ancestral lineages in the 5-species model. The sharp “jag” in the model with gene flow represents a small N_e_ estimate for the *M*. *mittermeieri-lehilahytsara* ancestor, which exists for an extremely short time in this model (see Fig. 6B,C), likely preventing proper estimation of N_e_.

## Discussion

We used a MSC-based framework for genomic species delimitation and identified rapid and recent diversification of mouse lemurs in a relatively small area in northeastern Madagascar. The same region was previously identified to harbor high levels of lemur microendemism that is vulnerable to the effects of climate change (Brown and Yoder 2015) and anthropogenic habitat alteration (Schüßler et al. 2020), marking it as a region of conservation concern. Species-level divergence was strongly supported for *M.* sp. #3 and its sister species *M. macarthurii* (Clade I, Fig. 2), but not for the pair of *M*. *mittermeieri* and *M. lehilahytsara* (Clade II, Fig. 2), despite our baseline assumption that the latter were distinct species (Olave et al., 2014). We inferred that the focal species all diverged from their common ancestors within the past million years and documented two cases of sympatric occurrence, each with one representative from Clade I and one from Clade II. The combined findings of recent divergence and sympatric overlap suggest that reproductive isolation can evolve rapidly in mouse lemurs.

### Support for Separate Sister Species Differs Sharply Between the Two Clades

Evidence for distinguishing *M.* sp. #3 and *M. macarthurii* as separate species was strong and consistent across analyses. They were reciprocally monophyletic across all phylogenetic analyses of RADseq data (Fig. 2C; Fig. S2; Fig. S3), separated unambiguously in clustering and PCA analyses (Fig. 2B; Fig. 3BC; Fig. S4; Fig. S6-S10), were strongly supported as separate lineages using SNAPP Bayes factors (`Table 1`) and BPP (`Fig. S12`), and passed the heuristic species delimitation criterion of *gdi* (Fig. 3A). A comparison of genetic and geographic distances moreover showed a clear distinction between intra- and interspecific genetic distances (Fig. 4). Finally, gene flow between these two lineages was estimated to have occurred at very low levels (G-PhoCS migration band = 0.08; Fig. 5c).

In contrast, separate species status of *M*. *lehilahytsara* and *M*. *mittermeieri* (Clade II) was not supported by genomic data. These species were paraphyletic in RAxML and SVDquartets analyses (Fig. 2A,C; Fig. S2) and not as clearly separated in clustering and PCA analyses (Fig. 2B; Fig. 3BD; Fig. S5, S7-S9; Fig. S10). Although the Bayes factor support from SNAPP was strong by standard guidelines (Kass and Raftery 1995), the evidence was much weaker relative to species in Clade I and decreased when more individuals were included (`Table S6`). It is unsurprising that Bayes factors will support splitting lineages with genetic structure (Sukumaran and Knowles 2017; Leaché et al. 2019) even with low levels of gene flow (Barley et al. 2018). Therefore, standard guidelines for interpreting Bayes factors may be of limited value for delimiting species, as informed by the lack of monophyly, high levels of inferred gene flow, and failure of additional delimitation tests observed here. Guided delimitation also separated *M*. *lehilahytsara* and *M*. *mittermeieri* (Fig. S11), but similar criticisms of oversplitting (e.g. Barley et al. 2018) lead us to not interpret MSC delimitation results as evidence of species status. Most strikingly, reciprocal *gdi* statistics for Clade II were <0.2, thus falling in the range suggested to unambiguously indicate a single species (Jackson et al. 2017; Leaché et al. 2019; Fig. 3A). Finally, comparing genetic and geographic distances within Clade II showed that a single isolation-by-distance pattern fits both intra-and interspecific comparisons (Fig. 4). While the range of *M*. *lehilahytsara* expands considerably further south than the populations examined here, our results strongly suggest that *M*. *mittermeieri* and *M*. *lehilahytsara* are best considered a single species. Sampling gaps are expected to cause false positive species delimitations rather than false negatives (Barley et al. 2018; Chambers and Hillis 2020; Mason et al. 2020), therefore additional sampling of *M. lehilahytsara* populations farther south should not affect our recommendation to synonymize *M. mittermeieri* as *M. lehilahytsara*.

### Mitonuclear Discordance and Gene Flow

Mitonuclear discordance was observed for a subset of *M*. *macarthurii* individuals from Anjiahely. These individuals carried mtDNA similar to that of *M.* sp. #3 (see Radespiel et al. 2008) but had nDNA indistinguishable from sympatric *M*. *macarthurii*. Although genealogical discordance could be due to incomplete lineage sorting (e.g., Heckman et al. 2007; Weisrock et al. 2010), mitochondrial introgression is supported by D-statistics (`Fig. S15`) and the inferred low levels of gene flow from the northern *M.* sp. #3 population into *M*. *macarthurii* by G-PhoCS (`Fig. S13`). Besides a possible case in Sgarlata et al. (2019), of mitochondrial introgression has not previously been reported in mouse lemurs. Somewhat curiously, the discovery of a divergent mtDNA lineage at Anjiahely (Radespiel et al. 2008), which prompted the current work, was apparently the result of mtDNA introgression from an undescribed species into its sister species.

### Population Size and Species Delimitation

The comparison of effective population sizes in Clades I and II reveals that they are markedly different, which can affect species delimitation tests such as *gdi* (Leaché et al., 2019). The *gdi* is calculated using population sizes and divergence times estimated under models with no gene flow, and since divergence time estimates in these models were highly similar in both clades (Fig. 5), differences in effective population sizes also appear to play a role in the stark difference in *gdi*. Indeed, *gdi* aims to quantify the probability that two sequences from the focal taxon coalesce more recently than the divergence time between the taxa, and larger effective population sizes result in slower sorting of ancestral polymorphisms (Maddison 1997).

Assessing “progress” in speciation by quantifying rates of neutral coalescence, however, implies that the magnitude of genetic drift is a good predictor of species limits. At least when considering reproductive isolation (i.e., biological species), this can be problematic, given that the role of drift in speciation is generally thought to be small (Rice and Hostert 1993; Coyne and Orr 2004; Czekanski-Moir and Rundell 2019; but see Uyeda et al. 2009). Therefore, additional measures of divergence should be taken into account, including those that do not depend on population size, such as rates of gene flow and divergence time itself (Yang and Rannala 2010; Leaché et al. 2019).

### Sympatric Occurrence and the Tempo of Speciation in Mouse Lemurs

Sympatric *Microcebus* species were found at two study sites, with a representative of each of the two focal clades in Anjiahely (*M*. *macarthurii* and *M*. *mittermeieri*) as well as in Ambavala (*M*. sp. #3 and *M*. *lehilahytsara*; Fig. 1). These cases of sympatric occurrence, with no evidence for recent admixture, imply that the two clades are reproductively isolated. Though our methods cannot address the mechanisms underlying reproductive isolation, possible barriers include male advertisement calls, which tend to differ strongly among species (Braune et al. 2008), and timing of reproduction, which has previously been found to differ among sympatric mouse lemur species (Schmelting et al. 2000; Evasoa et al. 2018) including the focal species (Schüßler et al., in revision). Only six other cases of sympatry among mouse lemur species are known, five of which include *M*. *murinus* as one of the co-occurring species (Radespiel 2016; Sgarlata et al. 2019).

Given that the sympatrically occurring species were estimated to have had a common ancestor as recently as ~700-800 ka ago (i.e., the divergence time between Clade I and Clade II, see Fig. 5), this suggests rapid evolution of reproductive isolation and a short time to sympatry among mouse lemurs. By comparison, Pigot & Tobias (2015) estimated that after 5 Ma of divergence, only 21–23% of primate species have attained sympatry. In fact, the one sympatric pair within their dataset of 74 sister species pairs younger than 2.5 million years consisted of the sympatric *Galago gallarum* and *G. senegalensis* (Pigot and Tobias 2015), which are also Strepsirrhini. Moreover, Curnoe et al. (2006) compiled data for naturally hybridizing primate species, and found the median estimated divergence time to be 2.9 Ma. More broadly, primate speciation rates do not appear to be lower than those for other mammals or even vertebrates (Curnoe et al. 2006, Upham et al. 2019). It should be noted, however, that the temporal estimates reported in our study are based on MSC analyses using mutation rates estimated from pedigree studies, whereas dates for other primate clades were largely calculated from fossil-calibrated relaxed clock methods.

### Complexities of Divergence Time Estimates

There are two noteworthy discrepancies in divergence time estimates highlighted by this study. First, the age estimate between the *M. mittermeieri* and *M. lehilahytsara* lineages increased from approximately 100 kya (Fig. 5b) to more than 500 kya (Fig. 5c) when the MSC model allowed for gene flow. The substantial effect of incorporating or disregarding gene flow on divergence time estimation has been previously noted (Leaché et al. 2014; Tseng et al. 2014) and we here reiterate its significance. Second, the coalescent-based estimates of divergence times presented here differ drastically from estimates based on fossil-calibrated relaxed-clock methods. In the present study, we estimated the mean age of the most recent common ancestor (MRCA) of mouse lemurs to be under 1.5 Ma, with the highest upper bound of 95% HPDs across models at 2.40 Ma. This age estimate is in stark contrast to previous fossil-calibrated estimates of 8 – 10 Ma (Yang and Yoder 2003; dos Reis et al. 2018).

Several factors likely contribute to this large difference. First, the MSC estimate uses a *de novo* mutation rate sampled from a distribution based on available pedigree-based mutation rates in primates, including mouse lemurs (Campbell et al., 2019). This rate is nearly two-fold higher than the estimated substitution rate for *M. murinus* (dos Reis et al., 2018). Second, converting coalescent units to absolute time also requires a generation time estimate. We attempted to account for uncertainty in generation time by similarly drawing from a distribution based on empirical parent age estimates (Zohdy et al., 2014; Radespiel et al., 2019) in mouse lemurs. Thus, either overestimation of the mutation rate and/or underestimation of the generation time would lead to divergence time estimates that are too recent. However, theoretical considerations suggest that instead, mouse lemur divergence time estimates from fossil-calibrated clock models are too old.

When incomplete lineage sorting is common, clock models that assume a single topology underlies all loci can overestimate species divergences compared to MSC estimates that allow gene trees to vary (Stange et al., 2018; Feng et al., 2020). This is likely to apply to mouse lemurs given that high levels of incomplete lineage sorting have been previously documented (Heckman et al., 2007; Weisrock et al., 2010; Hotaling et al., 2016). Moreover, due to the absence of a post-K-Pg terrestrial fossil record for Madagascar, clock-model estimates of divergence times in mouse lemurs have relied on fossil calibrations from the distantly-related African sister lineage of lemurs, the Lorisiformes (Seiffert et al., 2003), as well as from anthropoid primates and other mammals. This scenario - estimation of divergence times for younger, internal nodes with calibrations placed on much older nodes - is expected to lead to overestimation of divergence times (Angelis and dos Reis 2015). Therefore, it is likely that divergence times between mouse lemur species have been overestimated by previous studies with fossil-calibrated clock models (e.g. Yang and Yoder 2003; dos Reis et al., 2018), and we suggest that the mutation rate-calibrated MSC divergence times presented here are more accurate.

Our estimates of divergence times imply that the entire mouse lemur radiation originated in the Pleistocene, in turn suggesting that Pleistocene climatic oscillations represent a likely factor leading to geographic isolation and subsequent genetic divergence. Periods of drought during glacial maxima are hypothesized to have caused dramatic contraction of forest habitats (Burney et al. 1997; Gasse and Van Campo 2001; Wilmé et al. 2006; Kiage and Liu 2016) and to isolation of previously connected populations. Notably, the patterns of differentiation observed in this study are consistent with the predictions of Wilmé et al. (2006) wherein Madagascar’s river drainage systems created high-elevation retreat-dispersal corridors during periods of climatic oscillation. That is, whereas the lineages in Clade I (highly differentiated and low N_e_) appear to occur only in lowland forests, those in the Clade II (poorly differentiated and high Ne) occur at both higher and lower elevations (Schüßler et al., in revision). Moreover, the Mananara river runs between the fairly distinct northern and southern populations of *M*. sp. #3, further emphasizing the potential of large rivers to act as phylogeographic barriers in lemurs (Martin 1972; Pastorini et al. 2003; Goodman and Ganzhorn 2004; Olivieri et al. 2007).

### Population Size Dynamics

A long-term decline in population size was inferred for the lineage leading to *M*. sp. #3. While changes in inferred *N_e_* may be confounded by changes in population structure – especially for single-population sequential Markovian coalescent (PSMC/MSMC) models that do not explicitly consider population subdivision (Mazet et al. 2016; Chikhi et al. 2018) – we recovered similar results in both MSMC and G-PhoCS analyses (Fig. 6A). This congruence is especially persuasive given the underlying differences between the G-PhoCS and MSMC models and their input data. Moreover, Markovian coalescent approaches have been shown to be robust to genome assembly quality (Patton et al. 2019), yielding further confidence in the results. The inferred decline and population subdivision of *M*. sp. #3 was initiated long before anthropogenic land use, supporting the emerging consensus that human colonization in Madagascar alone does not explain the occurrence of open habitats and isolated forest fragments (Quéméré et al. 2012; Vorontsova et al. 2016; Yoder et al. 2016; Salmona et al. 2017, 2020; Hackel et al. 2018). Conversely, results for the *M. mittermeieri* lineage do not indicate a declining *N_e_* through time (Fig. 6B). This latter result may well be a simple corollary of the evidence described above, that this lineage is part of a single species complex represented by Clade II and thus occurs at both higher and lower elevations in northeastern Madagascar.

### Conclusions

We have shown that substantial mouse lemur diversity exists within a 130-km-wide stretch in northeastern Madagascar, including two instances of sympatric occurrence between representatives of two closely related clades. Within one of these clades, our integrative approach indicates that the undescribed lineage *M.* sp. #3 represents a distinct species, while the two named species in the other clade, *M. mittermeieri* and *M. lehilahytsara*, are better considered a single, widespread species with significant population structure. Given that the original description of *M. lehilahytsara* precedes that of *M. mittermeieri*, primate taxonomists should synonymize the two as *M. lehilahytsara.*

The divergence times calculated here using pedigree-based mutation rate estimates with the MSC are much younger than those of previous studies that used external fossil-based calibrations with concatenated likelihood methods. The younger dates suggest rapid evolution of reproductive isolation in mouse lemurs as well as a Pleistocene origin of the radiation, likely following population isolation due to climatic oscillations. This departure from previous hypotheses of mouse lemur antiquity emphasizes the need for future studies focused on resolving discrepancies in divergence time estimates, both in mouse lemurs and in other recently evolved organismal groups for which such comparisons have yet to be made.

## Author contributions

- Conception and design of study:

JP, GPT, DS, JS, LC, UR, ADY

- Data collection:

DS, MBB, JBA, EELJ, DWR, RMR, PK, JMR, BR collected samples in the field.

- Data analysis and interpretation:

DS, JS, LC, OB, PE, CRC, PAL, ARS, DW, AIP, PH, KEH, EJ, SM, RCW, EELJ, UR, ADY generated sequencing data.

JP, GPT, DS, JS, LC, UR, ADY conducted population genetic and phylogenetic analyses.

- Drafting and revising manuscript:

JP, GPT, DS, JS, LC, UR, ADY drafted the manuscript.

All co-authors revised and agreed on the last version of the manuscript.

## Acknowledgments

This study was conducted under the research permit No. 197/17/MEEF/SG/DGF/DSAP/SCB.Re (DS), 072/15/MEEMF/SG/DGF/DCB.SAP/SCB (MBB), 137/13/MEF/SG/DGF/DCB.SAP/SCB (DWR), 175/14/MEF/SG/DGF/DCB.SAP/SCB (A. Miller), kindly issued by the directeur du système des aires protégées, Antananarivo and the regional authorities (Direction Régional de l’Environnement, de l’Ecologie et de Forêts). We are endebted to J.H. Ratsimbazafy, N.V. Andriaholinirina, C. Misandeau, B. Le Pors and S. Rasoloharijaona, for their help with administrative tasks, to A. Miller for sharing samples, and to G. Besnard for facilitating this study. We thank our field assistants (T. Ralantoharijaona, I. Sitrakarivo, C. Hanitriniaina and T. Ralantoharijaona), the Wildlife Conservation Society Madagascar and the ADAFAM (Association Des Amis de la Forêt d’Ambodiriana-Manompana, C. Misandeau in particular) for their valuable help during sample collection. We warmly thank the many local guides and cooks for sharing their incomparable expertise and help in the field, *misaotra anareo jiaby*.

Funding was granted by the Bauer Foundation and the Zempelin Foundation of the “Deutsches Stiftungszentrum” under grant no. T237/22985/2012/kg and T0214/32083/2018/sm to DS, Duke Tropical Conservation Initiative Grant to ADY, and Duke Lemur Center/SAVA Conservation research funds to MBB, the School of Animal Biology at The University of Western Australia to AM, the Fundação para a Ciência e a Tecnologia, Portugal (PTDC/BIA-BEC/100176/2008, PTDC/BIA-BIC/4476/2012, and SFRH/BD/64875/2009), the Groupement de Recherche International (GDRI) Biodiversité et développement durable – Madagascar, the Laboratoire d’Excellence (LABEX) TULIP (ANR-10-LABX-41) and CEBA (ANR-10-LABX-25-01, the Instituto Gulbenkian de Ciência, Portugal to LC and JS, the ERA-NET BiodivERsA project: INFRAGECO (Inference, Fragmentation, Genomics, and Conservation, ANR-16-EBI3-0014 & FCT-Biodiversa/0003/2015) the LIA BEEG-B (Laboratoire International Associé – Bioinformatics, Ecology, Evolution, Genomics and Behaviour, CNRS) to LC and JS. Further financial support came from the Institute of Zoology, University of Veterinary Medicine Hannover and UR acknowledges the long-term support of the late Elke Zimmermann for her research activities on Madagascar. The genomic data were generated with funds from NSF DEB-1354610 to ADY and DWW and from the EDB Lab to JS. ADY also gratefully acknowledges support from the John Simon Guggenheim Memorial Foundation and the Alexander von Humboldt Foundation. EELJ would like to acknowledge support from the Ahmanson Foundation for the data generation. This work was performed in collaboration with the GeT core facility, Toulouse, France (http://get.genotoul.fr), and was supported by France Génomique National infrastructure, funded as part of “Investissement d’avenir” program managed by Agence Nationale pour la Recherche (contract ANR-10-INBS-09). JS, UR & LC also gratefully acknowledge support from the Get-Plage sequencing and Genotoul bioinformatics (BioinfoGenotoul) platforms Toulouse Midi-Pyrenees. This is DLC publication #XXXX.

## References

Alexander D.H., Novembre J., Lange K. 2009. Fast model-based estimation of ancestry in unrelated individuals. Genome Res. 19:1655–1664.

Ali O.A., O’Rourke S.M., Amish S.J., Meek M.H., Luikart G., Jeffres C., Miller M.R. 2016. RAD capture (Rapture): Flexible and efficient sequence-based genotyping. Genetics. 202:389–400.

Angelis K., Dos Reis M. 2015. The impact of ancestral population size and incomplete lineage sorting on Bayesian estimation of species divergence times. Curr. Zool. 61:874–885.

Barley A.J., Brown J.M., Thomson R.C. 2018. Impact of model violations on the inference of species boundaries under the multispecies coalescent. Syst. Biol. 67:269–284.

Beaumont M.A. 2004. Recent developments in genetic data analysis: what can they tell us about human demographic history? Heredity. 92:365–379.

Bickford D., Lohman D.J., Sodhi N.S., Ng P.K.L., Meier R., Winker K., Ingram K.K., Das I. 2007. Cryptic species as a window on diversity and conservation. Trends Ecol. Evol. 22:148–155.

Blanco M.B., Rasoazanabary E., Godfrey L.R. 2015. Unpredictable environments, opportunistic responses: Reproduction and population turnover in two wild mouse lemur species (Microcebus rufus and M. griseorufus) from eastern and western Madagascar. Am. J. Primatol. 77:936–947.

Boetzer M., Henkel C.V., Jansen H.J., Butler D., Pirovano W. 2011. Scaffolding pre-assembled contigs using SSPACE. Bioinformatics. 27:578–579.

Braune P., Schmidt S., Zimmermann E. 2008. Acoustic divergence in the communication of cryptic species of nocturnal primates (*Microcebus* ssp.). BMC Biology. 6:19.

Brown J.L., Yoder A.D. 2015. Shifting ranges and conservation challenges for lemurs in the face of climate change. Ecology and Evolution. 5:1131–1142.

Bryant D., Bouckaert R., Felsenstein J., Rosenberg N.A., RoyChoudhury A. 2012. Inferring species trees directly from biallelic genetic markers: bypassing gene trees in a full coalescent analysis. Mol. Biol. Evol. 29:1917–1932.

Burney D., James H., Grady F., Rafamantanantsoa J.-G., Ramilisonina, Wright H., Cowart J. 1997. Environmental change, extinction and human activity: evidence from caves in NW Madagascar. Journal of Biogeography. 24:755–767.

Campbell C.R., Tiley G.P., Poelstra J.W., Hunnicutt K.E., Larsen P.A., Reis M. dos, Yoder A.D. 2019. Pedigree-based measurement of the de novo mutation rate in the gray mouse lemur reveals a high mutation rate, few mutations in CpG sites, and a weak sex bias. bioRxiv.:724880.

Carstens B.C., Dewey T.A. 2010. Species delimitation using a combined coalescent and information-theoretic approach: an example from North American *Myotis* bats. Syst. Biol. 59:400–414.

Chambers E.A., Hillis D.M. 2020. The multispecies coalescent over-splits species in the case of geographically widespread taxa. Syst. Biol. 69:184–193.

Chifman J., Kubatko L. 2014. Quartet inference from SNP data under the coalescent model. Bioinformatics. 30:3317–3324.

Chikhi L., Rodríguez W., Grusea S., Santos P., Boitard S., Mazet O. 2018. The IICR (inverse instantaneous coalescence rate) as a summary of genomic diversity: insights into demographic inference and model choice. Heredity. 120:13–24.

Chikhi L., Sousa V.C., Luisi P., Goossens B., Beaumont M.A. 2010. The confounding effects of population structure, genetic diversity and the sampling scheme on the detection and quantification of population size change. Genetics. 186:983–995.

Curnoe D., Thorne A., Coate J.A. 2006. Timing and tempo of primate speciation. J. Evol. Biol. 19:59–65.

Czekanski-Moir J.E., Rundell R.J. 2019. The ecology of nonecological speciation and nonadaptive radiations. Trends Ecol. Evol. 34:400–415.

Dalquen D.A., Zhu T., Yang Z. 2017. Maximum likelihood implementation of an isolation-with-migration model for three species. Syst. Biol. 66:379–398.

Dávalos L.M., Russell A.L. 2014. Sex-biased dispersal produces high error rates in mitochondrial distance-based and tree-based species delimitation. J. Mammal. 95:781–791.

DePristo M.A., Banks E., Poplin R., Garimella K.V., Maguire J.R., Hartl C., Philippakis A.A., del Angel G., Rivas M.A., Hanna M., McKenna A., Fennell T.J., Kernytsky A.M., Sivachenko A.Y., Cibulskis K., Gabriel S.B., Altshuler D., Daly M.J. 2011. A framework for variation discovery and genotyping using next-generation DNA sequencing data. Nature Genetics. 43:491–498.

Dincă V., Lee K.M., Vila R., Mutanen M. 2019. The conundrum of species delimitation: a genomic perspective on a mitogenetically super-variable butterfly. Proc. R. Soc. Lond. B Biol. Sci. 286:20191311.

dos Reis M., Gunnell G.F., Barba-Montoya J., Wilkins A., Yang Z., Yoder A.D. 2018. Using phylogenomic data to explore the effects of relaxed clocks and calibration strategies on divergence time estimation: primates as a test case. Syst. Biol. 67:594–615.

Du Puy D.J., Moat J. 1998. Vegetation mapping and classification in Madagascar (using GIS): implications and recommendations for the conservation of biodiversity. In: Huxley C., Lock J., Cutler D., editors. Chorology, Taxonomy and Ecology of the Floras of Africa and Madagascar. Kew: Royal Botanical Gardens. p. 97–117.

Edwards D.L., Knowles L.L. 2014. Species detection and individual assignment in species delimitation: can integrative data increase efficacy? Proc. R. Soc. Lond. B Biol. Sci. 281:20132765.

Eriksson A., Manica A. 2012. Effect of ancient population structure on the degree of polymorphism shared between modern human populations and ancient hominins. Proc. Natl. Acad. Sci. U.S.A. 109:13956–13960.

Estrada A., Garber P.A., Rylands A.B., Roos C., Fernandez-Duque E., Fiore A.D., Nekaris K.A.-I., Nijman V., Heymann E.W., Lambert J.E., Rovero F., Barelli C., Setchell J.M., Gillespie T.R., Mittermeier R.A., Arregoitia L.V., Guinea M. de, Gouveia S., Dobrovolski R., Shanee S., Shanee N., Boyle S.A., Fuentes A., MacKinnon K.C., Amato K.R., Meyer A.L.S., Wich S., Sussman R.W., Pan R., Kone I., Li B. 2017. Impending extinction crisis of the world’s primates: Why primates matter. Science Advances. 3:e1600946.

Feder J.L., Egan S.P., Nosil P. 2012. The genomics of speciation-with-gene-flow. Trends in Genetics. 28:342–350.

Feng B, Merilä J, Matschiner M, Momigliano P. 2020. Estimating uncertainty in divergence times among three-spined stickleback clades using the multispecies coalescent. Mol Phylogent Evol. 142:106646

Flouri T., Jiao X., Rannala B., Yang Z. 2018. Species tree inference with BPP using genomic sequences and the multispecies coalescent. Mol. Biol. Evol. 35:2585–2593.

Fujita M.K., Leaché A.D., Burbrink F.T., McGuire J.A., Moritz C. 2012. Coalescent-based species delimitation in an integrative taxonomy. Trends Ecol. Evol. 27:480–488.

Fumagalli M., Vieira F.G., Linderoth T., Nielsen R. 2014. ngsTools: methods for population genetics analyses from next-generation sequencing data. Bioinformatics. 30:1486–1487.

Gasse F., Van Campo E. 2001. Late Quaternary environmental changes from a pollen and diatom record in the southern tropics (Lake Tritrivakely, Madagascar). Palaeogeography, Palaeoclimatology, Palaeoecology. 167:287–308.

Gavrilets S., Boake C.R. 1998. On the evolution of premating isolation after a founder event. Am. Nat. 152:706–716.

Gonen S., Bishop S.C., Houston R.D. 2015. Exploring the utility of cross-laboratory RAD-sequencing datasets for phylogenetic analysis. BMC Research Notes. 8:299.

Goodman S.M., Benstead J.P. 2005. Updated estimates of biotic diversity and endemism for Madagascar. Oryx. 39:73–77.

Goudet J. 2005. hierfstat, a package for r to compute and test hierarchical F-statistics. Mol. Ecol. Notes. 5:184–186.

Gronau I., Hubisz M.J., Gulko B., Danko C.G., Siepel A. 2011. Bayesian inference of ancient human demography from individual genome sequences. Nat. Genet. 43:1031–1034.

Gruber B., Unmack P.J., Berry O.F., Georges A. 2018. dartr: An r package to facilitate analysis of SNP data generated from reduced representation genome sequencing. Mol. Ecol. Res. 18:691–699.

Grummer J.A., Bryson R.W., Reeder T.W. 2014. Species delimitation using Bayes factors: simulations and application to the *Sceloporus scalaris* species group (Squamata: Phrynosomatidae). Syst. Biol. 63:119–133.

Hackel J., Vorontsova M.S., Nanjarisoa O.P., Hall R.C., Razanatsoa J., Malakasi P., Besnard G. 2018. Grass diversification in Madagascar: In situ radiation of two large C3 shade clades and support for a Miocene to Pliocene origin of C4 grassy biomes. Journal of Biogeography. 45:750–761.

Heckman K.L., Mariani C.L., Rasoloarison R., Yoder A.D. 2007. Multiple nuclear loci reveal patterns of incomplete lineage sorting and complex species history within western mouse lemurs (*Microcebus*). Mol. Phylogenet. Evol. 43:353–367.

Heller R., Chikhi L., Siegismund H.R. 2013. The confounding effect of population structure on Bayesian skyline plot inferences of demographic history. PLoS ONE. 8:e62992.

Huang J.-P., Leavitt S.D., Lumbsch H.T. 2018. Testing the impact of effective population size on speciation rates –a negative correlation or lack thereof in lichenized fungi. Scientific Reports. 8:1–6.

Hundsdoerfer A.K., Lee K.M., Kitching I.J., Mutanen M. 2019. Genome-wide SNP data reveal an overestimation of species diversity in a group of hawkmoths. Genome Biol. Evol. 11:2136–2150.

Hunnicutt K.E., Tiley G.P., Williams R.C., Larsen P.A., Blanco M.B., Rasoloarison R.M., Campbell C.R., Zhu K., Weisrock D.W., Matsunami H., Yoder A.D. 2020. Comparative genomic analysis of the pheromone receptor class 1 family (V1R) reveals extreme complexity in mouse lemurs (Genus, *Microcebus*) and a chromosomal hotspot across mammals. Genome Biol. Evol. 12:3562–3579.

Jackson N.D., Carstens B.C., Morales A.E., O’Meara B.C. 2017. Species delimitation with gene flow. Syst. Biol. 66:799–812.

Jombart T., Ahmed I. 2011. adegenet 1.3-1: new tools for the analysis of genome-wide SNP data. Bioinformatics. 27:3070–3071.

Kappeler P.M., Rasoloarison R.M., Razafimanantsoa L., Walter L., Roos, C. 2005. Morphology, behaviour and molecular evolution of giant mouse lemurs (Mirza spp.) Gray, 1870, with description of a new species. Primate Rep. 71:3–26.

Kass R.E., Raftery A.E. 1995. Bayes Factors. J. Am. Stat. Assoc. 90:773–795.

Kiage L.M., Liu K. 2016. Late Quaternary paleoenvironmental changes in East Africa: a review of multiproxy evidence from palynology, lake sediments, and associated records. Progress in Physical Geography. 30:633–658.

Kim B.Y., Wei X., Fitz-Gibbon S., Lohmueller K.E., Ortego J., Gugger P.F., Sork V.L. 2018. RADseq data reveal ancient, but not pervasive, introgression between Californian tree and scrub oak species (*Quercus* sect. *Quercus*: *Fagaceae*). Mol. Ecol. 27:4556–4571.

Knaus B.J., Grünwald N.J. 2017. vcfr: a package to manipulate and visualize variant call format data in R. Mol. Ecol. Res. 17:44–53.

Knoop S., Chikhi L., Salmona J. 2017. Mouse lemurs’ and degraded habitat. bioRxiv.:216382.

Korneliussen T.S., Albrechtsen A., Nielsen R. 2014. ANGSD: Analysis of Next Generation Sequencing Data. BMC Bioinformatics. 15:356.

Langmead B., Trapnell C., Pop M., Salzberg S.L. 2009. Ultrafast and memory-efficient alignment of short DNA sequences to the human genome. Genome Biology. 10:R25.

Leaché A.D., Fujita M.K., Minin V.N., Bouckaert R.R. 2014. Species delimitation using genome-wide SNP data. Syst. Biol. 63:534–542.

Leaché A.D., Zhu T., Rannala B., Yang Z. 2019. The spectre of too many species. Syst Biol. 68:168–181.

LeCompte E., Crouau-Roy B., Aujard F., Holota H., Murienne J. 2016. Complete mitochondrial genome of the gray mouse lemur, *Microcebus murinus* (Primates, Cheirogaleidae). Mitochondrial DNA A DNA Mapp. Seq. Anal. 27:3514–3516.

Linck E., Epperly K., Van Els P., Spellman G.M., Bryson R.W., McCormack J.E., Canales-Del-Castillo R., Klicka J. 2019. Dense geographic and genomic sampling reveals paraphyly and a cryptic lineage in a classic sibling species complex. Syst. Biol. 68:956–966.

Louis E.E., Coles M.S., Andriantompohavana R., Sommer J.A., Engberg S.E., Zaonarivelo J.R., Mayor M.I., Brenneman R.A. 2006. Revision of the mouse lemurs (*Microcebus*) of eastern Madagascar. Int. J. Primatol. 27:347–389.

Louis E.E. Jr., Engberg S.E., McGuire S.M., McCormick M.J., Randriamampionona R., Ranaivoarisoa J.F., Bailey C.A., Mittermeier R.A., Lei R. 2008. Revision of the mouse lemurs, *Microcebus* (Primates, Lemuriformes), of northern and northwestern Madagascar with descriptions of two new species at Montagne d’Ambre National Park and Antafondro Classified Forest. Primate Conservation. 23:19–38.

Louis E.E. Jr., Lei R. 2016. Mitogenomics of the family Cheirogaleidae and relationships to taxonomy and biogeography in Madagascar. In: Lehman S., Radespiel U., Zimmermann E., editors. The Dwarf and Mouse Lemurs of Madagascar: Biology, Behavior and Conservation Biogeography of the Cheirogaleidae. Cambridge, UK: Cambridge University Press. p. 54–93.

Lozier J.D. 2014. Revisiting comparisons of genetic diversity in stable and declining species: assessing genome-wide polymorphism in North American bumble bees using RAD sequencing. Mol. Ecol. 23:788–801.

Luo A., Ling C., Ho S.Y.W., Zhu C.-D. 2018. Comparison of methods for molecular species delimitation across a range of speciation scenarios. Syst. Biol. 67:830–846.

Maddison W.P. 1997. Gene trees in species trees. Syst. Biol. 46:523–536.

Mallet J., Besansky N., Hahn M.W. 2016. How reticulated are species? Bioessays. 38:140–149.

Markolf M., Brameier M., Kappeler P.M. 2011. On species delimitation: Yet another lemur species or just genetic variation? BMC Evol. Biol. 11:216.

Mason NA, Fletcher NK, Gill BA, Funk WC, Zamudio KR. 2020. Coalescent-based species delimitation is sensitive to geographic sampling and isolation by distance. Systematics and Biodiversity, 18:269–280.

Matute D.R. 2013. The role of founder effects on the evolution of reproductive isolation. J. Evol. Biol. 26:2299–2311.

Mazet O., Rodríguez W., Grusea S., Boitard S., Chikhi L. 2016. On the importance of being structured: instantaneous coalescence rates and human evolution--lessons for ancestral population size inference? Heredity. 116:362–371.

McLaughlin J.F., Winker K. 2020. An empirical examination of sample size effects on population demographic estimates in birds using single nucleotide polymorphism (SNP) data. bioRxiv.:2020.03.10.986463.

Mittermeier R. A., Louis E. E. Jr., Richardson M., Schwitzer C., Langrand O., Rylands A. B., Hawkins F., Rajaobelina S., Ratsimbazafy J., Rasoloarison R., Roos C., Kappeler P. M., Mackinnon J.. 2010. Lemurs of Madagascar, 3rd Edition. Conservation International Tropical Field Guide Series, Washington, USA.

Myers N., Mittermeier R.A., Mittermeier C.G., da Fonseca G.A.B., Kent J. 2000. Biodiversity hotspots for conservation priorities. Nature. 403:853–858.

Nielsen R., Korneliussen T., Albrechtsen A., Li Y., Wang J. 2012. SNP calling, genotype calling, and sample allele frequency estimation from new-generation sequencing data. PLoS ONE. 7:e37558.

O’Leary S.J., Puritz J.B., Willis S.C., Hollenbeck C.M., Portnoy D.S. 2018. These aren’t the loci you’re looking for: Principles of effective SNP filtering for molecular ecologists. Mol. Ecol. 27:3193–3206.

Olave M, Sola E, Knowles LL. 2014. Upstream analyses create problems with DNA-based species delimitation. Syst Biol, 63:263–271.

Olivieri G, Zimmermann E, Randrianambinina B, Rasoloharijaona S, Rakotondravony D, Guschanski K, Radespiel U. 2007. The ever-increasing diversity in mouse lemurs: Three new species in north and northwestern Madagascar. Mol. Phy. Evol., 43:309–327.

Palkopoulou E., Lipson M., Mallick S., Nielsen S., Rohland N., Baleka S., Karpinski E., Ivancevic A.M., To T.-H., Kortschak R.D., Raison J.M., Qu Z., Chin T.-J., Alt K.W., Claesson S., Dalén L., MacPhee R.D.E., Meller H., Roca A.L., Ryder O.A., Heiman D., Young S., Breen M., Williams C., Aken B.L., Ruffier M., Karlsson E., Johnson J., Palma F.D., Alfoldi J., Adelson D.L., Mailund T., Munch K., Lindblad-Toh K., Hofreiter M., Poinar H., Reich D. 2018. A comprehensive genomic history of extinct and living elephants. Proc. Natl. Acad. Sci. U.S.A. 115:E2566–E2574.

Pamilo P., Nei M. 1988. Relationships between gene trees and species trees. Mol. Biol. Evol. 5:568–583.

Patton AH, Margres MJ, Stahlke AR, Hendricks S, Lewallen K, Hamede RK, Ruiz-Aravena M, Ryder O, McCallum HI, Jones ME, et al. 2019. Contemporary Demographic Reconstruction Methods Are Robust to Genome Assembly Quality: A Case Study in Tasmanian Devils. Mol Biol Evol, 36:2906–2921.

Pastorini J., Thalmann U., Martin R.D. 2003. A molecular approach to comparative phylogeography of extant Malagasy lemurs. Proc. Natl. Acad. Sci. U.S.A. 100:5879–5884.

Patterson N., Moorjani P., Luo Y., Mallick S., Rohland N., Zhan Y., Genschoreck T., Webster T., Reich D. 2012. Ancient admixture in human history. Genetics. 192:1065–1093.

Pedersen C.-E.T., Albrechtsen A., Etter P.D., Johnson E.A., Orlando L., Chikhi L., Siegismund H.R., Heller R. 2018. A southern African origin and cryptic structure in the highly mobile plains zebra. Nat. Ecol. Evol. 2:491–498.

Pigot A.L., Tobias J.A. 2015. Dispersal and the transition to sympatry in vertebrates. Proc. R. Soc. Lond. B Biol. Sci. 282:20141929.

de Queiroz K. 2007. Species concepts and species delimitation. Syst. Biol. 56:879–886.

Quéméré E., Amelot X., Pierson J., Crouau-Roy B., Chikhi L. 2012. Genetic data suggest a natural prehuman origin of open habitats in northern Madagascar and question the deforestation narrative in this region. Proc. Natl. Acad. Sci. U.S.A. 109:13028–13033.

R Core Development Team. 2013. R: A Language and Environment for Statistical Computing. R Foundation for Statistical Computing. Vienna, Austria.

Radespiel U. 2016. Can behavioral ecology help to understand the divergent geographic range sizes of mouse lemurs? In: Lehman S., Radespiel U., Zimmermann E., editors. The Dwarf and Mouse Lemurs of Madagascar: Biology, Behavior and Conservation Biogeography of the Cheirogaleidae. Cambridge, UK: Cambridge University Press. p. 498–519.

Radespiel U., Lutermann H., Schmelting B., Zimmermann E. 2019. An empirical estimate of the generation time of mouse lemurs. Am. J. Primatol. 81:e23062.

Radespiel U., Olivieri G., Rasolofoson D.W., Rakotondratsimba G., Rakotonirainy O., Rasoloharijaona S., Randrianambinina B., Ratsimbazafy J.H., Ratelolahy F., Randriamboavonjy T., Rasolofoharivelo T., Craul M., Rakotozafy L., Randrianarison R.M. 2008. Exceptional diversity of mouse lemurs (*Microcebus* spp.) in the Makira region with the description of one new species. Am. J. Primatol. 70:1033–1046.

Radespiel U., Sarikaya Z., Zimmermann E., Bruford M.W. 2001. Sociogenetic structure in a free-living nocturnal primate population: sex-specific differences in the grey mouse lemur (*Microcebus murinus*). Behav. Ecol. Sociobiol. 50:493–502.

Rannala B., Yang Z. 2003. Bayes estimation of species divergence times and ancestral population sizes using DNA sequences from multiple loci. Genetics. 164:1645–1656.

Rannala B., Yang Z. 2013. Improved reversible jump algorithms for Bayesian species delimitation. Genetics. 194:245–253.

Rasolooarison R.M., Goodman S.M., Ganzhorn J.U. 2000. Taxonomic revision of mouse lemurs (*Microcebus*) in the western portions of Madagascar. Int. J. Primatol. 21:963–1019.

Rice W.R., Hostert E.E. 1993. Laboratory experiments on speciation: what have we learned in 40 years? Evolution. 47:1637–1653.

Evasoa M., Radespiel U., Hasiniaina A.F., Rasoloharijaona S., Randrianambinina B., Rakotondravony R., Zimmermann E. 2018. Variation in reproduction of the smallest-bodied primate radiation, the mouse lemurs (*Microcebus* spp.): A synopsis. Am. J. Primatol. 80:e22874.

Rodríguez W., Mazet O., Grusea S., Arredondo A., Corujo J.M., Boitard S., Chikhi L. 2018. The IICR and the non-stationary structured coalescent: towards demographic inference with arbitrary changes in population structure. Heredity. 121:663–678.

Salmona J., Heller R., Quéméré E., Chikhi L. 2017. Climate change and human colonization triggered habitat loss and fragmentation in Madagascar. Mol. Ecol. 26:5203–5222.

Salmona J., Olofsson J.K., Hong-Wa C., Razanatsoa J., Rakotonasolo F., Ralimanana H., Randriamboavonjy T., Suescun U., Vorontsova M.S., Besnard G. 2020. Late Miocene origin and recent population collapse of the Malagasy savanna olive tree (*Noronhia lowryi*). Biol. J. Linn. Soc. 129:227–243.

Schmelting B., Ehresmann P., Lutermann H., Randrianambinina B., Zimmermann, E. 2000. Reproduction of two sympatric mouse lemur species (*Microcebus murinus* and *M*. *ravelobensis*) in northwest Madagascar: first results of a long term study. In: Lourenço W.R., Goodman S.M. editors. Diversité et Endémisme à Madagascar. Paris: Société de Biogéographie. p. 165–175.

Schüßler D., Mantilla-Contreras J., Stadtmann R., Ratsimbazafy J.H., Radespiel U. 2020. Identification of crucial stepping stone habitats for biodiversity conservation in northeastern Madagascar using remote sensing and comparative predictive modeling. Biodivers. Conserv. https://doi.org/10.1007/s10531-020-01965-z.

Seiffert ER, Simons EL, Attia Y. 2003. Fossil evidence for an ancient divergence of lorises and galagos. Nature, 422:421–424.

Sgarlata G.M., Salmona J., Pors B.L., Rasolondraibe E., Jan F., Ralantoharijaona T., Rakotonanahary A., Randriamaroson J., Marques A.J., Aleixo-Pais I., Zoeten T. de, Ousseni D.S.A., Knoop S.B., Teixeira H., Gabillaud V., Miller A., Ibouroi M.T., Rasoloharijaona S., Zaonarivelo J.R., Andriaholinirina N.V., Chikhi L. 2019. Genetic and morphological diversity of mouse lemurs (*Microcebus* spp.) in northern Madagascar: The discovery of a putative new species? Am. J. Primatol. 81:e23070.

Skotte L., Korneliussen T.S., Albrechtsen A. 2013. Estimating individual admixture proportions from next generation sequencing data. Genetics. 195:693–702.

Stamatakis A. 2014. RAxML version 8: a tool for phylogenetic analysis and post-analysis of large phylogenies. Bioinformatics. 30:1312–1313.

Stange M, Sánchez-Villagra MR, Salzburger W, Matschiner M. 2018. Bayesian divergence-time estimation with genome-wide single-nucleotide polymorphism data of sea catfishes (Ariidae) supports Miocene closure of the Panamanian Isthmus. Syst Biol. 67:681–699.

Sukumaran J., Knowles L.L. 2017. Multispecies coalescent delimits structure, not species. Proc. Natl. Acad. Sci. U.S.A. 114:1607–1612.

Tattersall I. 2007. Madagascar’s lemurs: Cryptic diversity or taxonomic inflation? Evolutionary Anthropology: Issues, News, and Reviews. 16:12–23.

Tseng S.-P., Li S.-H., Hsieh C.-H., Wang H.-Y., Lin S.-M. 2014. Influence of gene flow on divergence dating-implications for the speciation history of *Takydromus* grass lizards. Mol. Ecol. 23:4770–4784.

Upham N.S., Esselstyn J.A., Jetz W. 2019. Inferring the mammal tree: Species-level sets of phylogenies for questions in ecology, evolution, and conservation. PLOS Biology 17(12): e3000494.

Uyeda J.C., Arnold S.J., Hohenlohe P.A., Mead L.S. 2009. Drift promotes speciation by sexual selection. Evolution. 63:583–594.

Vorontsova M.S., Besnard G., Forest F., Malakasi P., Moat J., Clayton W.D., Ficinski P., Savva G.M., Nanjarisoa O.P., Razanatsoa J., Randriatsara F.O., Kimeu J.M., Luke W.R.Q., Kayombo C., Linder H.P. 2016. Madagascar’s grasses and grasslands: anthropogenic or natural? Proc. R. Soc. Lond. B Biol. Sci. 283:20152262.

Wang K., Mathieson I., O’Connell J., Schiffels S. 2020. Tracking human population structure through time from whole genome sequences. PLoS Genetics. 16:e1008552.

Warmuth V.M., Ellegren H. 2019. Genotype-free estimation of allele frequencies reduces bias and improves demographic inference from RADSeq data. Mol. Ecol. Res. 19:586–596.

Weisrock D.W., Rasoloarison R.M., Fiorentino I., Ralison J.M., Goodman S.M., Kappeler P.M., Yoder A.D. 2010. Delimiting species without nuclear monophyly in Madagascar’s mouse lemurs. PLoS ONE. 5:e9883.

Wen D., Nakhleh L., Kubatko L. 2018. Coestimating reticulate phylogenies and gene trees from multilocus sequence data. Syst. Biol. 67:439–457.

Wilmé L., Goodman S.M., Ganzhorn J.U. 2006. Biogeographic evolution of Madagascar’s microendemic biota. Science. 312:1063–1065.

Yang Z., Rannala B. 2010. Bayesian species delimitation using multilocus sequence data. Proc. Natl. Acad. Sci. U.S.A. 107:9264–9269.

Yang Z., Yoder A.D. 2003. Comparison of likelihood and Bayesian methods for estimating divergence times using multiple gene Loci and calibration points, with application to a radiation of cute-looking mouse lemur species. Syst. Biol. 52:705–716.

Yoder A.D., Campbell C.R., Blanco M.B., Reis M. dos, Ganzhorn J.U., Goodman S.M., Hunnicutt K.E., Larsen P.A., Kappeler P.M., Rasoloarison R.M., Ralison J.M., Swofford D.L., Weisrock D.W. 2016. Geogenetic patterns in mouse lemurs (genus *Microcebus*) reveal the ghosts of Madagascar’s forests past. Proc. Natl. Acad. Sci. U.S.A. 113:8049–8056.

Yoder A.D., Rasoloarison R.M., Goodman S.M., Irwin J.A., Atsalis S., Ravosa M.J., Ganzhorn J.U. 2000. Remarkable species diversity in Malagasy mouse lemurs (primates, *Microcebus*). Proc. Natl. Acad. Sci. U.S.A. 97:11325–11330.

Zimin A.V., Marçais G., Puiu D., Roberts M., Salzberg S.L., Yorke J.A. 2013. The MaSuRCA genome assembler. Bioinformatics. 29:2669–2677.

Zimmermann E., Cepok S., Rakotoarison N., Zietemann V., Radespiel U. 1998. Sympatric mouse lemurs in north-west Madagascar: a new rufous mouse lemur species (*Microcebus ravelobensis*). Folia Primatol. 69:106–114.

Zimmermann E., Radespiel U. 2014. Species concepts, diversity, and evolution in primates: Lessons to be learned from mouse lemurs. Evolutionary Anthropology: Issues, News, and Reviews. 23:11–14.

Zohdy S., Gerber B.D., Tecot S., Blanco M.B., Winchester J.M., Wright P.C., Jernvall J. 2014. Teeth, sex, and testosterone: aging in the world’s smallest primate. PLoS ONE. 9:e109528.

