## Supplemental Information for "Cryptic Patterns of Speciation in Cryptic Primates: Microendemic Mouse Lemurs and the Multispecies Coalescent"

**Table of Contents**

1.1 Sampling	2

1.2 RAD sequencing libraries preparation protocols	3

1.3 mtDNA amplification and sequencing	4

1.4 M. sp. #3 genome annotation	6

1.5 RADseq datasets	6

1.6 RADseq genotyping and filtering	7

1.7 Conversion to full-sequence fasta files	11

1.8 Phylogenetic Analyses	11

1.9 SNAPP	12

1.10 PCA and clustering methods	13

1.11 Priors for SNAPP and BPP	13

1.12 Guided Species Delimitation with BPP	14

1.13 Isolation-by-distance analysis	14

1.14 Inference of gene flow with G-PhoCS	14

1.15 D-statistics	15

1.16 Divergence time estimation with BPP	15

1.17 Inference of effective population size with MSMC	15

1.18 Genetic diversity and differentiation	16

2 Supplementary results	17

2.1 Phylogenetic analyses	17

2.2 Species Delimitation: Guided Species Delimitation with BPP	18

2.3 Gene flow	18

2.4 Genetic diversity	19

3 Supplementary Figures	20

4 Supplementary Tables	40

5 Supplementary References	47

#### 1.1 Sampling

*Microcebus* spp. were sampled in the protected areas of Marojejy National Park (NP) and Anjanaharibe-Sud Special Reserve, within the Antainambalana-Voloina inter-river system (IRS) around the village of Anjiahely (Makira region; where the holotype of *M. macarthurii* was obtained; Radespiel et al. 2008), in the fragmented forests of the Mananara-Fahambahy IRS at the village of Ambavala (Schüßler et al. 2018), in the Mananara-Anve IRS within the Mananara-Nord NP (Ivontaka-Sud section), around Antanambe village in the vicinity of the Mananara-Nord NP, and in the Ambodiriana community protected area at Manmpana within the Anve-Simianna IRS (Miller et al. 2015, 2018*;* Fig. 1).

*Microcebus* were caught either with Sherman Life traps or directly by hand during nocturnal surveys (Radespiel et al. 2008). Ear biopsies (~2 mm²) were collected and stored in Queen’s lysis buﬀer (Seutin et al. 1991). Animals were released unharmed within 24 hours at their exact place of capture.

DNA was extracted for further analyses from tissue samples (Tab. S1) using a QIAGEN DNeasy Blood & Tissue Kit modified protocol described in Sgarlata et al. (2018) or by a standardized phenol/chloroform extraction technique (Maniatis et al. 1982). We quantified DNA using Qubit ® technology (Invitrogen).

#### 1.2 RAD sequencing libraries preparation protocols

Three different but compatible protocols for library generation were used.

Protocol 1: Oregon

We prepared restriction-site associated DNA (RAD) sequencing libraries using 40-100 ng of genomic DNA from each sample and one negative control made up of water following the protocol described in . Brieﬂy, samples were digested with SbfI (New England Biolabs) before ligation of the P1 adapter (Etter et al. 2011; genomic Resources Development Consortium et al. 2015). We pooled 24 or 48 samples each in sub-libraries, and used proportionally more of the reaction if less than 100 ng of DNA was used in the digestion, before shearing in a Bioruptor for 6 min to obtain fragments with an average size of 500 bp. Following end repair, we ligated the P2 adapter before 14 cycles of PCR amplification . We normalized the library fragment size using AMPure XP beads (Agencourt), pooled the sub-libraries based on yield and sequenced libraries on an Illumina HiSeq 2000 lanes (single end, 100pb, 96 individuals / 2 lanes) at the University of Oregon Core facility.

Protocol 2: Toulouse

We prepared restriction-site associated DNA (RAD) sequencing libraries using 40-200 ng of genomic DNA from each sample and one negative control made up of water following the protocol described in . Brieﬂy, samples were digested with SbfI (New England Biolabs) before P1 adapter ligation . After adapter ligation, we pooled 10 µl of each 24 samples in sub-libraries, before shearing them in a Covaris^®^ M220 for 45 sec to obtain fragments with an average size of 500 bp. Following end repair, we conducted size selection using AMPure XP beads (Agencourt). After 3’adenylation, we ligated the P2 adapter. The libraries were finally enriched with 10 cycles of PCR . Library quality was verified using Fragment Analyzer^®^ (Advanced Analytical) and quantified using qPCR (QuantStudio6^®^, Applied Biosystems). We made equimolar pooling with the sub-libraries and sequenced them on an Illumina HiSeq 3000 (paired end, 150pb, 96 individuals / lane) at the Gentoul Sequencing platform facility (Toulouse, France).

Protocol 3: Idaho

We prepared restriction-site associated DNA (RAD) sequencing libraries using 50 ng of genomic DNA from each sample following the protocol of Ali et al. (2016). Briefly, samples were digested with SbfI (New England Biolabs), followed by ligation with custom biotinylated adaptors containing 8bp barcodes unique to each sample. We pooled 48 samples in a single library along with 4 technical replicates and sheared to an average fragment size of 400bp using a Covaris M220. RAD fragments were enriched with a streptavidin bead pull-down and prepared as a sequencing library using a NEBNext Ultra DNA Library Prep Kit (New England Biolabs). The final library was sequenced in a single lane of paired-end 150bp sequencing on an Illumina HiSeq4000 (paired end, 150bp, 48-96 individuals / lane) at the Vincent J. Coates genomic Sequencing Laboratory, University of California Berkeley, or at the Duke Center for genomics and Computational Biology sequencing facility.

#### 1.3 mtDNA amplification and sequencing

Protocol 1: Duke

DNA was extracted with the QIAGEN DNeasy tissue kit. A 1140 bp fragment of Cyt B was amplified with the forward primer 5’-TGA-YTA-ATG-AYA-TGA-AAA-AYC-ATC-GTT-G-3’ and the reverse primer 5’-TCT-CCA-TTT-CTG-GTT-TAC-AAG-ACC-A-3’. A 684 bp fragment of COII was amplified using primers L7553 and H8320 (Adkins and Honeycutt 1994). Both PCR reactions used the following thermocycler recipe: Denature for 2 minutes at 96 °C followed by 24 cycles of 30 s at 96 °C, 20 s at 50 °C, and 4 minutes at 60 °C. Sequencing reactions used BigDye Terminator V1.1 per manufacturer specifications and sequencing was performed with an Applied Biosystems 3730xl. Sequencing was done at the Duke Sequencing Core resource. Chromatograms were assembled into consensus sequences with Geneious (Kearse et al. 2012).

Protocol 2: Hannover

DNA was extracted using the QIAGEN DNeasy Blood & Tissue Kit https://www.qiagen.com/us/) following the provided protocol or using a standardized Phenol/Chloroform extraction (Maniatis et al. 1982). Three different mitochondrial loci (*cytb*, COII, d-loop) were amplified using primers given in Olivieri et al. (2007) and in addition CytBmac_F/CytBmac_R Primer (CytBmac_F: 5'-CAGCACCATCCAACATTTCTT, CytBmac_R: 5'-GCTACGAAAGGGTATTCAACG) or L14724 and H15915 (Irwin et al. 1991) to sequence a longer part of Cyt B . Polymerase chain reaction (PCR) followed the protocol by Guschanski et al. (2007) for COII/D-loop and 45 cycles of 30 s at 95 °C, 30 s at 54 °C, and 30 s at 72 °C for Cyt B . PCR products were cleaned after successful amplification (checked on a 1.5% agarose gel containing 1.3 x 10^-4^ mg/ml ethidium bromide) using the standard protocols of the Invisorb® Spin PCRapid kit (Invitek) or the MSB Spin PCRapace kit (Stratec). Resulting PCR products were sent to Macrogen Ltd. (Seoul, South Korea[http://dna.macrogen.com](http://dna.macrogen.com/)) for sequencing on an ABI 3730XL automatic DNA sequencer or to GATC Biotech (https://www.gatc-biotech.com/en/) for Sanger sequencing. All sequences were then analyzed, edited and aligned using Sequencher TM 4.0.5 (Gene Codes, Ann Arbor, MI, US) for all samples collected before 2014 and with Geneious 11.1.2 (Kearse et al. 2012<https://www.geneious.com/>) for all samples collected after 2014. Sequences were first blasted to confirm species identity. DNA samples are stored at the University of Veterinary Medicine Hanver. Not all loci are available for all samples; Table S1 provides an overview on the sequences included in this study.

#### 1.4 M. sp. #3 genome annotation

Gene annotations were generated *in silico* with MAKER v2.31.9 (Cantarel et al. 2008) for individual MBB030 (Tab. S1). First, we trained an *ab initio* gene predictor, SNAP (Korf 2004), with a *de novo* transcriptome assembly from *Microcebus murinus*. We used publicly available RNAseq data (Peng et al. 2014) that included technical replicates from multiple tissue types to assemble a single transcriptome with TRINITY v2.2.0 (Haas et al. 2013). For an independently trained gene predictor, we optimized the AUGUSTUS v3.3 (Stanke et al. 2006) human parameters with the Microcebus *murinus* 3.0 assembly (https://www.ncbi.nlm.nih.gov/genome/?term=Microcebus+murinus; last accessed 8 November 2017) using BUSCO v3.0.2 (Simão et al. 2015). Final annotations were generated with a single MAKER run using a combination of transcriptome and homologous protein alignment as well as gene predictions. Assembly and annotation completeness was evaluated with BUSCO v3.0.2 (Simão et al. 2015).

To ensure that only autosomal data was used for analyses, potential mitochondrial or X-linked scaffolds were removed from the assembly by performing a BLASTN (Altschul et al. 1990) search. For mitochondrial screening, we blasted against available *Cheirogaleidae* mitochondrial genomes from NCBI. Scaffolds with 90% or more sequence identity over 50% of the scaffold length compared to any *Cheirogaleidae* mitochondrial genome were removed (32,511 base pairs in total). For X-chromosome screening, we blasted against the *Microcebus murinus* superscaffold NC_033692.1, which is designated as the X-chromosome. To be conservative, we removed scaffolds with 85% sequence similarity over 10% of the scaffold length; 76,444,357 base pairs across 3,538 scaffolds. Although the *M. murinus* X-chromosome is nearly twice that in length, these genomes are repeat-rich (Larsen et al. 2017) and our single Illumina genome was likely unable to assemble repetitive regions better captured by long-read and structural data.

#### 1.5 RADseq datasets

In Table S1, we specify which samples were used for different subsets of the data. Phylogenetic, clustering, and PCA analyses were performed with all individuals that passed QC, as specified in the “SNP calling” column in Table S1. For computational reasons, multispecies coalescent methods were ran on subsets of the individuals. SNAPP analyses were done in parallel using a 22-individual dataset (“SNAPP-22” column in Table S1) and a 12-individual dataset (“SNAPP-12, BPP, G-PhoCS-all” column in Table S1). This same 12-individual SNAPP dataset was also used for BPP and G-PhoCS analyses, but with full-sequence fasta files rather than SNPs (see “Conversion to full-sequence fasta files” section below). Additionally, G-PhoCS was run with only the lineages in Clade I and M. lehilahytsara, as specified in the “G-PhoCS-cladeI” column in Table S1.

#### 1.6 RADseq genotyping and filtering

Read processing

Raw reads were demultiplexed in Stacks v2.0b (Catchen et al. 2013) using default parameters, and quality filtered using Trimmomatic (Bolger et al. 2014) with the following parameters: Leading: 3, Trailing: 3, Slidingwindow: 4:15, Minlen: 60. Using BWA MEM v0.7.15 (Li 2013), reads were then aligned to autosomal *M.* sp. #3 scaffolds and to the *M. murinus* mitochondrial genome (Lecompte et al. 2015). Alignments were sorted and PCR duplicates removed in the “rmdup” program in SAMtools v1.6 (Li et al. 2009).

GATK
For compatibility with GATK v. 4.0.7.0, which is designed to deal with only a small number of scaffolds, we first used ScaffoldStitcher (<https://bitbucket.org/dholab/scaffoldstitcher/src/master/https://bitbucket.org/dholab/scaffoldstitcher>) to "stitch" the *M*. sp. #3 reference genome into "superscaffolds" of up to 100 Mbp, with 1 kbp of Ns between each actual scaffold to prevent mapping errors[. Next, fastq files were mapped to the stitched genme using BWA MEM and bam files were processed prior to gentyping (see Main Text for details).](https://bitbucket.org/dholab/scaffoldstitcher)

We used GATK‘s “HaplotypeCaller” tool to produce GVCF files for each sample, which were then merged to multi-sample, single-scaffold "GenomicsDB Workspaces" using GATK‘s “GenmicsDBImport” tool (this merging step is necessary because in GATK v4 since the joint genotyping tool n longer accepts multiple GVCF files), and these Workspaces were then used as input for GATK‘s “GenotypeGVCFs” tool, which was run using the "--use-new-qual-calculator" option.

VCF files were filtered according to recommendations from O‘Leary et al. (2018), following their "FS6” filtering steps (see Table 2 in O‘Leary et al. 2018). First, we selected only SNPs (i.e., discarding structural variants) using GATK‘s SelectVariants tool with option "-selectType SNP", and annotated the VCF file with the Allele Balance statistic using GATK’s “VariantAnnotator” tool with option “-A AlleleBalance”. The FS6 filtering involved removing the following data in the following order:

1. Genotypes with a minimum per-sample depth of 5 (using "--minDP 5" in vcftools); these are set to “missing”.
2. Sites with an across-sample genotype quality (Qual) lower than 20 (using "--minQual 20" in vcftools).
3. Sites with an across-sample mean depth lower than 15 (using "--min-meanDP 15" in vcftools).
4. Sites with a minor allele count (MAC) below 3 ***or*** 1 (using "--mac 3" or " --mac 1" in vcftools). Thus, this step represents a fork in the pipeline, with parallel filtering occurring for files with “MAC=1” and with “MAC=3” hereafter.
5. Sites/individuals with specified amounts of missing data. A progressive series of filtering steps for missing data was performed, alternating between filtering at the site level (wherein sites with more than the specified percentage of missing data are removed) and filtering at the individual level (wherein sites with more than the specified percentage of missing data are removed). Site-filtering was done using "--max-missing [threshold]" in vcftools, whereas individual-filtering first required the computation of per-individual statistics on missing data (using "--missing-indv" in vcftools), followed by the removal of individuals exceeding the threshold using "--remove [individual_ID]" in vcftools. (note that this procedure is the same as in O‘Leary et al.’s FS6, but the following notation is slightly different.) Filtering steps:

a) site: 50%

b) individual: 90%

c) site: 40%

d) individual: 70%

e) site: 30%

f) individual: 50%

6. Sites with more than two alleles. Note that this step is not present in O‘Leary‘s FS6 filter.
7. Sites not passing any of the following filters based on statistics in the “INFO” field of the VCF file.

a) Allele balance (ABHet < 0.2 or ABHet > 0.8)

b) Mapping quality ratio of reference vs. alternative allele (MQRankSum < -12.5)

c) Strandedness of reference vs. alternative allele (FS > 60.0)

d) Quality-by-depth ratio (QD < 2.0)

e) Absolute mapping quality (MQ < 40.0)

f) Read position of reference vs. alternative allele (ReadPosRankSum < -8)

These sites were set to “FILTER” status using GATK‘s “VariantFiltration” tool followed by removal using "--remove-filtered-all" in Vcftools. The threshold values used here are based on GATK guidelines (<https://gatkforums.broadinstitute.org/gatk/discussion/2806/howto-apply-hard-filters-to-a-call-set>), except for item a), the allele balance filter, which is not present in the GATK recommendations. We used the GATK guidelines here since O‘Leary et al. (2018) were not always explicit with respect to thresholds and/or performed filtering with custom procedures rather than “VariantFiltration” parameters. Additional minor differences with O‘Leary’s et al. (2018) framework are that we also included filtering based on absolute mapping score (item e) above) and read position (item f) above), and that we removed non-properly paired reads during bam-file processing steps rather than in the VCF.

8. Sites with a depth larger than the mean depth plus twice the standard deviation in depth. The maximum depth was first computed and sites with excessive depth were then removed using “–max-meanDP” in Vcftools.
9. A final round of filtering sites and individuals with excessive missing data (as in step 5, see above) were removed using:

a) individual: 25%

b) site: 5%

When applied to the unfiltered VCF with all 65 samples included in this study, this filtering procedure resulted in filtered VCF files with 53 individuals and 61,561 and 34,409 SNPs for MAC1 and MAC3, respectively. For analyses on subsets of the data, such as the 6-species and 4-species datasets used in the G-PhoCS analysis, we started by subsetting the *raw* VCF with the focal samples, and then applying the filters as listed above. Except for analyses using full-sequence fasta files (BPP and G-PhoCS), all the results presented are based on VCF files with a MAC of 3 (parallel analyses were done using files with a MAC of 1, and consistently returned highly similar results – not shown).

ANGSD
We first estimated locus depth as the F1 read starting position (sbf1 cutting site) depth using the “samtools-view” command (Li et al. 2009) in R (R Core Team 2013). We then estimated genotype likelihoods (“-GL 1”) using only loci with (1) a total depth of at least twice the number of individuals (“setMinDepth”), (2) a total depth of at most (“-setMaxDepth”) the sum of the 0.95 quantile of the depth distribution of considered individuals, and (3) an individual depth of at most (“-setMaxDepthInd”) the maximum 0.95 quantile of considered individuals. This strategy allows discarding loci present only in a very low number of individuals, or over-covered loci that are likely to be repeated or paralog regions (Pedersen et al. 2018; Heller et al. submitted). In addition, we considered only bases with a minimum quality (“-minQ”) of 20, reads mapping uniquely (“-uniqueOnly”) with a minimum quality (“-minMapQ”) of 30 and associated with their pair (“-only_proper_pair”; for PE only), we only kept biallelic variants (“-skipTriallelic”) with a probability (“-SNP_pval”) below 1e-6 and a minor allele frequency (“-minMaf”; MAF) of at least 0.05.

Variant calling with ANGSD (“-doGen -4”) applied the filtering thresholds specified above for genotype likelihood estimations. The resulting .plink format file (“-doPlink 2”) was subsequently filtered in PLINK (Purcell et al. 2007) to remove samples (“--mind; >75%”), and loci “(--gen; >70%”) with a high proportion of missing data, a MAF below 0.05, and keeping 1 locus per 10kb (“--bp-space”) to limit linkage disequilibrium among loci. Finally, we called haplotypes from the unique sbf1 mtDNA loci using ANGSD (Nielsen et al. 2012; Korneliussen et al. 2014).

#### 1.7 Conversion to full-sequence fasta files

To produce full-sequence fasta files from the VCF files, we used the following procedure: First, we ran the GATK v3.8 tool “FastaAlternateReferenceMaker” for each sample, producing a single-sample whole-genome fasta file based on the reference genome but replacing sites that were called as non-reference alleles in the *unfiltered* VCF file. Then, the following classes of bases were masked: (1) non-reference bases that did not pass filtering (see RADseq genotyping and filtering section above); (2) sites that were classified as non-callable using GATK v3.8 “CallableLoci” (applying a minimum depth of 3). Next, “loci” were extracted by (1) defining loci at an individual level as stretches of at least 25 consecutive called (non-N) bases or multiple such stretches separated by at most 10 consecutive Ns; (2) intersecting these stretches across individuals, requiring at least 1 bp of overlap, and trimming from both ends any bases with fewer than 80% of focal individuals represented; (3) filtering the resulting loci to include only loci (a) with at least 90% of focal individuals represented, (b) of at least 100 bp length, (c) with at most 10% missing data.

For the main, SNAPP-12 dataset (Table S1), we ended up with 15,267 loci, 1,822 of which were invariant. The mean locus length was 231 bp, and the mean number of variable and parsimony-informative sites was 4.7 and 3.29, respectively. The mean amount of missing data was 5.53%.

#### 1.8 Phylogenetic Analyses

For maximum likelihood (ML) analyses, we separately analyzed mtDNA and nDNA. We concatenated mtDNA loci from RADseq data with Sanger sequences of cytB, COII and d-loop, and inferred phylogenetic relationships in RAxML using the GTR substitution model with CAT rate heterogeneity. We inferred a phylogeny from nuclear RADseq SNPs (dataset VCa1, Tab. S6) in RAxML using the GTR substitution model with GAMMA rate heterogeneity, applying an acquisition bias correction for binary SNP data (Lewis 2001; Leaché et al. 2015). Both analyses were conducted with 200 rapid bootstraps followed by a 10-step ML search optimization.

SVDquartets uses topological invariants for quartets to estimate a single species tree, but unlike the ML analyses, allows for topological discordance among sites. We ran SVDquartets in PAUP* v4a163 (Swofford 2003) and randomly sampled 5x10^6^ quartets with 100 bootstraps. We ran two analyses, one that assigns all individuals to their species *a priori*, and one that treats all individuals as a separate lineage. ML and SVDquartets trees were plotted using the ape package in R (R Core Development Team 2013; Paradis et al. 2018).

#### 1.9 SNAPP

SNAPP was run using BEAST2 v2.4.8 (Bouckaert et al. 2014). We applied informed and proper priors (Supplementary Material) to sample model parameters so we could estimate marginal likelihoods (Gelman and Meng 1998) for downstream species delimitation analyses while using the conditional likelihoods for SNP data (Leaché et al. 2015). To explore the effects of the number of individuals included in analyses and potential geographic sampling gaps, we analyzed a data set of 12 individuals that used 2 samples per species and a data set of 22 individuals that used 4 samples per ingroup species. The 2 samples per species in the 12 individual dataset typically spanned the two farthest sampling locations, and the 4 samples per species in the 22 individual analyses filled in sites in between, when available. All analyses used *M. murinus* samples as the outgroup. The 12-individual analyses ran for 10 million generations after a burn-in of 500,000 generations. We sampled every 500 generations to generate posteriors of 20,000 samples. The 22-individual analyses required significantly more computation; these analyses only ran for 5 million generations to generate posteriors of 10,000 samples. Two independent chains were run for both analyses to check convergence. Effective sample sizes and mixing were evaluated with TRACER v1.7 (Rambaut et al. 2018). SNAPP trees were plotted with the ggTree v1.15.1 package in R (Yu et al. 2017). Marginal likelihood estimation used stepping-stone sampling (Xie et al. 2011) with 20 steps. Marginal likelihoods were used to test the species delimitations between *M.* sp #3 and *M. macathurii* as well as *M. lehilahytsara* and *M. mittermeieri*. Specifically, we estimated marginal likelihoods for the following three hypotheses:

1) (*M. murinus*, ((*M. simmonsi*, (*M. lehilahytsara*, *M. mittermeieri*)),(*M.* sp. #3, *M. macarthurii*)));

2) (*M. murinus*, ((*M. simmonsi*, *M. lehilahytsara* and *M. mittermeieri*),(*M.* sp. #3, *M. macarthurii*)));

3) (*M. murinus*, ((*M. simmonsi*, (*M. lehilahytsara*, *M. mittermeieri*)),*M.* sp. #3 and *M. macarthurii*));

XML files for analyses are available in Dryad.

#### 1.10 PCA and clustering methods

PCAs of genotype likelihoods were performed using default parametrization, and glPCA was run using “scale” set to “TRUE” and retaining the first four principal components. NgsAdmix and ADMIXTURE admixture proportions were inferred with the number of clusters (*K*) ranging from two to the number of hypothesized taxa plus two (i.e., eight for the full dataset, and four for the subset). NgsAdmix was run for 20 iterations per *K* value and a stringent tolerance of 10^-6^ for convergence. We compared *K* values using the ΔK procedure (Evann et al. 2005). ADMIXTURE was run using the default optimization method and the default termination criterion, which is to stop when the log-likelihood increases by less than 10^-4^ between iterations, and *K* values were compared using the cross-validation method (“--cv” flag in ADMIXTURE) and by comparing likelihoods across runs. NgsAdmix results were plotted geographically using the *RgoogleMap* and *scatterpie* R packages (Loecher & Ropkins 2015).

#### 1.11 Priors for SNAPP and BPP

SNAPP
For the coalescent speciation rate λ, we assigned a prior distribution of Γ(3,0.00414), which provides a diffuse distribution around a mean of 725 using the expectations of the number of species for a Yule prior (Drummond and Bouckart 2015) and a tree length (0.002) approximated from a previous coalescent analysis of mouse lemurs (Yoder et al. 2016). The ancestral nucleotide diversity estimates, θ, were assigned independent Γ(3,3000) distributions, given that empirical interspecific θ estimates for mouse lemurs range from 0.0007 to 0.02 across a small sampling of nuclear genes (Blair et al. 2014).

BPP
To generate diffuse priors while placing some biologically realistic constraints on parameter space for uninformative loci, θ was assigned a prior inverse gamma distribution with α = 2 and β = 0.0017. The coalescent tree length was distributed as inverse gamma with α = 3 and β = 0.0041. Priors were visually inspected using the *invgamma* R package (Kahle and Stamey 2017).

#### 1.12 **Guided Species Delimitation with BPP**

Although BPP is capable of unguided species delimitation, we were confident in our species tree topology and implemented several guided analyses to explore sensitivity of results to algorithm and prior choice. Guided species delimitation can use two different algorithms: 1) ALG 0 which relies only on a single tuning parameter ε to determine node height and ancestral nucleotide diversity proposals after splitting and merging of species, and 2) ALG 1 which uses a gamma distribution with shape α and a tuning parameter *m* that determines the mean given the previous θ state. We performed analyses using both algorithms to ensure that support for all species was not the result of inefficient mixing. Analyses used θ and τ priors as specified above, but we recognize that our θ prior make strong assumptions that may bias analyses in cases where data is not informative. A small θ would also imply a faster rate of coalescence and the splitting of species, perhaps providing false confidence in species tree bipartitions. Therefore, we replicated analyses with large priors on θ; the mean was 100 times greater than our original prior mean, an inverse gamma prior with α = 3 and β = 0.17. Large ancestral θ estimates would require more generation for lineages to coalesce, potentially weakening support for species hypotheses.

#### 1.13 Isolation-by-distance analysis

The main VCF file produced by GATK-genotyping was read into R using the read.vcfR() function and then converted to a “genlight” object using the vcfR2genlight() function, both from the package vcfR 1.10.0 (Knaus and Grünwald 2017). GPS coordinates for each site were added to the “@other$latlong” slot of the genlight object, such that gl.ibd() could calculate the genetic and geographic distance matrices directly from the provided genlight object. The isolation-by-distance test was then performed on the distance matrix (using the natural log of the Euclidean distance in meters) against the population-based pairwise FST/1-FST matrix.

#### 1.14 Inference of gene flow with G-PhoCS

In G-PhoCS, migration (gene flow) is modeled using one or more discrete unidirectional migration bands between a pair of extant or ancestral lineages that overlap in time. However, simultaneously estimating migration rates for all possible migration bands (over 40 in total for the 6-species dataset) is not feasible (Gronau et al. 2011). Moreover, comparing likelihoods across different models is currently not possible in G-PhoCS (pers. com. I. Gronau). Therefore, we started by testing reciprocal pairs of migration bands (e.g., a model with migration from A→B and B→A), and next combined significant migration bands in a final model with multiple migration bands. The proportion of migrants per generation is calculated by multiplying m by the per-generation mutation rate (μ).

#### 1.15 D-statistics

Significance of D-values was determined using the default Z-value reported by qpDstat, which is determined by weighted block jackknifing and is conservative for RADseq data given that linkage disequilibrium (LD) is, on average, expected to be lower across a pair of RADseq SNPs than across a pair of SNPs derived from whole-genome sequencing (Patterson et al. 2012; Kim et al. 2018).

#### 1.16 Divergence time estimation with BPP

We ran four chains of 1.2 million samples, sampling every 100 generations. The first 5,000 samples were discarded as burn-in and the other 7,000 samples retained as the posterior distributions. Convergence across chains was checked by visual inspection of posteriors and with potential scale reduction factors in the R package CODA (Plummer et al. 2006). Final parameter estimates were obtained by combining all four chains for a posterior of 28,000 samples. Coalescent node heights (τ) were converted into absolute time by assuming time in years = (τ * generation time in years)/ mutation rate. Estimates of θ were converted to effective population sizes by *N_e_* = θ/(4 * mutation rate). Marginal priors were obtained from BPP by setting usedata = 0 in the control file.

#### 1.17 Inference of effective population size with MSMC

We used whole-genme sequencing data for one *M.* sp. #3 individual (MBB030, Tab. S1) and one *M. mittermeieri* individual (reads available at the SRA at [SRR8456525](https://dataview.ncbi.nlm.nih.gov/object/SRR8456525" \l "_blank)) to infer population sizes through time with MSMC. When using a single individual, unphased genme-wide genotypes are used to create MSMC input files. Since MSMC relies on estimates of local heterozygosity, which are easily underestimated for low- to medium-coverage data, relatively high coverage is needed for accurate inference. Our coverage of 47.84x (*M.* sp. #3) and 48.40x (M. mittermeieri) are both well above the lower threshold of 18x recommended by Nadachowska-Brzyska et al. (2016). Reads were mapped to the genme assembly for *M. murinus* (v3.0, https://www.ncbi.nlm.nih.gov/genme/777?genme_assembly_id=308207; Larsen et al. 2017) rather than the *M.* sp. #3 assembly generated in this study, given that the assembly for *M. murinus* is considerably more contiguous (N50: 210 kb) and MSMC benefits from long stretches of contiguous sequence. Sequences were mapped using BWA MEM 0.7.15 (Li 2013), and bam files were sorted and deduplicated using the Picard Tools (version 2.13.2, http://broadinstitute.github.io/picard) “SortSam” and “MarkDuplicates” commands, respectively. Next, genotypes were called for scaffolds greater than 1 Mb from bam files using the Samtools v1.6 “mpileup” program, following the MSMC documentation (https://github.com/stschiff/msmc-tools), with the following settings: minimum mapping quality of 20 (flag “-q 20”), minimum genotype quality of 20 (flag “-Q 20”), and a coefficient for downgrading mapping quality for reads that contain excessive mismatches (flag “-C 50”). In accordance with recommendations in the MSMC documentation, we further masked sites with less than half or more than double the mean coverage. The MSMC input file was created from a VCF file using the “generate_multihetsep.py” script provided with the MSMC distribution. The default time period scheme of “10×1 + 15×2” was used. To convert the scaled population size θ to Ne with θ = 4 × Ne × μ, we used a generation time of 3.75 years and an estimate of the mutation rate as described in the methods.

#### 1.18 Genetic diversity and differentiation

For comparison with clustering results, heterozygosity and F_ST_ were estimated with the R packages adegenet v2.1.1 and hierfstat (Goudet 2005), respectively, using variable sites inferred from ANGSD. Heterozygosity was estimated from ANGS GLs as the proportion of variable sites. Population based observed (*H*_o_) and expected heterozygosity (*H*_e_) as well as among populations measure of divergence (*F*_ST_, Weir and Cockerham 1984) were estimated using the adegenet v2.1.1 and hierfstat R packages (Goudet 2005; Jombart 2008), using variable sites only.

### 2 Supplementary results

#### 2.1 Phylogenetic analyses

RAxML

The resulting data matrix had 28,738 SNPs with and an average of 15% missing data (VCa1 dataset; Table S2). ML topologies from both mtDNA and nDNA data from RADseq recovered paraphyly among *M. lehilahytsara* and *M. mittermeieri*. Additionally, the mtDNA tree implied the presence of mtDNA from *M. macarthurii* in the *M.* sp. #3 lineage. Although bootstrap support is typically weak in the mtDNA tree with many short branches subtending the *M. macarthurii* individuals with *M.* sp. #3 DNA, suggested a small number of sites unite these lineages together in the mtDNA tree.

SVDquartets

Individual-level and species-level analyses with SVDquartets recovered strongly supported topologies mostly congruent with the maximum likelihood topologies shown in Fig. 3. All bipartitions in the species-level SVDquartets analysis were recovered with a bootstrap support of 100%.

SNAPP

For the twelve-individual analyses, a total of biallelic 20,624 GATK-called SNPs went into the analysis, of which 19,691 were retained after SNAPP's filtering. Using two independent runs, we determined that SNAPP had largely converged and although there was some uncertainty in a couple of θ parameters that may have benefited from more sampling, it is unlikely that uncertainty in coalescent branch length distributions and posterior node heights would have reduced any further (Fig. S14; Tab. S7). Median node heights between runs were nearly identical after collecting 20,000 posterior tree samples (Fig. S14). There was n uncertainty in the species tree topology, with perfect posterior probabilities at all nodes in both the twelve-individual and 22-individual analyses (Fig. S1). 22-individual analyses also converged based on analyses of posterior distributions of sampled parameters and median node heights (Tab. S7; Fig. S15). However, 22-individual analyses estimated branch lengths to be an order of magnitude shorter, although proportional compared to the twelve-individual analyses (Fig. S1).

#### 2.2 Species Delimitation: Guided Species Delimitation with BPP

As an alternative to Bayes factors with SNAPP, we used guided delimitation (Yang and Rannala 2010) with BPP. Guided delimitation estimates the posterior probabilities of different species delimitation models while marginalizing over MSC parameters for a fixed species tree topology. Different species delimitation hypotheses are explored with rjMCMC by collapsing internal nodes on a fixed guide tree, greatly reducing the space being explored by the rjMCMC algorithm to only 8 delimitation hypotheses. Unsurprisingly, guided analyses with algorithms 0 and 1 recovered all species tree splits with posterior probabilities of 1. We increased the priors on θ to check if our delimitations were sensitive to assumptions about nucleotide diversity and ancestral population sizes in mouse lemurs – a larger θ should make collapsing nodes more likely in the absence of information-rich data. The larger, likely mis-specified, priors did not reduce posterior probabilities for the species delimitation hypothesis that does not collapse any of our assigned lineages . Therefore, our species delimitation analyses are likely robust to reasonable prior mis-specifications.

#### 2.3 Gene flow

Tests for gene flow between non-sister species using D-statistics

The D-statistic was first calculated for all possible configurations in which gene flow between ingroup non-sister species could be evaluated. While none of these comparisons were significant using a conservative threshold, negative values of D with Z-scores larger than 2 were obtained for all four comparisons with *M. lehilahytsara / M. mittermeieri* as P1, M. *simmonsi* as P2, and *M. macarthurii / M*. sp. #3 as P3, suggesting a slight excess of shared derived alleles and therefore gene flow between the *M. lehilahytsara / M. mittermeieri* clade and the *M. macarthurii / M.* sp. #3 clade (Fig S16).

Inference of gene flow between the *M. macarthurii / M.* sp. #3 clade and *M. lehilahytsara*

In the 3-species model, G-PhoCS also inferred gene flow from the *M. macarthurii / M.* sp. #3 clade to *M. lehilahytsara,* but was not able to pinpoint the source of gene flow (which could be either *M. macarthurii*, northern *M.* sp. #3, southern *M.* sp. #3, ancestral *M.* sp. #3, or the ancestor of *M. macarthurii* and *M*. sp. #3) in models with multiple migration bands combined (Fig. S10A). Because *M. lehilahytsara* currently occurs sympatrically with northern *M.* sp. #3 only, we included a migration band from northern *M*. sp. #3 in the final model with multiple migration bands.

#### 2.4 Genetic diversity

Individual-based diversity was highly variable among sampling sites (Fig. S13). *M. lehilahytsara* exhibits the highest diversity, followed by *M. mittermeieri*, M. sp. #3, *M. simmonsi* and *M. macarthurii*. Except for *M. simmonsi* and *M. sp. #3*, these results are reflected in the estimates of effective populations sizes (Fig. 7). Species that are occur in sympatry, *M. sp. #3* and *M. lehilahytsara* in Ambalava, and *M. macarthurii* and *M. mittermeieri* in Anjiahely, show clearly distinct ranges of genetic diversity (Fig. S13).

### 3 Supplementary Figures

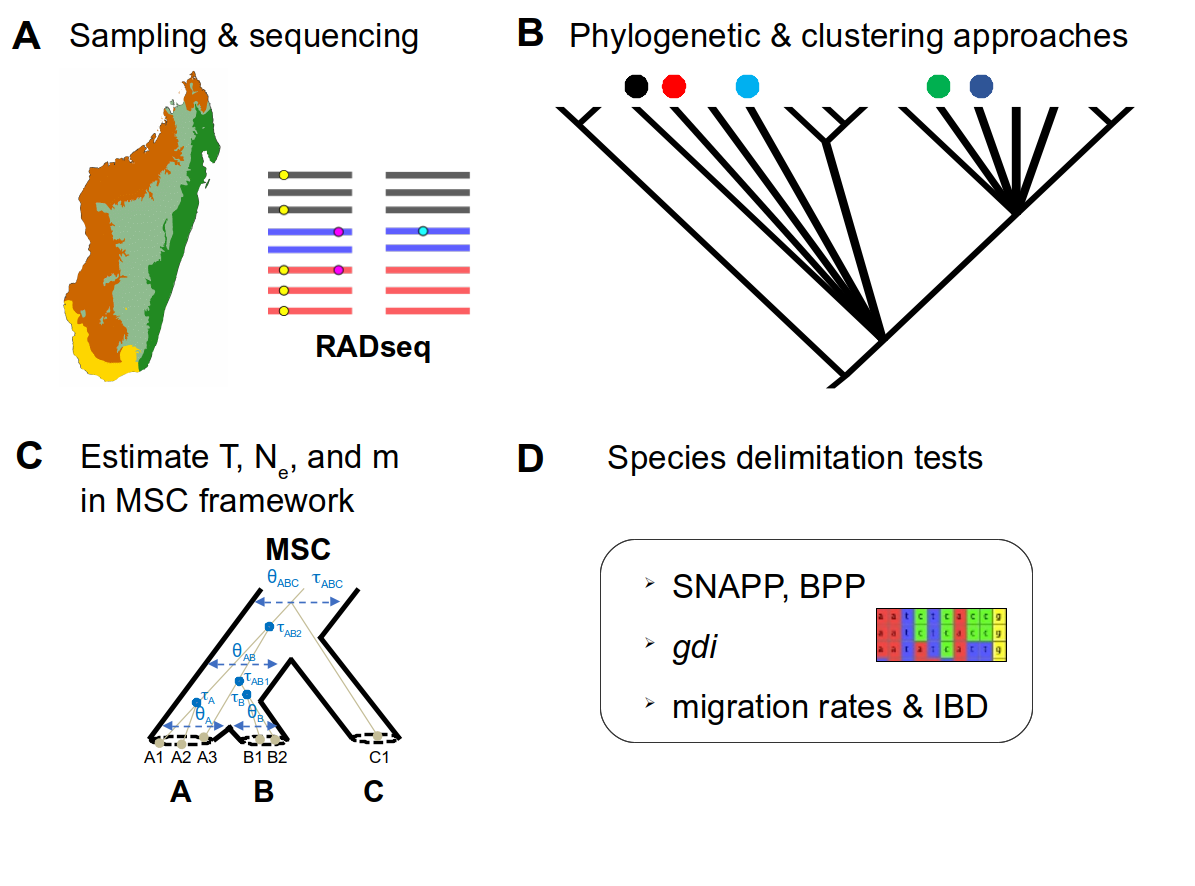

**Figure S1: Framework for delimiting species diversity in a cryptic radiation**
We use a structured framework for assessing species diversity within mouse lemurs in northeastern Madagascar. **a)** To prevent spurious genetic clustering due to isolation-by-distance, geographic sampling included intervening populations, with genomic characterization for all individuals. **b)** Using mtDNA barcoding, populations were placed in phylogenetic context with existing mtDNA sequences, thus allowing assignment to named species. Previous lack of assignment for a single M. sp. #3 individual provided the first indication that it might represent an undescribed species. **c)** Novel genomic data using RADseq were analyzed within the MSC framework (redrawn from Fujita et al. 2012) to determine divergence times (τ/T), effective population sizes (θ/Ne), and gene flow (m) for all populations. **d)** Genomic data were examined with multiple species delimitation tests to determine the degree of evolutionary distinctiveness of populations and phylogenetic lineages.

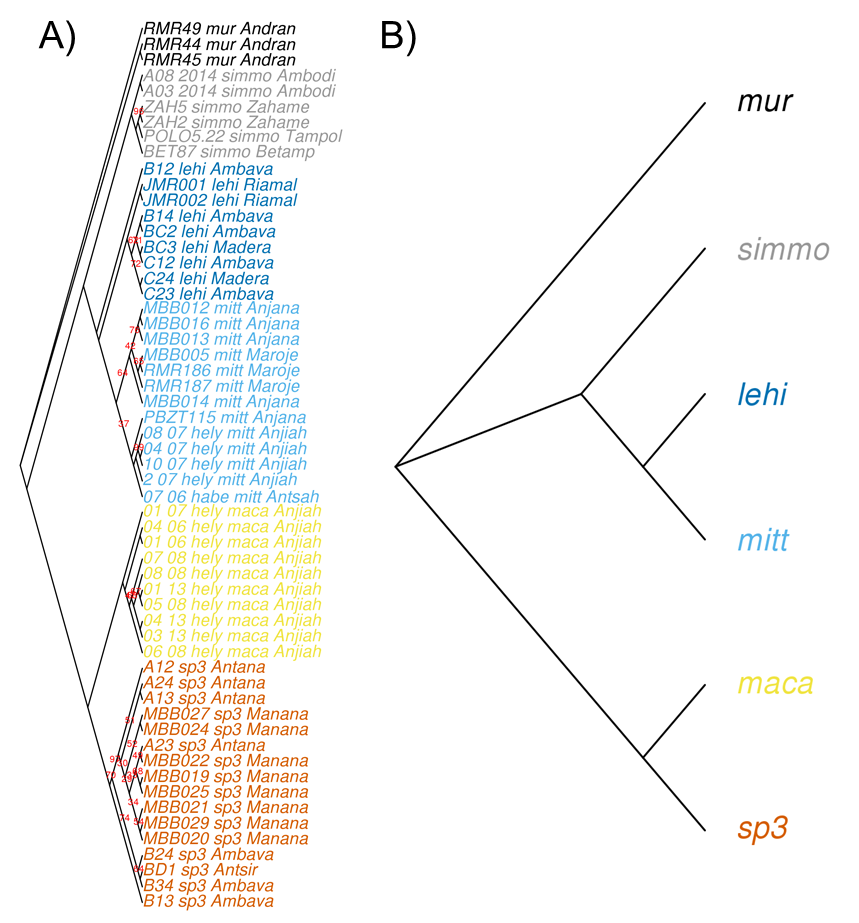

**Figure S2: Phylogenies inferred using SVDquartets**.
**(A)** SVDquartets individual-level topology inferred using RADseq nuclear data (nDNA) of 57 samples for which SNP calling was possible (Table S1).
**(B)** SVDquartets species based topology inferred using the same dataset with individuals attributed to groups based on the NgsAdmix results. SVDquartets topology node bootstrap values of a 100% are not indicated. Abbreviations: sp3: *M.* sp. #3; maca: *M. macarthurii*, mitt: *M. mittermeieri*; lehi: *M. lehilahytsara*; simmo: *M. simmonsi*; mur: *M. murinus*. For all trees, colors correspond to the NgsAdmix results at K = 6 and *M. murinus* is used as outgroup (Fig. S3).

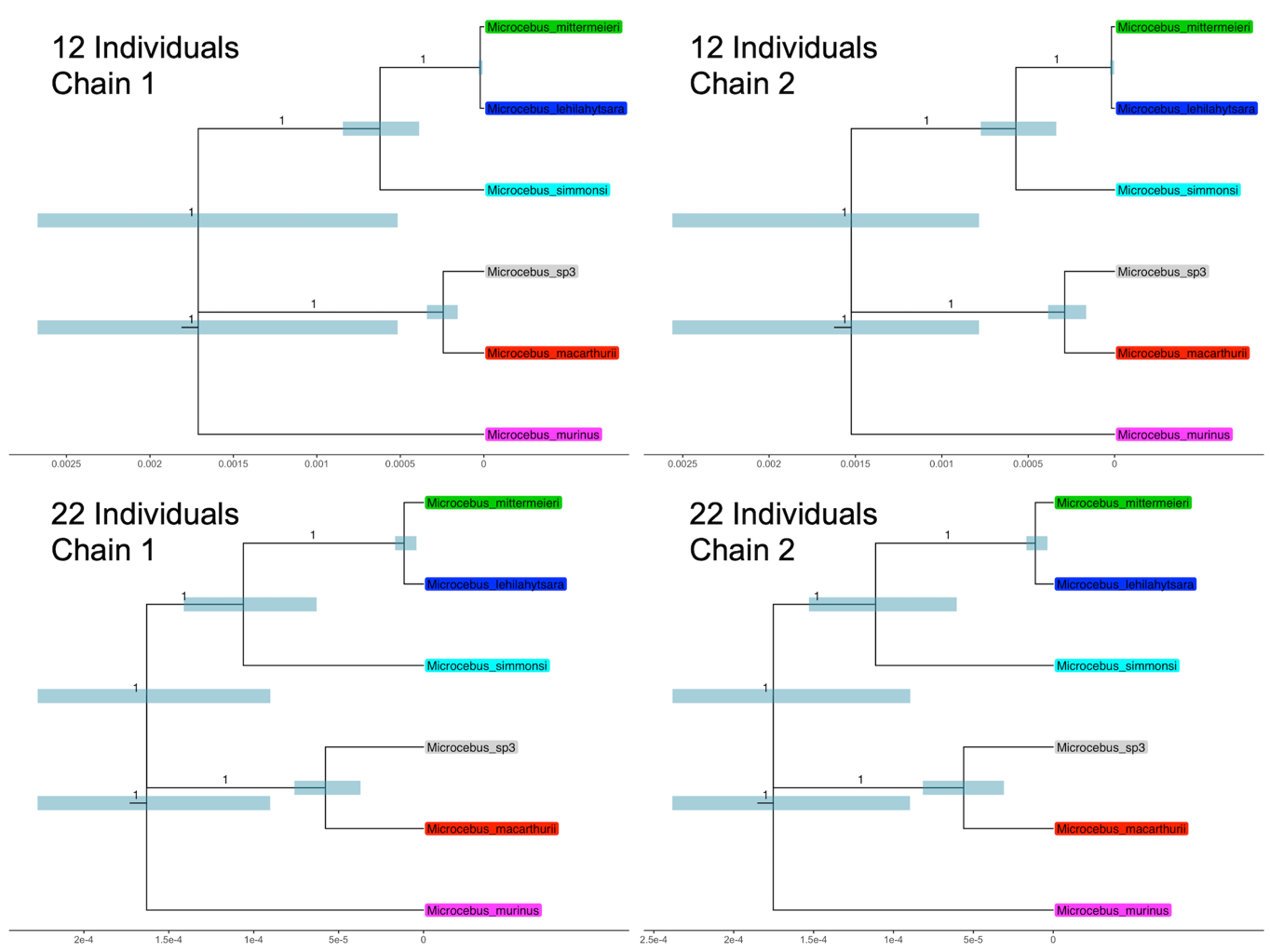
**Figure S3: MCCTs from SNAPP Analyses.**
Branch lengths are given in coalescent units. Error bars are the 95% highest posterior density interval around the median node height. All posterior probabilities are 1.0 and displayed for each node on subtending branches. node heights for 22-individual analyses are approximately ten times lower than twelve-individual analyses.

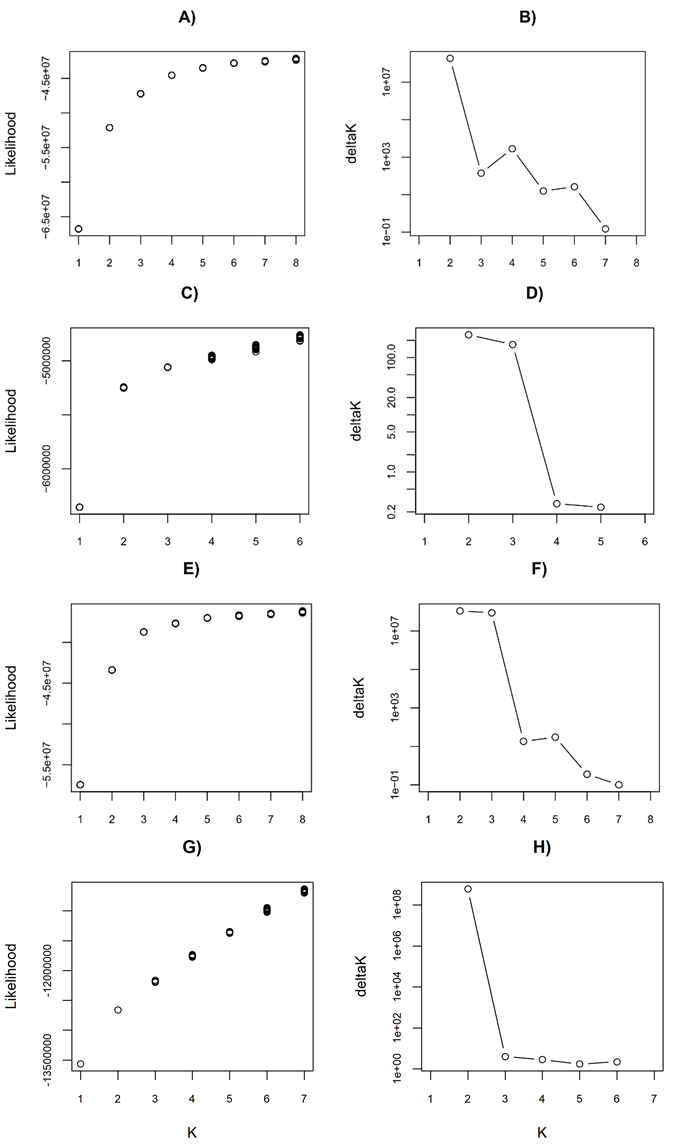

**Figure S4: Number of clusters inferred by NgsAdmix for different subsets.**
Likelihood **(A)** and deltaK **(B)** results for the complete dataset, including the outgroup *M. murinus*. Although the likelihood continues to increase slightly for values of K > 6, the results for K = 2, 4 and 6 likely best explain the data.

Likelihood (**C**) and deltaK (**D**) results for *M. macarthurii* and *M*. sp. #3 only. Although the likelihood continues to increase slightly for values of K>2, K=2 and 3 best explain the data following the deltaK method.

Likelihood (**E**) and deltaK (**F**) results for all species except outgroup (*M. murinus*). Although the likelihood continues to increase slightly for values of K > 5, the results for K = 2, 3 and 5 likely best explain the data.

Likelihood **(G)** and deltaK **(H)** results for *M. lehilahytsara* and *M. mittermeieri* only. The likelihood values continue to increase for values of K>2, K=2 and 3 best explain the data following the deltaK method.

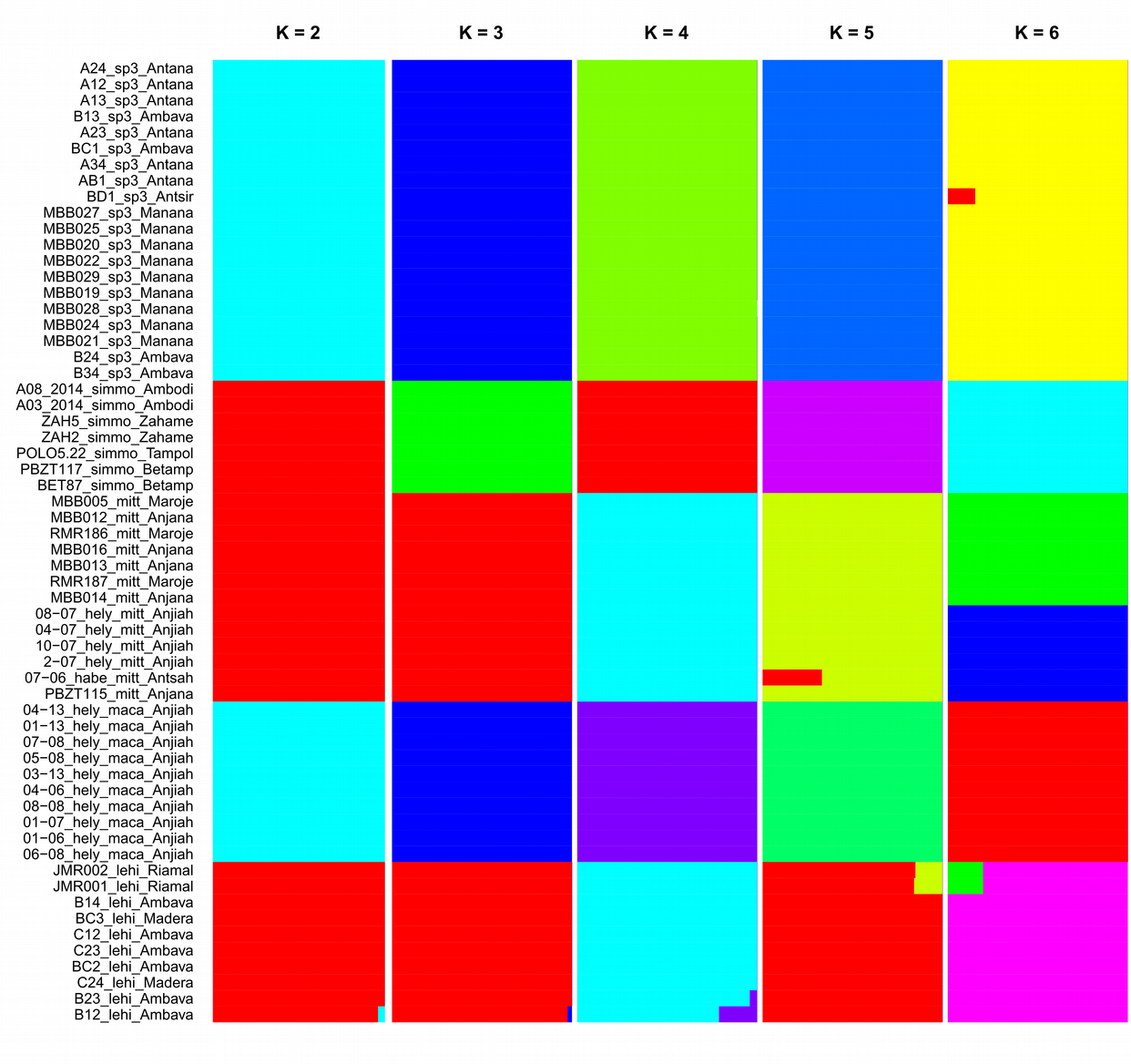

**Figure S5: NgsAdmix ancestry proportions for all individuals except the outgroup.**
Likelihood and deltaK analyses (Fig. S4E,F) suggest that the results for K=2, 3 and 5 likely best explain the data. Results for K = 7-8 are not shown.

Abbreviations: sp3: *M.* sp. #3; maca: *M. macarthurii*, mitt: *M. mittermeieri*; lehi: *M. lehilahytsara*; simmo: *M. simmonsi*.

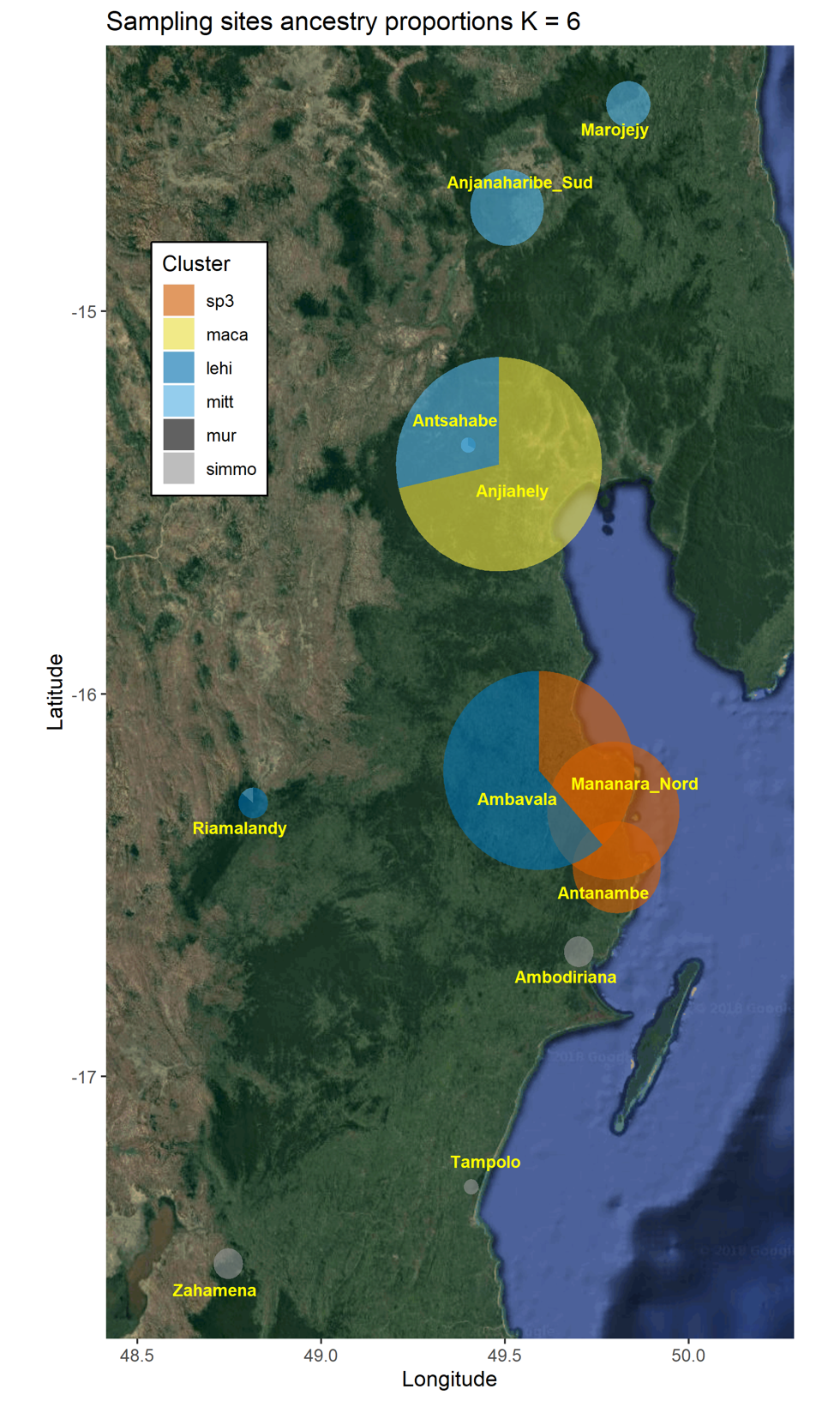

**Figure S6:** **NgsAdmix ancestry proportions for all individuals, plotted geographically.**
Pie chart sizes are proportional to the number of samples represented. Pie shares are proportional to the sums of individual ancestry proportions represented in Figure 4B. Results are represented for K = 6 according to likelihood and deltaK results in Figure S5.

Abbreviations: sp3: *M.* sp. #3; maca: *M. macarthurii*; mitt: *M. mittermeieri*; lehi: *M. lehilahytsara*; simmo: *M. simmonsi*.

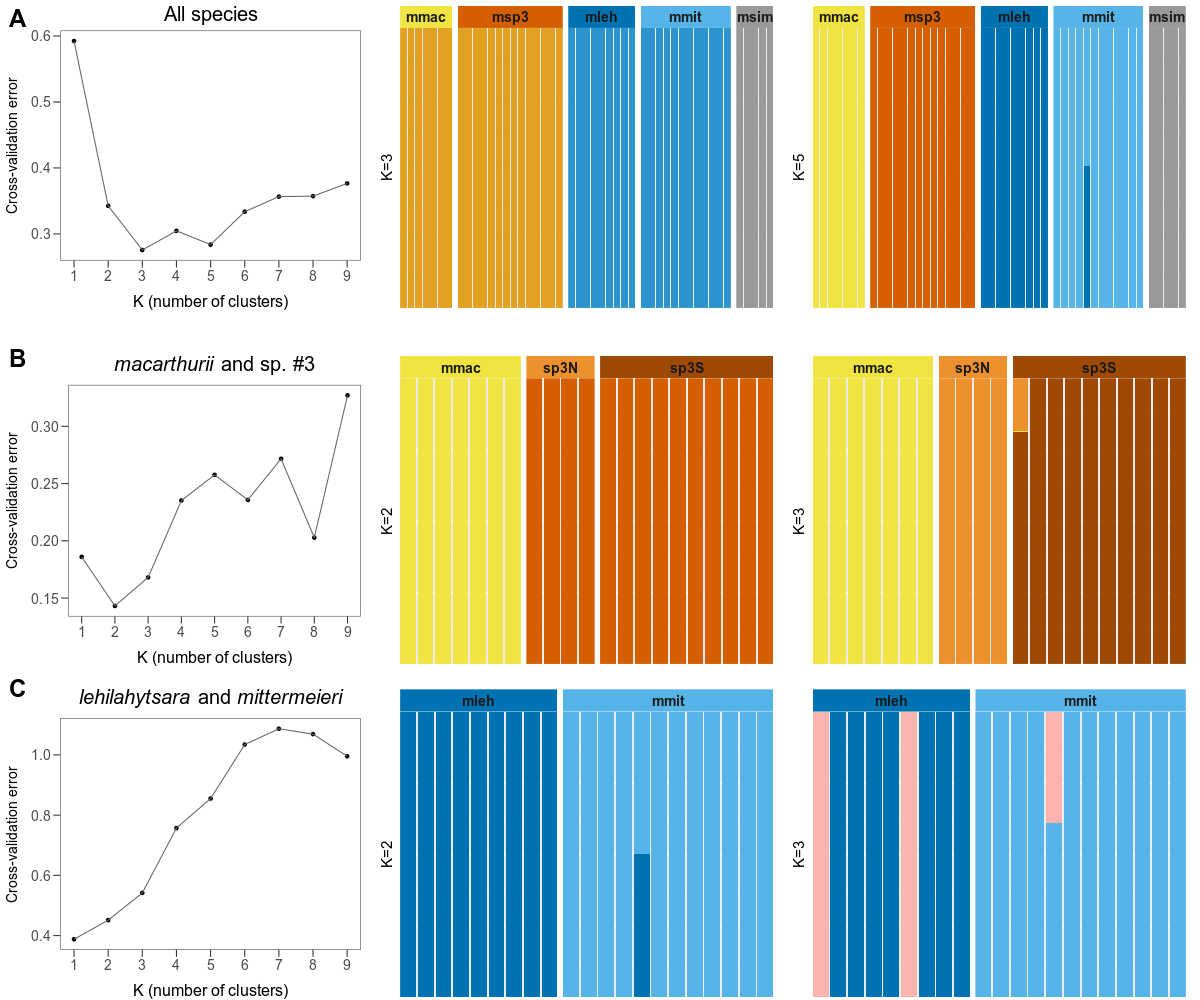
**Figure S7: ADMIXTURE results.**

Figures on the left show the cross-validation (CV) error for K from 1 to 9, with a lower error providing a better fit. On the right, cluster assignment results for the two best-supported values of K are shown.

**A)** Using individuals from all five focal lineages. K=3 and K=5 have the lowest CV error, and barplots for those two values of K are shown.

**B)** Using only individuals from *M. macarthurii* and *M.* sp. #3*.* K=2 has the lowest CV error. At K=3, *M*. sp. #3 individuals are separated into northern (“sp3N”) and southern (“sp3S”) population groups.

**C)** Using only individuals from *M. lehilahytsara* and *M. mittermeieri*. K=1 has the lowest CV error. Nevertheless, at K=2, ancestry is almost fully separated by nominal species. Results for K=3 are also plotted.

Abbreviations: mleh: *M. lehilahytsara*, mmac: *M. macarthurii*, mmit: *M. mittermeieri*, msim: *M. simmonsi*, msp3: *M*. sp. #3.

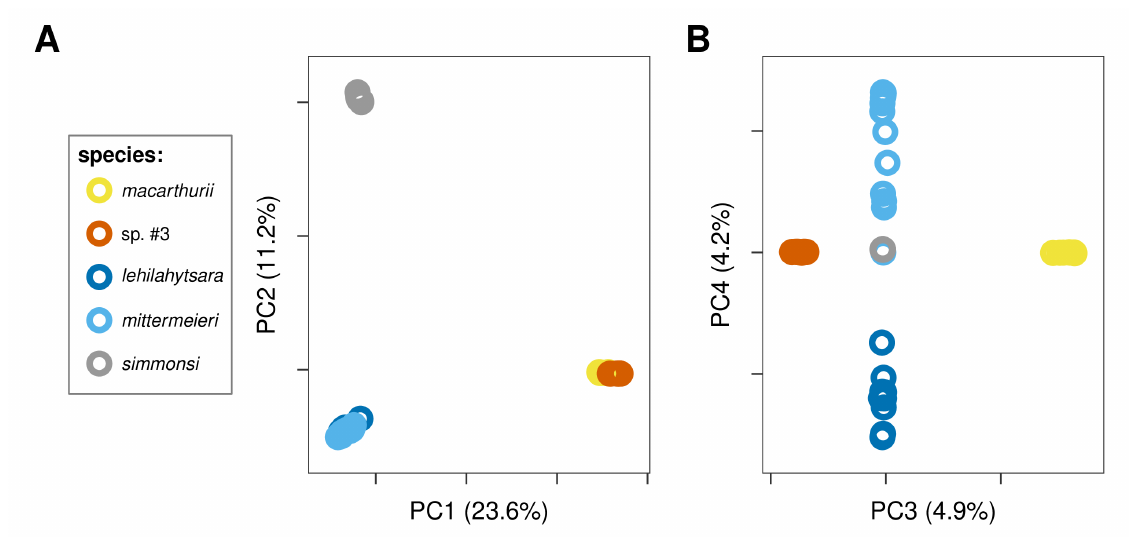

**Figure S8: PCA results (using GATK genotypes) for all 5 ingroup species.**

**A)** PC1 and PC2

**B)** PC3 and PC4

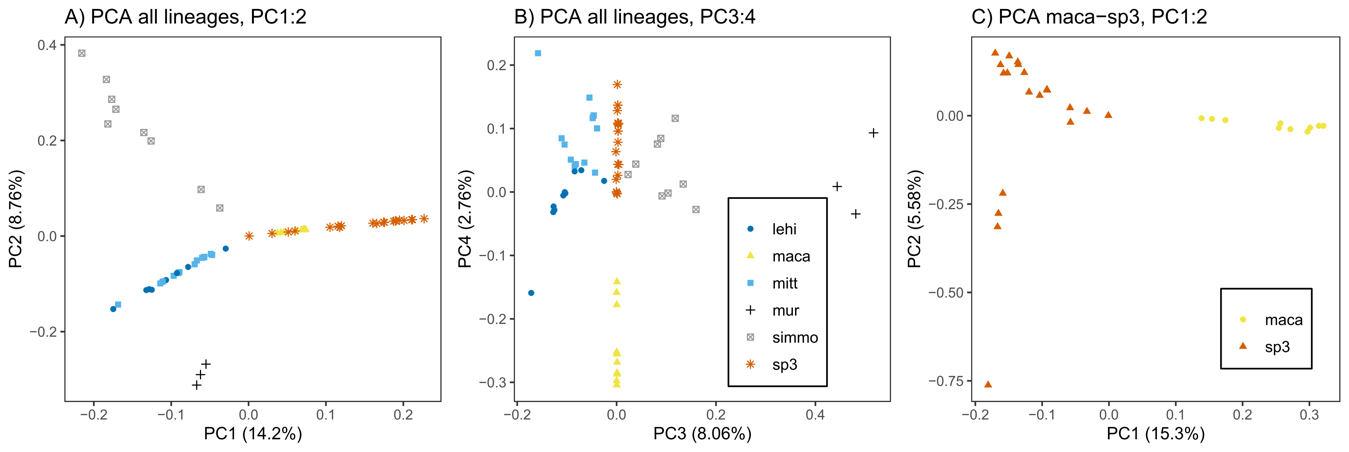

**Figure S9: ANGSD-PCA results for all 5 species.**

Results are shown for all species (**A & B**) and among *M. macarthurii* and *M.* sp. *#3* only (**C**). Abbreviations: lehi = *M. lehilahytsara*, maca = *M. macarthurii*, mitt = *M. mittermeieri*, mur = *M. murinus*, simmo = *M. simmonsi*, sp3 = *M*. sp. #3.

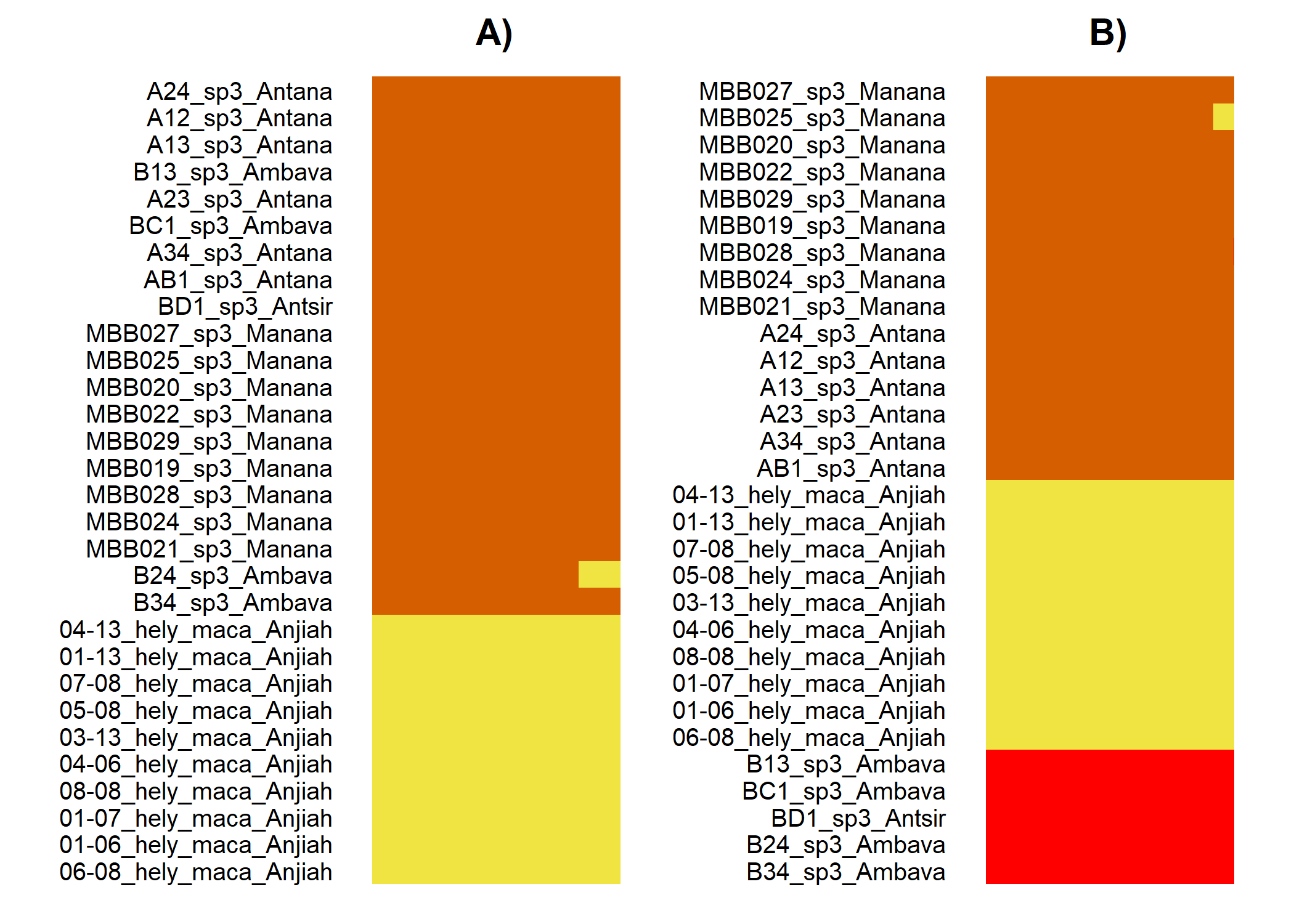

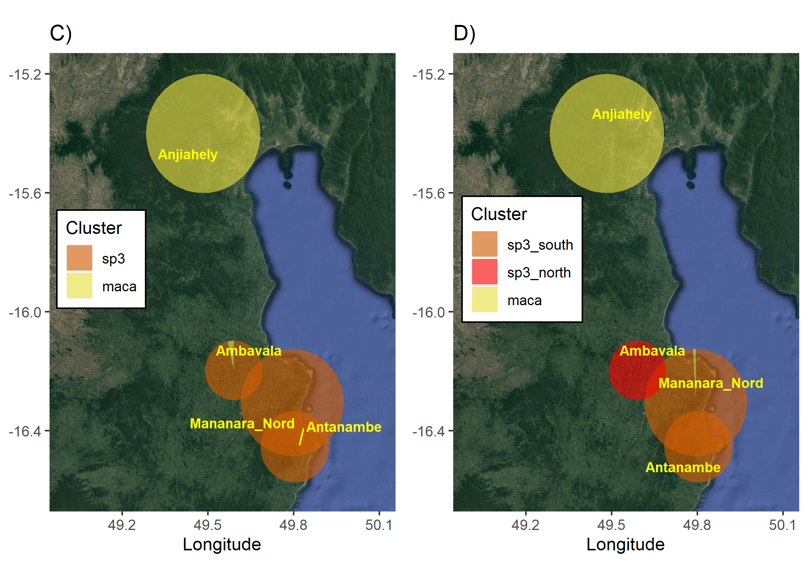

**Figure S10: ngsAdmix ancestry proportions for *M. macarthurii* and *M.* sp. *#3.***

**(A,B)** Individual ancestry proportions inferred in NgsAdmix. **(C,D)** Geographic representation of ancestry proportions per site. Results are represented for K = 2 **(A,C)** and K = 3 **(B,D)** according to likelihood and deltaK results in Figure S5. Pie chart size is proportional to the number of samples represented. Pie shares **(C,D)** are proportional to the sums of individual ancestry proportions in **A** and **B**. Abbreviations: sp3: *M.* sp. #3; maca: *M. macarthurii.*

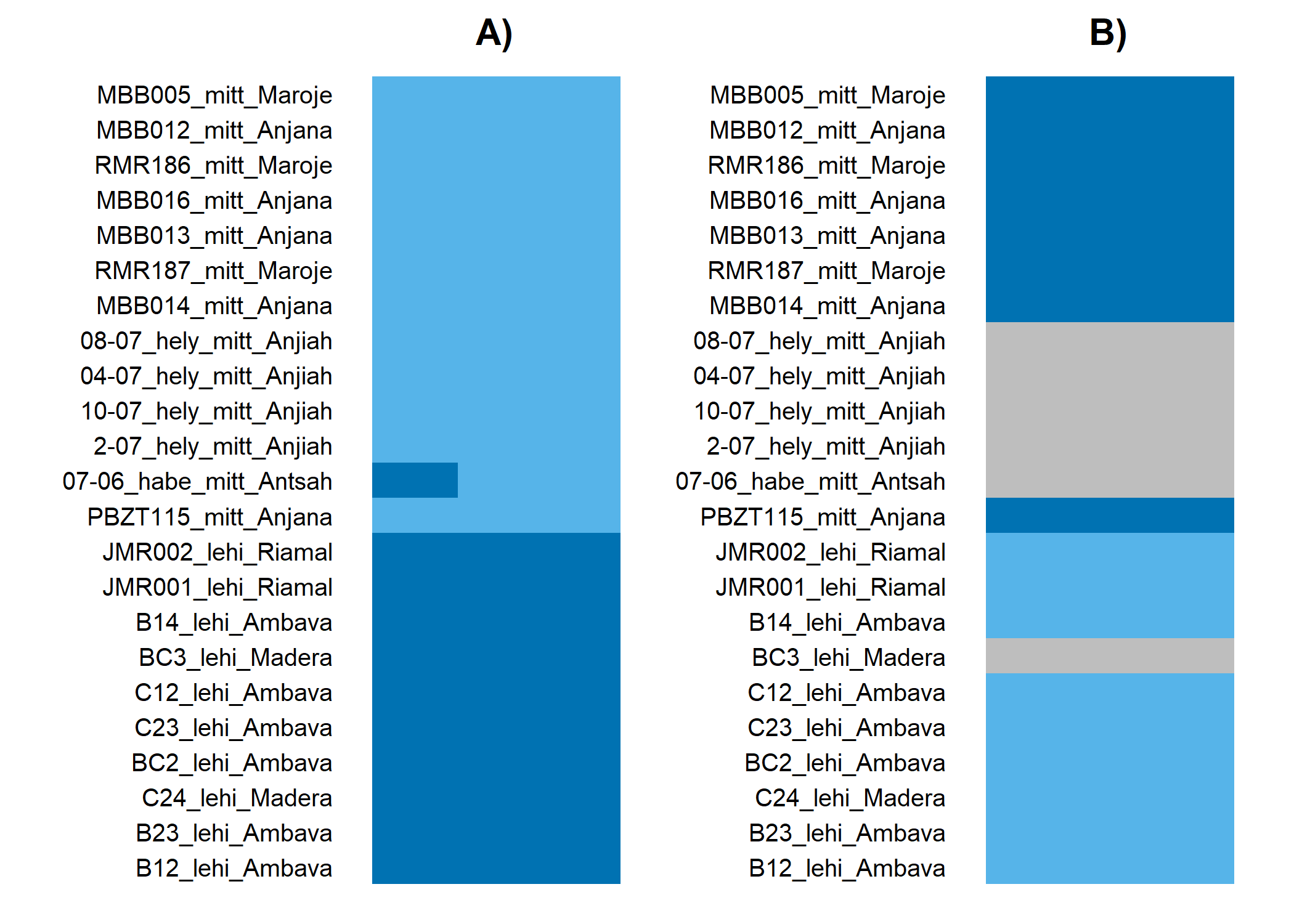

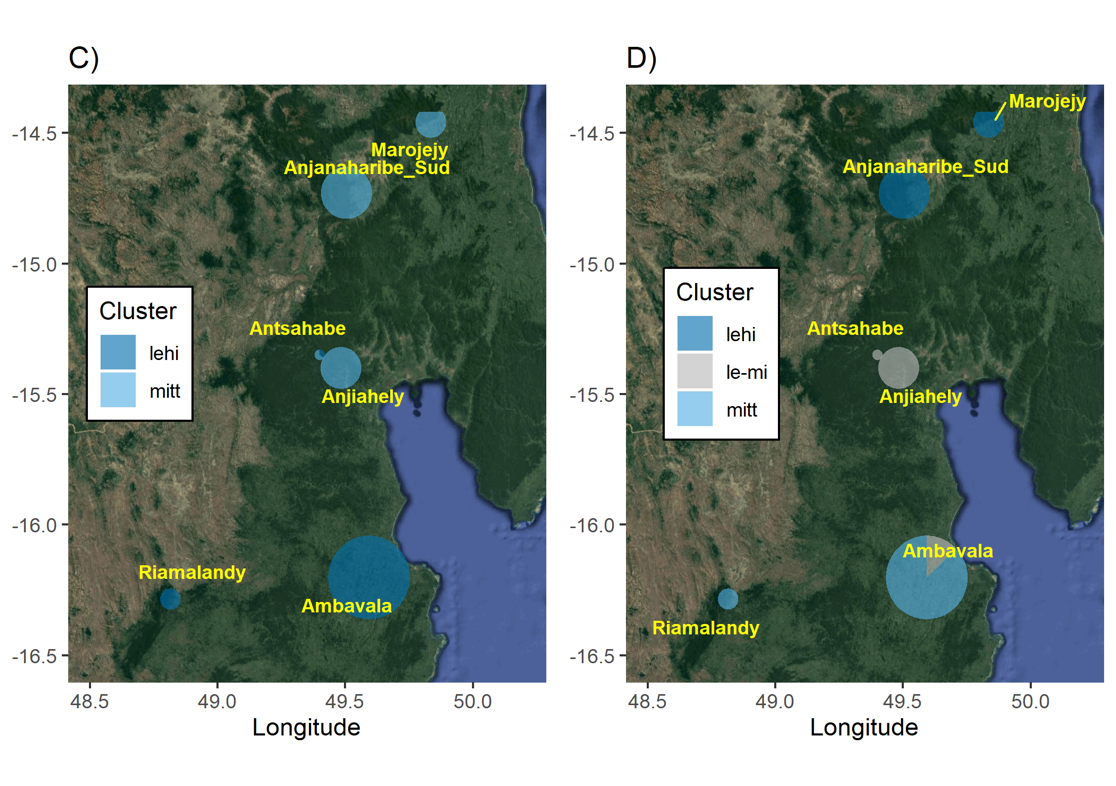

**Figure S11: ngsAdmix ancestry proportions for *M. lehilahytsara* and *M.* mittermeieri**

**(A,B)** Individual ancestry proportions inferred in NgsAdmix. **(C,D)** Geographic representation of ancestry proportions per site. Results are represented for K = 2 **(A,C)** and K = 3 **(B,D)** according to likelihood and deltaK results in Figure S5. Pie chart size is proportional to the number of samples represented. Pie shares **(C,D)** are proportional to the sums of individual ancestry proportions in **A** and **B**.

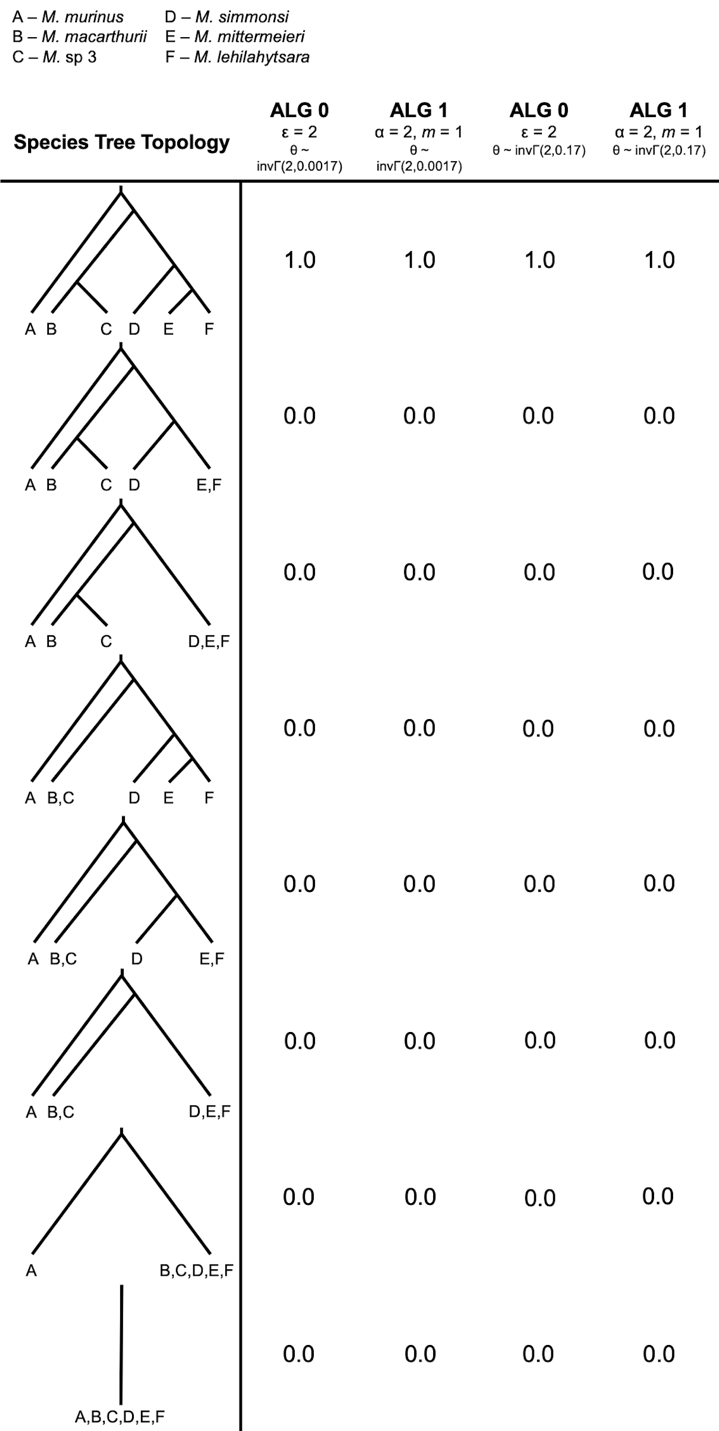
 **Figure S12: Guided species delimitation with BPP.**
Two different algorithms were used with two different priors for θ. θ ~ invΓ(2,0.0017) is our informed prior based on previous nucleotide diversity estimates while θ ~ invΓ(2,0.17) is assumed to be mis-specified and inappropriately high. Still, the species delimitation model that did not lump any named lineages was recovered in all posteriors across all four algorithm and prior combinations.

**Figure S13: Effective population sizes and migration rates inferred by G-PhoCS for a 3-species model.**

*M. macarthurii, M. sp. #3* (divided in northern and southern population groups) and *M. lehilahytsara* are included.

**(A)** Effective population sizes inferred under an isolation model (green) and a model with gene flow (orange).

**(B)** Migration rates between *M. lehilahytsara* and lineages in the *M.* sp. #3 – *M. macarthurii* clade. For models using single migration bands (orange), low but significant gene flow is inferred for each possible comparison, with highest levels of gene flow inferred from northern and southern *M.* sp. #3 to *M. lehilahytsara.* However, a model with multiple migration bands (green) instead infers migration from the lineage ancestral to *M.* sp. #3 and *M. macarthurii* to *M. lehilahytsara* to be significant (i.e., 95% HPD intervals exclude 0), while migration from northern *M.* sp. #3 to *M. lehilahytsara* is n longer significant. Therefore, we cannot conclusively determine the relative timing and direction of migration between *M. lehilahytsara* and lineages in the *M.* sp. #3 – *M. macarthurii* clade.

*Abbreviations: mac =* M. macarthurii*, sp3N = northern* M. *sp. #3, sp3S = southern* M. *sp. #3, sp3A =* M. *sp. #3 ancestral lineage (to northern and southern populations); anc_ms = lineage ancestral to* M. macarthurii *and* M. sp. #3; leh = M. lehilahytsara; root = lineage ancestral to all included extant lineages.

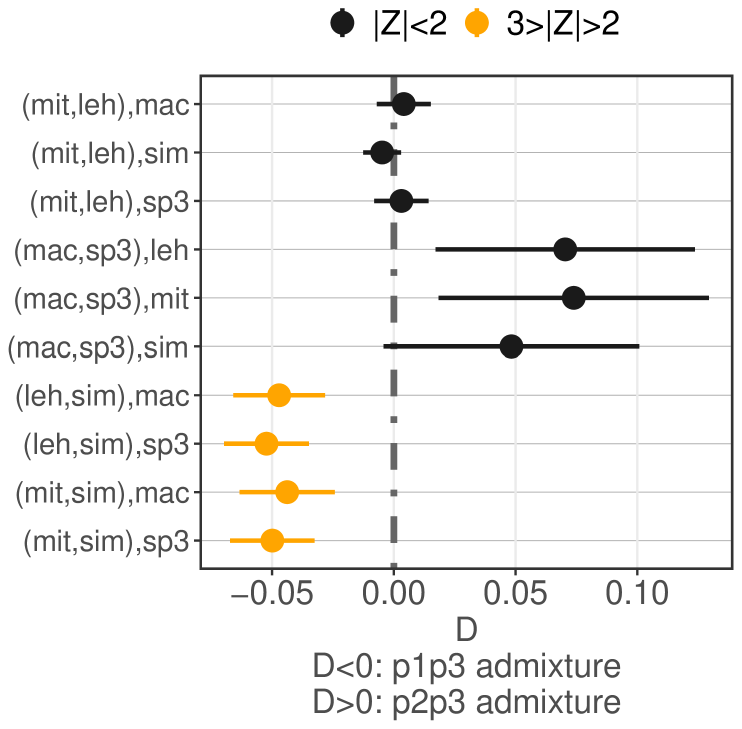

**Figure S14: D-statistics testing for introgression among non-sister species.**

The lower four comparisons (in orange) suggest an excess of allele sharing between *lehilahytsara*/*mittermeieri* on the one hand, and *macarthurii*/sp. #3 on the other hand, although the effect is not significant (Z-score below 3). Axis annotations denote “(P1, P2),P3”, wherein gene flow is tested between P3 and either P1 (negative D) or P2 (positive D). In all cases, P4 is the outgroup *M. murinus*.

**Figure S15: Gene flow between *M*. sp. #3 and *M. macarthurii*.**
Gene flow inferred by G-PhoCS under a model with three species, in units of **(A)** the population migration rate (2Nm) and **(B)** the percentage of migrants in each generation.
**(C)** Four-taxon D-statistics suggest gene flow between *M. macarthurii* with mitonuclear discordance (“mac*”) and northern *M*. sp. #3 (“sp3N”). *M. macarthurii* with and without mitonuclear discordance (“mac”) were tested separately. Axis annotations denote “(P1, P2),P3”, wherein gene flow is tested between P3 and either P1 (negative D) or P2 (positive D). Significant comparisons (|Z| > 3) are in red.

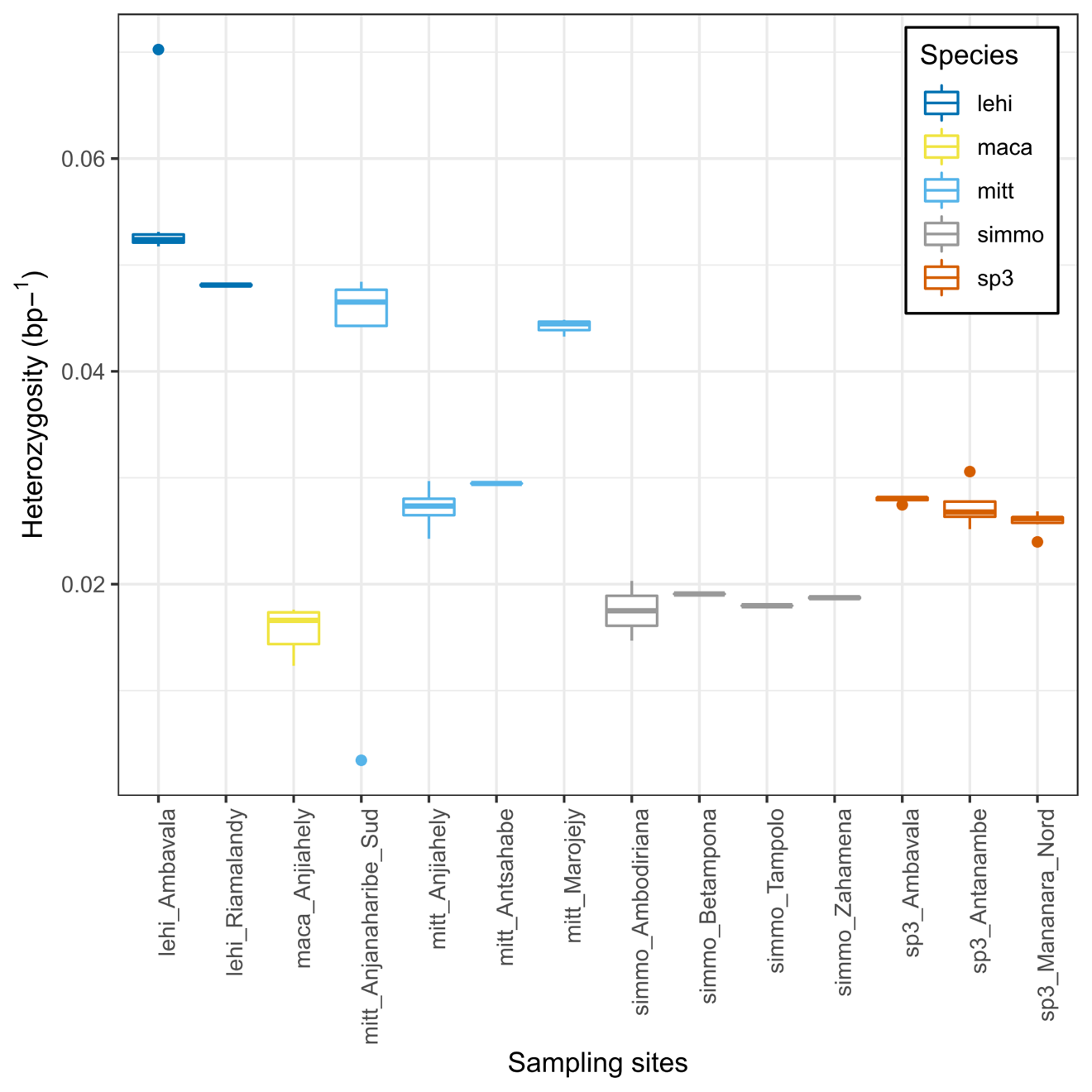

**Figure S16: Genetic diversity.**

Heterozygosity per species and sampling site given as boxplots of heterozygosity per base pair.
Abbreviations: lehi = *M. lehilahytsara*, maca = *M. macarthurii*, mitt = *M. mittermeieri*, simmo = *M. simmonsi*, sp3 = *M*. sp. #3.

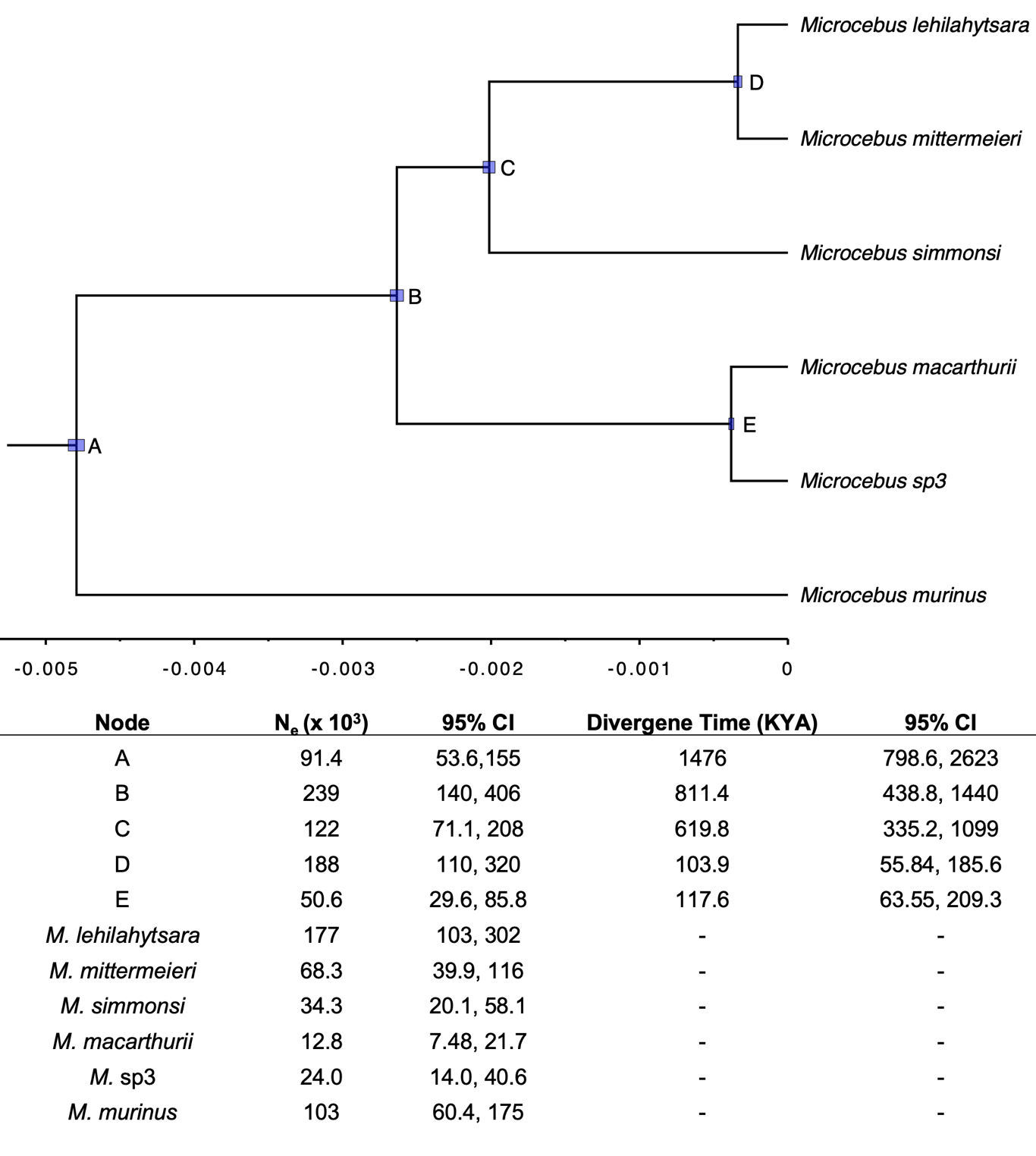

**Figure S17: Divergence times and *N_e_* estimates from BPP for the 12-individual dataset.**

Branch lengths are in coalescent units and error bars represent 95% highest posterior densities. Median parameter estimates and 95% HPDs are given in the table below.

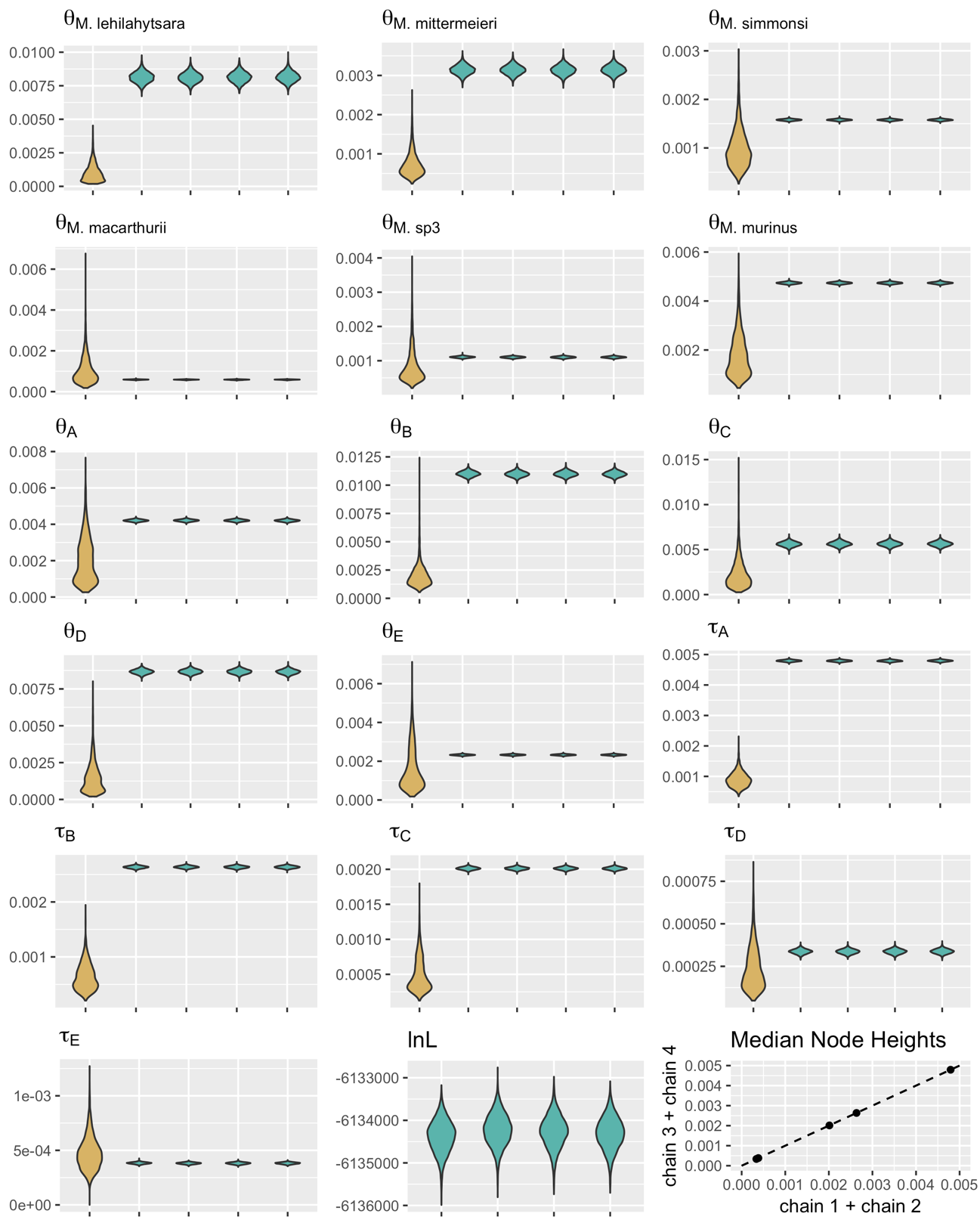

**Figure S18: Posterior distributions of BPP chains and marginal priors.**
7000 post-burn-in samples were collected for each of four independent chains. The marginal prior distributions are shown in yellow while posteriors for each model parameter are shown in green. The log-likelihoods of each chain were similar after collecting 7000 samples. Chains 1 and 2 were combined and compared to node heights from chains 3 and 4 to check convergence. node abbreviations are given on Figure S11.

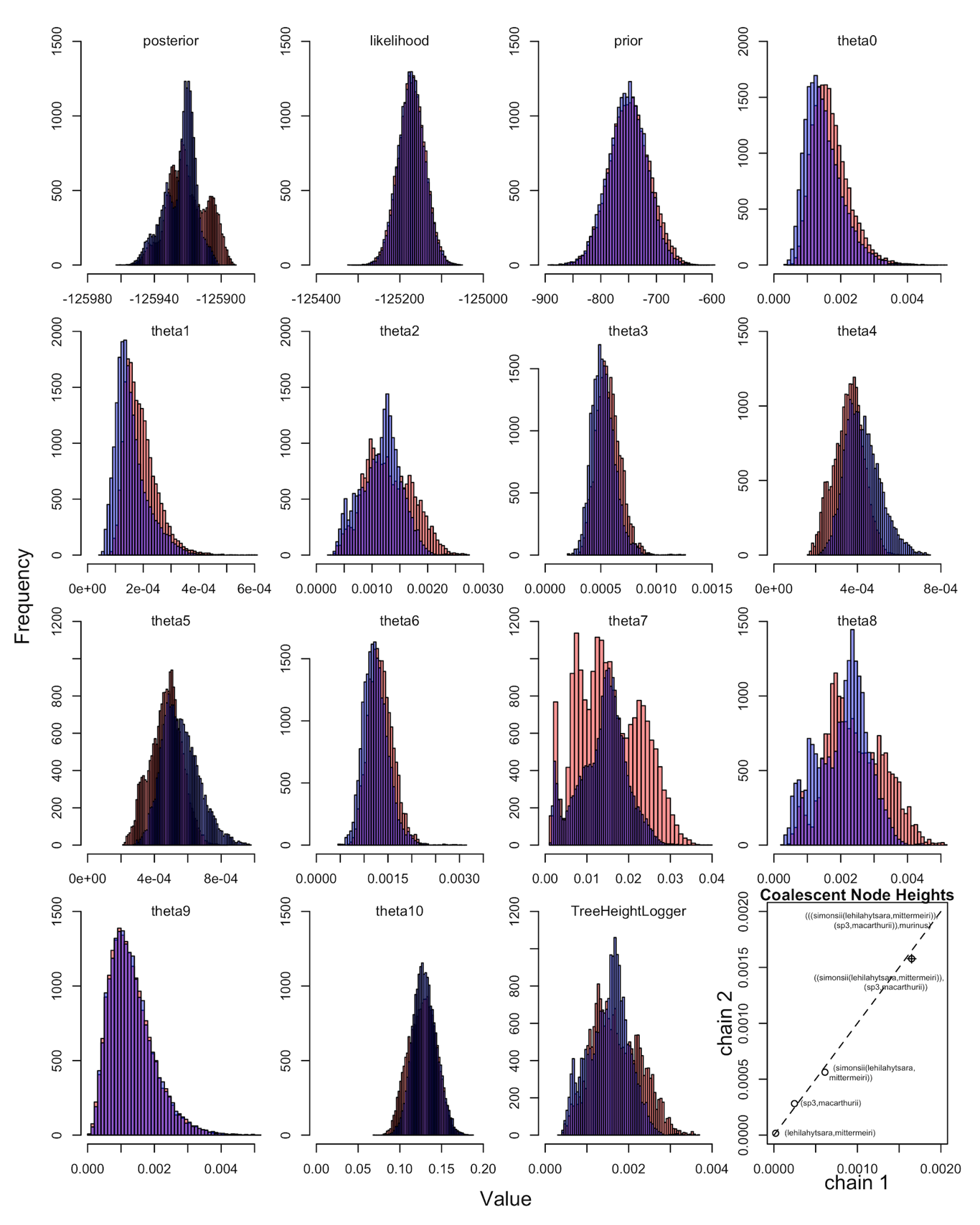

**Figure S19: Posterior Distributions of 12-individual SNAPP chains.**
Analyses were performed with the 12-individual dataset, collecting 20,000 post-burn-in samples for each posterior. Chain 1 is shown in red and chain 2 is shown in blue. Median tree heights measured in coalescent units are also given for the two species tree estimates.

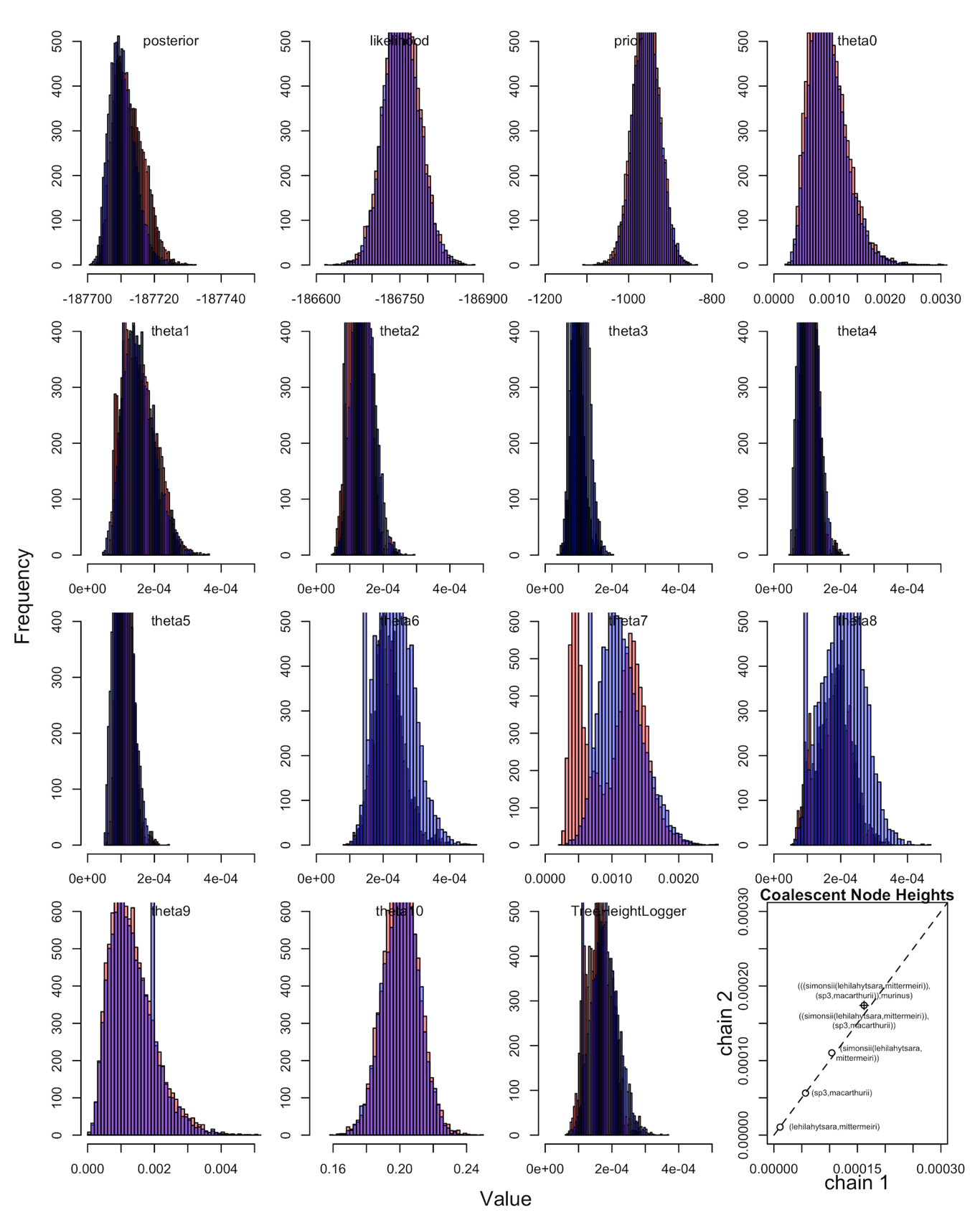

**Figure S20: Posterior Distributions of 22-individual SNAPP Chains.**
Both posteriors collected 10000 post-burn-in samples. Chain 1 is shown in red and chain 2 is shown in blue. Median tree heights measured in coalescent units are also given for the two species tree estimates.

### 4 Supplementary Tables

**Table S1: Samples and genetic data used in this study.**
“Read type” column: pe: paired-end reads; se: single-end reads.
“Library prep” column: IDH: Idaho; ORG: Oregon; TLS: Toulouse.
“Library protocol” column: 1-3 correspond to protocol numbers above described in supplementary methods.
“SNAPP-22” column: 22-individual SNAPP dataset.
“SNAPP-12, BPP, G-PhoCS-all” column: 12-individual SNAPP dataset which was also used for BPP and G-PhoCS (with all species) analyses.
“G-PhoCS-cladeI” column: G-PhoCS dataset with only *macarthurii*, *M.* sp. #3, and *lehilahytsara*.
Other abbreviations: QC: quality control and filtering; F&R: Forward and reverse reads. F1: forward first in pair read.

| Species | Individual ID | Locality | Sex | Latitude | Longitude | Read type | Library prep | Library protocol | Illumina sequencer | # of raw reads | # of reads after QC | # of aligned reads F&R | # of loci >=3X | Mean F1 coverage | SNP calling | SNAPP-22 | SNAPP-12, BPP, G-PhoCS-all | G-PhoCS-CladeI | dloop | coii | cytb | Collection institution |
| --- | --- | --- | --- | --- | --- | --- | --- | --- | --- | --- | --- | --- | --- | --- | --- | --- | --- | --- | --- | --- | --- | --- |
| *jonahi* | B34 | Ambavala | M | -16.20511 | 49.58952 | pe | TLS | 2 | HS3000 | 6 331 612 | 4 802 052 | 3 977 364 | 140 105 | 10,52 | y | **y** | **y** | **y** | n | y | n | Tiho |
| *jonahi* | B24 | Ambavala | M | -16.19493 | 49.59785 | pe | TLS | 2 | HS3000 | 19 347 868 | 15 049 782 | 10 892 366 | 166 715 | 21,76 | y | n | n | n | n | y | n | Tiho |
| *jonahi* | BC1 | Ambavala | F | -16.19976 | 49.59566 | pe | TLS | 3 | HS4000 | 810 532 | 643 440 | 527 546 | 39 846 | 2,23 | **n** | n | n | n | n | y | y | Tiho |
| *jonahi* | B13 | Ambavala | M | -16.20228 | 49.58858 | pe | TLS | 3 | HS4000 | 6 808 818 | 5 467 724 | 4 453 669 | 143 784 | 11,19 | y | n | n | **y** | n | y | y | Tiho |
| *jonahi* | AB1 | Antanambe | F | -16.45540 | 49.80431 | pe | TLS | 3 | HS4000 | 457 202 | 362 414 | 297 959 | 14 288 | 1,65 | **n** | n | n | n | n | y | y | Tiho |
| *jonahi* | A34 | Antanambe | M | -16.43802 | 49.81510 | pe | TLS | 3 | HS4000 | 1 016 652 | 630 092 | 460 018 | 30 357 | 1,99 | **n** | n | n | n | n | y | y | Tiho |
| *jonahi* | A23 | Antanambe | M | -16.46017 | 49.80275 | pe | TLS | 3 | HS4000 | 4 909 794 | 3 030 744 | 2 538 688 | 128 556 | 7,12 | y | n | n | n | n | y | y | Tiho |
| *jonahi* | A13 | Antanambe | M | -16.45520 | 49.80404 | pe | TLS | 3 | HS4000 | 1 906 404 | 1 490 586 | 1 130 519 | 95 455 | 3,74 | y | n | n | n | n | y | y | Tiho |
| *jonahi* | A12 | Antanambe | M | -16.45230 | 49.79654 | pe | TLS | 3 | HS4000 | 5 731 186 | 4 563 310 | 3 668 882 | 140 965 | 9,40 | y | **y** | **y** | **y** | n | y | y | Tiho |
| *jonahi* | A24 | Antanambe | M | -16.45879 | 49.80289 | pe | TLS | 3 | HS4000 | 4 440 680 | 3 539 618 | 2 727 330 | 133 817 | 7,49 | y | n | n | n | n | y | y | Tiho |
| *jonahi* | BD1 | Antsiradran | F | -16.18398 | 49.57840 | pe | TLS | 3 | HS4000 | 7 127 576 | 5 683 252 | 4 498 153 | 145 431 | 11,40 | y | n | n | **y** | n | y | y | Tiho |
| *jonahi* | MBB021 | Mananara_nrd | un | -16.30482 | 49.79468 | pe | IDH | 3 | HS4000 | 3 427 586 | 2 999 486 | 1 288 988 | 77 080 | 5,11 | y | n | n | n | n | y | y | Duke Univ |
| *jonahi* | MBB024 | Mananara_nrd | un | -16.30482 | 49.79468 | pe | IDH | 3 | HS4000 | 4 063 008 | 3 707 652 | 999 804 | 77 333 | 3,65 | y | n | n | n | n | y | y | Duke Univ |
| *jonahi* | MBB028 | Mananara_nrd | un | -16.30482 | 49.79468 | pe | IDH | 3 | HS4000 | 33 038 | 30 382 | 13 349 | 20 | 1,05 | **n** | n | n | n | n | y | y | Duke Univ |
| *jonahi* | MBB019 | Mananara_nrd | un | -16.30482 | 49.79468 | pe | IDH | 3 | HS4000 | 11 918 486 | 10 802 820 | 3 322 977 | 111 564 | 10,79 | y | **y** | n | **y** | n | y | y | Duke Univ |
| *jonahi* | MBB029 | Mananara_nrd | un | -16.30482 | 49.79468 | pe | IDH | 3 | HS4000 | 6 533 662 | 5 923 460 | 1 827 833 | 118 802 | 5,95 | y | n | n | n | n | y | y | Duke Univ |
| *jonahi* | MBB022 | Mananara_nrd | un | -16.30482 | 49.79468 | pe | IDH | 3 | HS4000 | 5 730 988 | 5 218 624 | 1 582 689 | 74 793 | 6,74 | y | n | n | n | n | y | y | Duke Univ |
| *jonahi* | MBB020 | Mananara_nrd | un | -16.30482 | 49.79468 | pe | IDH | 3 | HS4000 | 7 480 922 | 6 890 592 | 2 321 942 | 102 460 | 8,06 | y | n | n | **y** | n | y | y | Duke Univ |
| *jonahi* | MBB025 | Mananara_nrd | un | -16.30482 | 49.79468 | pe | IDH | 3 | HS4000 | 12 525 250 | 11 358 552 | 3 534 544 | 121 620 | 10,99 | y | n | n | n | n | y | y | Duke Univ |
| *jonahi* | MBB027 | Mananara_nrd | un | -16.30482 | 49.79468 | pe | IDH | 3 | HS4000 | 12 815 198 | 11 666 916 | 3 529 347 | 111 384 | 11,49 | y | **y** | n | **y** | n | y | y | Duke Univ |
| *lehilahytsara* | B12 | Ambavala | F | -16.20431 | 49.59652 | pe | TLS | 2 | HS3000 | 7 406 590 | 5 907 070 | 4 594 466 | 155 911 | 10,29 | y | **y** | **y** | n | n | y | y | Tiho |
| *lehilahytsara* | B23 | Ambavala | F | -16.20427 | 49.59650 | pe | TLS | 2 | HS3000 | 831 614 | 653 672 | 522 259 | 38 398 | 2,15 | **n** | n | n | n | n | y | y | Tiho |
| *lehilahytsara* | BC2 | Ambavala | F | -16.19803 | 49.59940 | pe | TLS | 2 | HS3000 | 4 542 814 | 3 461 628 | 2 775 412 | 137 731 | 7,39 | y | n | n | n | n | y | n | Tiho |
| *lehilahytsara* | C23 | Ambavala | M | -16.19823 | 49.59842 | pe | TLS | 2 | HS3000 | 4 857 192 | 3 698 298 | 2 957 616 | 140 887 | 7,60 | y | n | n | n | n | n | y | Tiho |
| *lehilahytsara* | C12 | Ambavala | F | -16.19782 | 49.59984 | pe | TLS | 2 | HS3000 | 5 886 578 | 4 663 676 | 3 660 306 | 149 276 | 9,11 | y | n | n | **y** | n | y | y | Tiho |
| *lehilahytsara* | B14 | Ambavala | M | -16.19767 | 49.59962 | pe | TLS | 2 | HS3000 | 4 731 492 | 3 623 670 | 2 947 864 | 141 129 | 7,69 | y | n | n | n | n | y | y | Tiho |
| *lehilahytsara* | C24 | Madera | M | -16.20589 | 49.57872 | pe | TLS | 2 | HS3000 | 7 581 894 | 5 779 868 | 4 613 584 | 155 293 | 11,01 | y | **y** | n | **y** | n | y | y | Tiho |
| *lehilahytsara* | BC3 | Madera | F | -16.21118 | 49.57797 | pe | TLS | 2 | HS3000 | 21 269 490 | 16 584 708 | 11 405 405 | 187 016 | 21,07 | y | n | n | n | n | y | y | Tiho |
| *lehilahytsara* | JMR001 | Riamalandy | un | -16.28500 | 48.81500 | pe | IDH | 3 | HS4000 | 5 143 110 | 4 657 274 | 1 775 476 | 109 391 | 5,01 | y | **y** | **y** | n | n | n | n | Duke Univ |
| *lehilahytsara* | JMR002 | Riamalandy | un | -16.28500 | 48.81500 | pe | IDH | 3 | HS4000 | 9 495 954 | 8 643 014 | 2 200 177 | 122 718 | 5,95 | y | **y** | n | n | n | n | n | Duke Univ |
| *macarthuri* | 06-08_hely | Anjiahely | M | -15.40000 | 49.48333 | se | ORG | 1 | HS2000 | 3 599 276 | 3 553 490 | 2 190 056 | 147 865 | 9,71 | y | **y** | n | **y** | y | y | y | Tiho |
| *macarthuri* | 01-06_hely | Anjiahely | M | -15.40000 | 49.48333 | se | ORG | 1 | HS2000 | 3 160 436 | 3 119 042 | 2 325 447 | 150 718 | 10,21 | y | n | n | n | y | y | y | Tiho |
| *macarthuri* | 01-07_hely | Anjiahely | F | -15.40000 | 49.48333 | se | ORG | 1 | HS2000 | 3 005 417 | 2 966 694 | 2 108 898 | 143 836 | 9,62 | y | n | n | n | y | y | n | Tiho |
| *macarthuri* | 08-08_hely | Anjiahely | M | -15.40000 | 49.48333 | se | ORG | 1 | HS2000 | 3 707 388 | 3 652 455 | 2 505 069 | 152 219 | 10,70 | y | **y** | **y** | **y** | y | y | n | Tiho |
| *macarthuri* | 04-06_hely | Anjiahely | F | -15.40000 | 49.48333 | se | ORG | 1 | HS2000 | 3 078 601 | 3 039 948 | 1 904 193 | 135 589 | 9,22 | y | n | n | n | y | y | y | Tiho |
| *macarthuri* | 03-13_hely | Anjiahely | F | -15.40000 | 49.48333 | se | ORG | 1 | HS2000 | 10 306 658 | 10 131 692 | 934 555 | 78 296 | 7,22 | y | n | n | n | n | n | y | Tiho |
| *macarthuri* | 05-08_hely | Anjiahely | M | -15.40000 | 49.48333 | se | ORG | 1 | HS2000 | 2 586 252 | 2 546 047 | 1 165 315 | 89 709 | 8,11 | y | n | n | n | y | y | y | Tiho |
| *macarthuri* | 07-08_hely | Anjiahely | M | -15.40000 | 49.48333 | se | ORG | 1 | HS2000 | 5 017 680 | 4 940 042 | 2 054 326 | 130 852 | 10,01 | y | **y** | n | n | y | y | y | Tiho |
| *macarthuri* | 01-13_hely | Anjiahely | M | -15.40000 | 49.48333 | se | ORG | 1 | HS2000 | 10 453 406 | 10 260 026 | 1 204 585 | 109 539 | 6,46 | y | n | n | n | n | n | y | Tiho |
| *macarthuri* | 04-13_hely | Anjiahely | F | -15.40000 | 49.48333 | se | ORG | 1 | HS2000 | 20 407 547 | 20 074 691 | 3 026 771 | 160 126 | 12,20 | y | **y** | **y** | **y** | n | n | y | Tiho |
| *mittermeieri* | PBZT115 | Anjanaharibe_Sud | un | -14.70692 | 49.54142 | pe | IDH | 3 | HS4000 | 18 911 748 | 15 880 074 | 6 573 187 | 194 266 | 6,10 | y | n | n | n | n | n | n | Omaha Zoo |
| *mittermeieri* | MBB014 | Anjanaharibe_Sud | un | -14.73483 | 49.49630 | pe | IDH | 3 | HS4000 | 2 432 618 | 2 150 664 | 857 599 | 63 670 | 3,41 | y | n | n | n | n | n | n | Duke Univ |
| *mittermeieri* | MBB013 | Anjanaharibe_Sud | un | -14.73483 | 49.49630 | pe | IDH | 3 | HS4000 | 10 048 050 | 9 167 206 | 2 297 644 | 103 237 | 7,37 | y | **y** | **y** | n | n | n | n | Duke Univ |
| *mittermeieri* | MBB016 | Anjanaharibe_Sud | un | -14.73483 | 49.49630 | pe | IDH | 3 | HS4000 | 9 798 820 | 8 952 200 | 2 292 249 | 114 259 | 7,04 | y | n | n | n | n | n | n | Duke Univ |
| *mittermeieri* | MBB012 | Anjanaharibe_Sud | un | -14.73483 | 49.49630 | pe | IDH | 3 | HS4000 | 7 979 794 | 7 256 250 | 2 133 437 | 95 885 | 7,36 | y | n | n | n | n | n | n | Duke Univ |
| *mittermeieri* | 2-07_hely | Anjiahely | M | -15.40000 | 49.48333 | se | ORG | 1 | HS2000 | 3 165 486 | 3 120 806 | 2 215 695 | 149 625 | 9,69 | y | n | n | n | y | n | n | Tiho |
| *mittermeieri* | 10-07_hely | Anjiahely | F | -15.40000 | 49.48333 | se | ORG | 1 | HS2000 | 4 764 527 | 4 600 941 | 3 050 452 | 175 709 | 11,41 | y | **y** | **y** | n | y | n | n | Tiho |
| *mittermeieri* | 04-07_hely | Anjiahely | F | -15.40000 | 49.48333 | se | ORG | 1 | HS2000 | 2 507 635 | 2 474 273 | 1 631 360 | 123 270 | 8,50 | y | n | n | n | y | n | n | Tiho |
| *mittermeieri* | 08-07_hely | Anjiahely | F | -15.40000 | 49.48333 | se | ORG | 1 | HS2000 | 3 122 027 | 3 081 386 | 2 160 238 | 146 723 | 9,68 | y | n | n | n | y | n | n | Tiho |
| *mittermeieri* | 07-06_habe | Antsahabe | M | -15.35000 | 49.40000 | se | ORG | 1 | HS2000 | 3 734 557 | 3 683 375 | 2 568 148 | 157 874 | 10,75 | y | **y** | n | n | y | y | y | Tiho |
| *mittermeieri* | RMR187 | Marojejy | un | -14.46722 | 49.83917 | pe | IDH | 3 | HS4000 | 10 442 144 | 9 665 430 | 2 939 717 | 121 575 | 8,28 | y | **y** | n | n | n | y | n | Duke Univ |
| *mittermeieri* | RMR186 | Marojejy | un | -14.46722 | 49.83917 | pe | IDH | 3 | HS4000 | 10 202 722 | 9 417 788 | 2 987 258 | 134 432 | 8,16 | y | n | n | n | n | y | n | Duke Univ |
| *mittermeieri* | MBB005 | Marojejy | un | -14.44183 | 49.82836 | pe | IDH | 3 | HS4000 | 4 592 378 | 4 196 120 | 1 265 583 | 85 303 | 4,60 | y | n | n | n | n | n | n | Duke Univ |
| *murinus* | RMR45 | Andranmena | un | -20.15000 | 44.55000 | pe | IDH | 3 | HS4000 | 7 887 466 | 7 237 190 | 2 654 886 | 144 551 | 7,44 | y | **y** | **y** | n | n | y | n | Duke Univ |
| *murinus* | RMR44 | Andranmena | un | -20.15000 | 44.55000 | pe | IDH | 3 | HS4000 | 9 692 978 | 8 480 102 | 3 429 888 | 144 995 | 9,05 | y | **y** | **y** | n | n | n | n | Duke Univ |
| *murinus* | RMR49 | Andranmena | un | -20.15000 | 44.55000 | pe | IDH | 3 | HS4000 | 13 879 770 | 12 657 850 | 3 619 021 | 154 257 | 9,76 | y | n | n | n | n | n | n | Duke Univ |
| *simmonsi* | A03_2014 | Ambodiriana | F | -16.67423 | 49.70128 | pe | TLS | 2 | HS3000 | 11 275 338 | 10 420 118 | 5 707 161 | 160 183 | 10,21 | y | **y** | **y** | n | n | n | n | IGC |
| *simmonsi* | A08_2014 | Ambodiriana | M | -16.67348 | 49.70018 | pe | TLS | 2 | HS3000 | 5 938 430 | 5 421 252 | 3 309 711 | 140 810 | 8,45 | y | **y** | n | n | n | n | n | IGC |
| *simmonsi* | BET87 | Betampona | un | -17.94086 | 49.20556 | pe | IDH | 3 | HS4000 | 2 076 776 | 1 740 114 | 769 320 | 55 865 | 2,13 | y | n | n | n | n | n | n | Omaha Zoo |
| *simmonsi* | PBZT117 | Betampona | un | -17.94081 | 49.20564 | pe | IDH | 3 | HS4000 | 1 304 936 | 1 090 922 | 485 010 | 27 770 | 1,74 | **n** | n | n | n | n | y | n | Omaha Zoo |
| *simmonsi* | POLO5.22 | Tampolo | un | -17.28839 | 49.40817 | pe | IDH | 3 | HS4000 | 8 415 742 | 7 087 654 | 3 112 336 | 145 968 | 4,56 | y | **y** | **y** | n | n | n | n | Omaha Zoo |
| *simmonsi* | ZAH2 | Zahamena | un | -17.48917 | 48.74722 | pe | IDH | 3 | HS4000 | 5 404 824 | 4 547 540 | 2 007 577 | 116 392 | 3,70 | y | **y** | n | n | n | n | n | Omaha Zoo |
| *simmonsi* | ZAH5 | Zahamena | un | -17.48917 | 48.74722 | pe | IDH | 3 | HS4000 | 5 060 014 | 4 266 274 | 1 891 031 | 104 351 | 3,66 | y | n | n | n | n | n | n | Omaha Zoo |

**Table S2: SNP datasets used across the analyses.**

#: number; sp: species; ind: individuals; GL: Genotype Likelihood; VCa: Variant Calling conducted with Angsd; VCg: Variant Calling conducted with GATK; Regional sampling includes *M. mittermeieri*, *M. lehilahytsara*, *M. macarthurii* and *M.* sp. #3.

| **Sampling** | **Process** | **Name** | **#sp** | **#ind** | **#GL** | **#SNP** |
| --- | --- | --- | --- | --- | --- | --- |
| All individuals | GL | GL1 | 6 | 63 | 1,096,639 |  |
| *M. maca-sp3* | GL | GL3 | 2 | 31 | 226,793 |  |
| All individuals | VCa | VCa1 | 6 | 63 |  | 28,738 |
| All pop. 22 ind. | VCa | VCa2 | 6 | 22 |  | 20,624 |
| All individuals | VCg | VCg | 6 | 52 |  | 61,519 |

**Table S3: *Microcebus* sp. #3 genome assembly statistics as calculated from SSPACE and BUSCO.**

The individual ID is MBB030 and was sampled along with other MBB *M.* sp. #3 individuals from Mananara-Nord NP (Tab. S1).

| **Assembly Statistic** | ***Microcebus* sp. #3** |
| --- | --- |
| Total Assembly Length (MB) | 2499.8 |
| Contig N50 (bp) | 36327 |
| Number of Contigs | 261207 |
| Scaffold N50 (bp) | 38968 |
| Number of Scaffolds | 243703 |
| Number of Gene Models | 42115 |
| % of Complete BUSCOS | 59.80% |
| % of Fragmented BUSCOS | 33.80% |
| % of Missing BUSCOS | 6.40% |
| **Annotation BUSCO Stats** |  |
| % of Complete BUSCOS | 62.10% |
| % of Fragmented BUSCOS | 25.00% |
| % of Missing BUSCOS | 12.90% |

**Table S5: Potential scale reduction factors for SNAPP analyses.**
The median potential scale reduction factors are calculated from posteriors of chain 1 and chain 2.

|  | **12-individual** | | **22-individual** | |
| --- | --- | --- | --- | --- |
| **Sampled Value** | **Median PSRF** | **Upper 97.5% PSRF** | **Median PSRF** | **Upper 97.5% PSRF** |
| posterior | 1.08 | 1.29 | 1.06 | 1.24 |
| likelihood | 1.00 | 1.00 | 1.00 | 1.00 |
| prior | 1.00 | 1.02 | 1.00 | 1.00 |
| theta0 | 1.06 | 1.25 | 1.00 | 1.02 |
| theta1 | 1.17 | 1.59 | 1.00 | 1.02 |
| theta2 | 1.07 | 1.25 | 1.06 | 1.22 |
| theta3 | 1.08 | 1.33 | 1.05 | 1.19 |
| theta4 | 1.38 | 2.18 | 1.04 | 1.08 |
| theta5 | 1.37 | 2.14 | 1.04 | 1.08 |
| theta6 | 1.08 | 1.30 | 1.06 | 1.19 |
| theta7 | 1.06 | 1.15 | 1.08 | 1.23 |
| theta8 | 1.10 | 1.38 | 1.12 | 1.42 |
| theta9 | 1.00 | 1.00 | 1.00 | 1.00 |
| theta10 | 1.02 | 1.08 | 1.00 | 1.00 |
| TreeHeightLogger | 1.07 | 1.25 | 1.06 | 1.22 |

**Table S6: Bayes factor support for the two sister lineage pairs.**
Marginal likelihoods were computed for a hypothesis of n speciation (Merge) and a hypothesis of a speciation event (Split). We tested the putative species pairs in both Clade I and Clade II with a 12- and 22-individual dataset. ^†^Bayes factors calculated as 2 * (lnL_Split_ - lnL_Merge_).

| **Species Pair** | **Number of Individuals** | **Merge Marginal lnL** | **Split Marginal lnL** | **2ln Bayes factor^†^** |
| --- | --- | --- | --- | --- |
| *M. macarthurii* -  *M.* sp*. #3* | 12 | -134,254 | -125,601 | 17,304 |
|  | 22 | -204,540 | -187,377 | 34,326 |
| *M. lehilahytsara* - *M. mittermeieri* | 12 | -126,515 | -125,601 | 1,828 |
|  | 22 | -187,873 | -187,377 | 993 |

**Table S7: Genetic divergence among species and sampling sites.**

*F*_st_ estimated among species (**first table**), and among sampling sites of each species (**following tables**) echoing the relationship revealed by phylogenetic approaches.

| *F*_ST_ | *M. lehilahytsara* | *M. macarthurii* | *M. mittermeieri* | *M. simmonsi* |  |  |  |
| --- | --- | --- | --- | --- | --- | --- | --- |
| *M. macarthuri* | 0.671 |  |  |  |  |  |  |
| *M. mittermeieri* | 0.077 | 0.657 |  |  |  |  |  |
| *M. simmonsi* | 0.506 | 0.786 | 0.491 |  |  |  |  |
| *M.* sp #3 | 0.657 | 0.416 | 0.663 | 0.728 |  |  |  |
| *F*_ST_ (*M. mittermeieri*) | Anjanaharibe_Sud | Anjiahely | Antsahabe |  | *F*_ST_ (*M.* sp. #3) | Ambavala | Antanambe |
| Anjiahely | 0.175 |  |  |  | Antanambe | 0.127 |  |
| Antsahabe | 0.120 | 0.122 |  |  | Mananara_nrd | 0.095 | 0.061 |
| Marojejy | 0.148 | 0.192 | 0.148 |  |  |  |  |
| *F*_ST_ (*M. simmonsi*) | Ambodiriana | Tampolo |  |  | *F*_ST_ (*M. lehilahytsara*) | Ambavala |  |
| Tampolo | 0.382 |  |  |  | Riamalandy | -0.019 |  |
| Zahamena | 0.398 | 0.190 |  |  |  |  |  |

### 5 Supplementary References

Adkins, R.M. and Honeycutt, R.L. 1994. Evolution of the primate cytochrome c oxidase subunit II gene. Journal of Molecular Evolution 38:215–231.

Ali, O.A., O’Rourke, S.M., Amish, S.J., Meek, M.H., Luikart, G., Jeffres, C., and Miller, M.R. 2016. RAD capture (rapture): Flexible and efficient sequence-based genotyping. Genetics 202:389–400.

Altschul, S.F., Gish, W., Miller, W., Myers, E.W., and Lipman, D.J. 1990. Basic local alignment search tool. Journal of Molecular Biology 215:403–410.

Blair, C., Heckman, K.L., Russell, A.L., and Yoder, A.D. 2014. Multilocus coalescent analyses reveal the demographic history and speciation patterns of mouse lemur sister species. BMC Evolutionary Biology 14:57.

Bolger, A. M., Lohse, M., and Usadel, B. 2014. Trimmomatic: a flexible trimmer for Illumina sequence data. *Bioinformatics***30**: 2114–2120.

Bouckaert, R., Heled, J., Kühnert, D., Vaughan, T., Wu, C.H., Xie, D., Suchard, M.A., Rambaut, A., and Drummond, A.J. 2014. Beast 2: a software platform for Bayesian evolutionary analysis. PLoS Computational Biology 10:e1003537.

Catchen, J., Hohenlohe, P. A., Bassham, S., Amores, A., and Cresko, W. A. 2013. Stacks: an analysis tool set for population genmics. *Mol. Ecol.* **22**: 3124–3140.

Drummond, A.J. and Bouckaert, R.R. 2015. Bayesian evolutionary analysis with BEAST. Cambridge University Press.

Etter, P.D., Preston, J.L., Bassham, S., Cresko, W.A., Johnson, E.A. (2011) Local de nvo assembly of RAD paired-end contigs using short sequencing reads. PLoS One 6:e18561.

Evann, G., Regnaut, G., and Goudet, J. (2005) Detecting the number of clusters of individuals using the software STRUCTURE: a simulation study. Molecular Ecology 14: 2611–2620.

Gelman, A., and Meng, X.L. (1998) Simulating nrmalizing constants: From importance sampling to bridge sampling to path sampling. Statistical Science 13:163-185.

Genmic Resources Development Consortium, Blanchet, S., Bouchez, O., Chapman, C.A., Etter, P.D., Goldberg, T.L., Johnson, E.A., Jones, J.H., Loot, G., Omeja, P.A., Rey, O., Ruiz-Lopez, M.J., Goudet, J. 2005. Hierfstat, a package for R to compute and test hierarchical f-statistics. Molecular Ecology ntes 5:184–186.

Gronau, I., Hubisz, M.J., Gulko, B., Danko, C.G., and Siepel, A. 2011. Bayesian inference of ancient human demography from individual genme sequences. Nature Genetics 43:1031–1034.

Guschanski, K., Olivieri, G., Funk, S., and Radespiel, U. 2007. MtDNA reveals strong genetic differentiation among geographically isolated populations of the golden brown mouse lemur, *Microcebus ravelobensis*. Conservation Genetics 8:809–821.

Haas, B.J., Papanicolaou, A., Yassour, M., Grabherr, M., Blood, P.D., Bowden, J., Couger, M.B., Eccles, D., Li, B., Lieber, M., MacManes, M.D., Ott, M., Orvis, J., Pochet, N., Strozzi, F., Weeks, N., Westerman, R., William, T., Dewey, C.N., Henschel, R., LeDuc, R.D., Friedman, N., and Regev, A. 2013. De nvo transcript sequence reconstruction from RNA-seq using the trinity platform for reference generation and analysis. Nature protocols 8:1494.

Irwin, D.M., Kocher, T.D., and Wilson, A.C. (1991) Evolution of the cytochrome b gene of mammals. Journal of Molecular Evolution 32:128-144.

Jombart, T. 2008. adegenet: a R package for the multivariate analysis of genetic markers. Bioinformatics 24:1403–1405.

Kahle, D. and Stamey, J. 2017. invgamma: The Inverse Gamma Distribution. URL https://CRAN.R-project.org/package=invgamma

Kearse, M., Moir, R., Wilson, A., Stones-Havas, S., Cheung, M., Sturrock, S., Buxton, S., Cooper, A., Markowitz, S., Duran, C., Thierer, T., Ashton, B., Meintjes, P., and Drummond, A. 2012. Geneious basic: An integrated and extendable desktop software platform for the organization and analysis of sequence data. Bioinformatics 28:1647–1649.

Korf, I. 2004. Gene finding in nvel genmes. BMC Bioinformatics 5:59.

Korneliussen, T.S., Albrechtsen, A., and Nielsen, R. 2014. ANGSD: analysis of next generation sequencing data. BMC Bioinformatics 15:356.

Larsen, P.A., Harris, R.A., Liu, Y., Murali, S.C., Campbell, C.R., Brown, A.D., Sullivan, B.A., Shelton, J., Brown, S.J., Raveendran, M., Dudchenko. O., Machol, I., Durand, N.C., Shamim, M.S., Lieberman Aiden, E., Muzny, D.M., Gibbs, R.A., Yoder, A.D., Rogers, J., and Worley, K.C. 2017. Hybrid *de nvo* genme assembly and centromere characterization of the gray mouse lemur (*Microcebus murinus*). BMC Biology 15:110.

Leaché, A.D., Banbury, B.L., Felsenstein, J., De Oca, A.N.M., and Stamatakis, A. 2015. Short tree, long tree, right tree, wrong tree: new acquisition bias corrections for inferring SNP phylogenies. Systematic Biology 64:1032–1047.

Lewis, P.O. 2001. A likelihood approach to estimating phylogeny from discrete morphological character data. Systematic Biology 50:913–925.

Li, H. 2013. Aligning sequence reads, clone sequences and assembly contigs with bwa-mem. arXiv preprint arXiv:1303.3997.

Li, H., Handsaker, B., Wysoker, A., Fennell, T., Ruan, J., Homer, N., Marth, G., Abecasis, G., and Durbin, R. 2009. The sequence alignment/map format and SAMtools. Bioinformatics 25:2078–2079.

Loecher, M. and Ropkins, K. 2015. RGoogleMaps and loa: Unleashing R graphics power on map tiles. Journal of Statistical Software 63.

Maniatis, T., Fritsch, E., and Sambrook, J. 1982. Molecular Cloning: a Laboratory Manual. New York: Cold.

Miller, A., Mills, H., Ralantoharijaona, T., Volasoa, N.A., Misandeau, C., Chikhi, L., Bencini, R., and Salmona, J. 2018. Forest type influences population densities of ncturnal lemurs in Manmpana, nrtheastern Madagascar. International Journal of Primatology 39:646–669.

Miller, A., Ralantoharijaona, T., Misandeau, C., Volasoa, N.A., Mills, H., Bencini, R., Chikhi, L., and Salmona, J. 2015. A biological survey of Antsahanadraitry forest (Manmpana) reveals the pressence of *Allocebus trichotis*. Lemur News 19:4–6.

Nadachowska‐Brzyska, K., Burri, R., Smeds, L., and Ellegren, H. (2016) PSMC analysis of effective population sizes in molecular ecology and its application to black-and-white *Ficedula* flycatchers. Molecular Ecology 25:1058-1072.

Nielsen, R., Korneliussen, T., Albrechtsen, A., Li, Y., and Wang, J. 2012. SNP calling, genotype calling, and sample allele frequency estimation from new-generation sequencing data. PLoS One 7:e37558.

O’Leary, S.J., Puritz, J.B., Willis, S.C., Hollenbeck, C.M., and Portny, D.S. 2018. These aren’t the loci you’re looking for: Principles of effective SNP filtering for molecular ecologists. Molecular Ecology 27:3193–3206.

Paradis, E., Claude, J., and Strimmer, K. 2004. APE: Analyses of phylogenetics and evolution in R language. Bioinformatics 20:289–290.

Pedersen, C.E.T., Albrechtsen, A., Etter, P.D., Johnson, E.A., Orlando, L., Chikhi, L., Siegismund, H.R., and Heller, R. 2018. A southern African origin and cryptic structure in the highly mobile plains zebra. Nature Ecology & Evolution 2:491.

Peng, X., Thierry-Mieg, J., Thierry-Mieg, D., Nishida, A., Pipes, L., Bozinski, M., Thomas, M.J., Kelly, S., Weiss, J.M., Raveendran, M., Gibbs, R.A., Rogers, J., Schroth, G.P., Katze, M.G., and Mason, C.E. 2014. Tissue-specific transcriptome sequencing analysis expands the nn-human primate reference transcriptome resource (nhprtr). Nucleic acids research 43:D737–D742.

Plummer, M., Best, N., Cowles, K., and Vines, K. 2006. Coda: convergence diagnsis and output analysis for MCMC. R News 6:7–11.

R Core Development Team. 2013. R: A Language and Environment for Statistical Computing. R Foundation for Statistical Computing, Vienna, Austria. URL: http://www.R-project.org/.

Radespiel, U., Olivieri, G., Rasolofoson, D.W., Rakotondratsimba, G., Rakotonirainy, O., Rasoloharijaona, S., Randrianambinina, B., Ratsimbazafy, J.H., Ratelolahy, F., Randriamboavonjy, T., Rasolofoharivelo, T., Craul, M., Rakotozafy, L., and Randrianarison, R.M. 2008. Exceptional diversity of mouse lemurs (*Microcebus* spp.) in the Makira region with the description of one new species. American Journal of Primatology 70:1033–1046.

Rambaut, A., Drummond, A.J., Xie, D., Baele, G., and Suchard, M.A. 2018. Posterior summarization in Bayesian phylogenetics using tracer 1.7. Systematic Biology 67:901–904.

Schüßler, D., Radespiel, U., Ratsimbazafy, J.H., and Mantilla-Contreras, J. 2018. Lemurs in a dying forest: Factors influencing lemur diversity and distribution in forest remnants of nrth-eastern Madagascar. Biological Conservation 228:17–26.

Seutin, G., White, B.N., and Boag, P.T. 1991. Preservation of avian blood and tissue samples for DNA analyses. Canadian Journal of Zoology 69:82–90.

Sgarlata, G.M., Salmona, J., Aleixo-Pais, I., Rakotonanahary, A., Sousa, A.P., Kun-Rodrigues, C., Ralantoharijaona, T., Jan, F., Zaranaina, R., Rasolondraibe, E., Zaonarivelo, J.R., Andriaholinirina, N.V., and Chikhi, L. 2018. Genetic differentiation and demographic history of the nrthern rufous mouse lemur (*Microcebus tavaratra*) across a fragmented landscape in nrthern Madagascar. International Journal of Primatology 39:65–89.

Simão, F.A., Waterhouse, R.M., Ioannidis, P., Kriventseva, E.V., and Zdobnv, E.M. 2015. Busco: Assessing genme assembly and anntation completeness with single-copy orthologs. Bioinformatics 31:3210–3212.

Stanke, M., Schöffmann, O., Morgenstern, B., and Waack, S. 2006. Gene prediction in eukaryotes with a generalized hidden markov model that uses hints from external sources. BMC Bioinformatics 7:62.

Swofford, D.L. 2003. Paup*: phylogenetic analysis using parsimony, version 4.0 b10.

Weir, B.S., and Cockerham, C.C. (1984). Estimating F-statistics for the analysis of population structure. Evolution, 38:1358-1370.

Xie W., Lewis P.O., Fan Y., Kuo L., Chen M.-H. 2011. Improving marginal likelihood estimation for Bayesian phylogenetic model selection. Syst. Biol. 60:150–160.

Yang, Z. and Rannala, B. 2010. Bayesian species delimitation using multilocus sequence data. Proceedings of the National Academy of Sciences 107:9264–9269.

Yoder, A.D., Campbell, C.R., Blanco, M.B., dos Reis, M., Ganzhorn, J.U., Goodman, S.M., Hunnicutt, K.E., Larsen, P.A., Kappeler, P.M., Rasoloarison, R.M., Ralison, J.M., Swofford, D.L., and Weisrock, D.W. 2016. Geogenetic patterns in mouse lemurs (genus *Microcebus*) reveal the ghosts of Madagascar’s forests past. Proceedings of the National Academy of Sciences 113:8049–8056.

Yu, G., Smith, D.K., Zhu, H., Guan, Y., and Lam, T.T.Y. 2017. ggtree: An R package for visualization and anntation of phylogenetic trees with their covariates and other associated data. Methods in Ecology and Evolution 8:28–36.
